# Predictive uncertainty in mechanistic models of cellular processes calibrated to experimental data

**DOI:** 10.1101/2021.05.18.444743

**Authors:** Michael W. Irvin, Arvind Ramanathan, Carlos F. Lopez

**Affiliations:** Department of Biochemistry, Vanderbilt University, Nashville, TN 37232; Data Science and Learning Division, Argonne National Laboratory, Lemont, IL 60439; Department of Biomedical Informatics, Vanderbilt University Medical Center, Nashville TN 37232

## Abstract

Mathematical models are often used to explore network-driven cellular processes from a systems perspective. However, a dearth of quantitative data suitable for model calibration leads to models with parameter unidentifiability and questionable predictive power. Here we introduce a Bayesian and Machine-Learning based Measurement Model approach to explore how quantitative and non-quantitative data constrain models of apoptosis execution within a missing data context. We find two orders of magnitude more ordinal (e.g. immunoblot) data are necessary to achieve accuracy comparable to quantitative (e.g. fluorescence) data. Notably, ordinal and nominal (e.g. immunostain) non-quantitative data synergize to reduce model uncertainty and improve accuracy. Further, model prediction accuracy and certainty strongly depend on rigorous data-driven formulations of the measurement, and the size and make-up of the datasets. Finally, we demonstrate the potential of a data-driven Measurement Model approach to identify model features that could lead to informative experimental measurements and improve model predictive power.

## Introduction

The combination of systems approaches and quantitative data, has promised a novel understanding of cellular mechanisms that could spur science-driven innovation in biology and medicine – as has happened in physics, chemistry, and engineering^1-3^. However, despite massive research efforts and data accumulation, our understanding of cellular regulation, signaling and many other processes as biomolecular *systems* remains rudimentary. The quantitative and systems biology fields continue to employ strategies from physics and engineering to construct models of biological mechanism from first principles^4,5^. Yet these strategies are incompatible with the types of measurements and observations that predominate biological investigations. Observations from biological experiments investigating cell fate outcomes (apoptosis, necroptosis, etc.) are collected as cellular phenotype (or fractions of cellular phenotype) and other categorical observations, which are hard to define in terms of variables encoded in mathematical mechanistic models of biological processes^6^. Therefore, the connection of mechanistic models to corresponding biological measurements is subject to practitioner interpretation. As a result, vast amounts of existing nonquantitative data in cell biology have led to mechanistic formulations based on simple inference and informal reasoning. In addition, the noise, complexity and hierarchical organization of organisms limits how one can experimentally perturb and measure biological systems^7,8^. Therefore, a relative dearth of quantitative data exists that reveals itself in mechanistic models with poorly constrained parameters and unreliable predictions. Unfortunately, both non-quantitative and quantitative data, collected in an unplanned manner, results in missed opportunities to quantitatively explain complex cellular mechanisms^9^.

This data-to-knowledge problem in biology has prompted researchers to incorporate nonquantitative data as a complement or substitute for quantitative data in the development of mechanistic models^10-13^. The traditional workflow employed to train mechanistic models to data comprises mechanistic models and experimental measurements linked through a calibration method (Box 1)^14, 15^. Such workflows have been adapted to incorporate nonquantitative data into mechanistic models and have revealed their intrinsic value in mechanistic hypothesis exploration. For example, pioneering work by Pargett and co-workers employed optimal scaling and multi-objective optimization for training mechanistic models to large ordinal datasets^10^. Schmiester et al. incorporated this strategy into PyPESTO, a model parameter estimation framework^11^. Their formulation imposes discrete boundaries on the mechanistic model to reflect discrete ordinal values in the data, but this approach limits their ability to integrate multiple datatypes or use Bayesian methods for training and uncertainty estimation of mechanistic models. More recently, Mitra et al. applied predefined constraint-based models of categorical data and modified their approach to allow definition of a likelihood function within a Bayesian formalism.^12, 13^ However, the *ad hoc* nature of their constraint models leaves room for biasing assumptions. Schmiester et al. introduces models of fractional cell fate that, while amenable to Bayesian methods, requires *ad hoc* speculation about which features describe cell fate and how those features map to values of fractional cell fate^16^. Given the limited application of Bayesian methods and biases introduced by *ad hoc* assumptions, the field still has a limited understanding of the contribution of nonquantitative and quantitative data to mechanistic knowledge in biological systems.

### Box 1: Objective functions and the role of a measurement model.

Mechanistic models of biological processes are typically encoded as systems of (ordinary) differential equations (Eq. 5). Model calibration relies on an objective function (Eq. 7) -- or in a Bayesian setting, a likelihood function (Eq. 8) -- quantifies the degree of dissimilarity or similarity between model variables and corresponding measurements. Note, the objective or likelihood function uses measurement model (Eq. 6) which converts modeled variables ***x***(*t*) to a quantity *y*(*t*_*i*_, ***θ***) that can be compared to data *ŷ*(*t*_*i*_). In physics and engineering, where measurements are typically quantitative, the measurement model can be neglected. For nonquantitative measurements and observations, the measurement model takes more consideration.

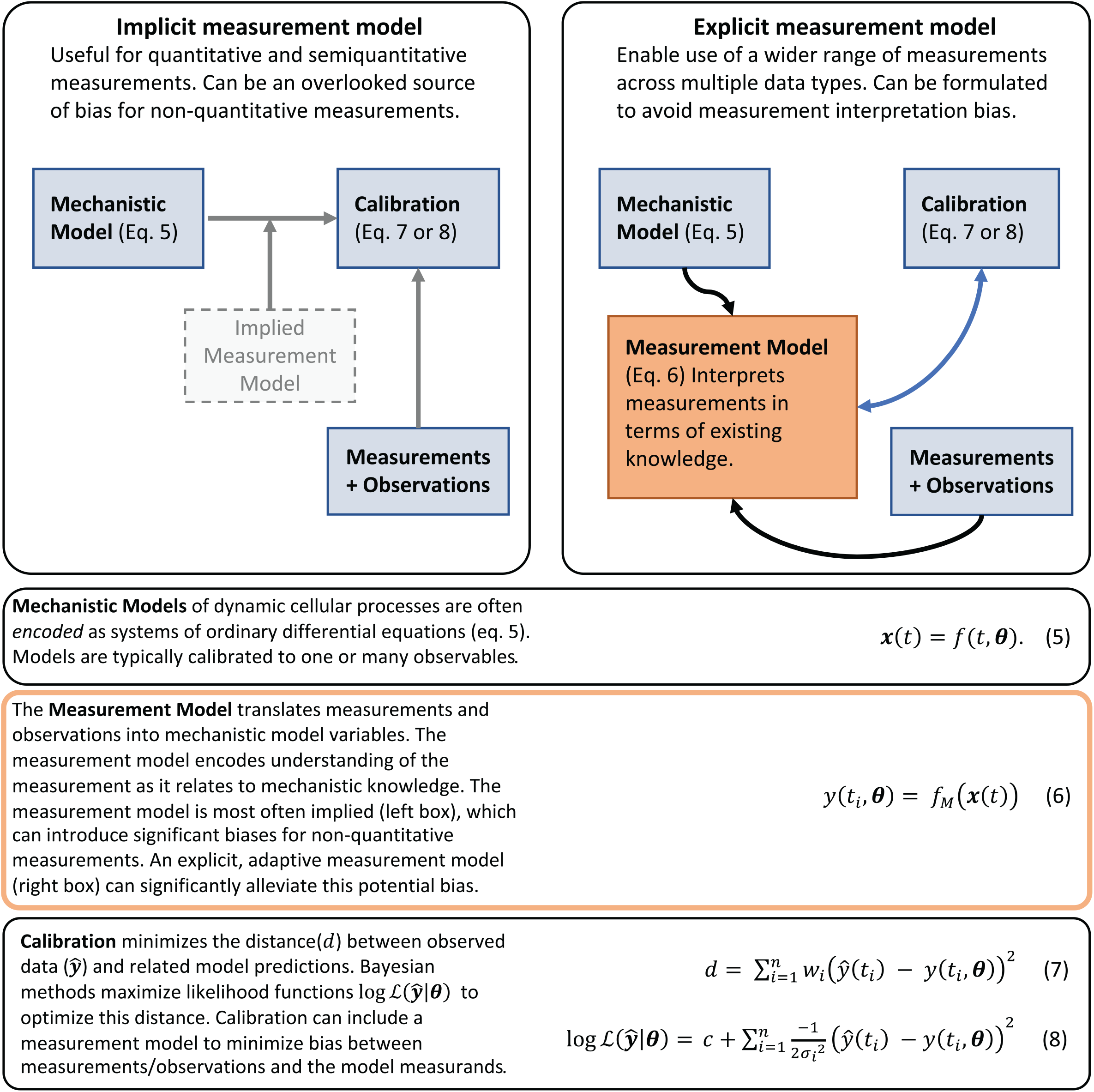

In this work, we tackle the data-to-knowledge challenge by introducing the concept of a *measurement model* -- machine learning construct -- into systems modeling approaches, which aims to rigorously define measurements and observations in terms of an underlying mechanism^17^. This definition entails formulation of a function that maps variables encoded in a mechanistic model to values in the observed data. Our approach departs from previous work in that it uses machine learning based classification and regression models whose free parameters are estimated to accomplish data-driven identification of measurement model properties. It also uses a probabilistic formulation that lends itself to Bayesian methods and can therefore provide an unbiased evaluation of the predictive power of models trained to nonquantitative data. In what follows, we present our findings about common types of biological measurements, followed by a presentation of our methodology. In this work we use a mechanistic model of apoptosis execution to demonstrate how the amount, and type, of data applied to a mechanistic model can affect its predictive power. It is well established that apoptosis signaling is involved in many cellular processes in health, disease, and development^18^. Its biological importance is further underscored by available quantitative and nonquantitative empirical data^19^. We also establish how an *ad hoc* formulation of a measurement model can lead to spurious results and further show how these *a priori* assumptions can be examined within a Bayesian, data-driven context. Further, we demonstrate the potential of a machine learning measurement model formulation to identify phenomenological links between features (e.g., predictors and drivers) of a biomolecular mechanism and emergent biological phenotype. Finally, we apply the measurement model concept to calibrate our mechanistic model of cell death to published fractional cell death data. In this example, our measurement model avoids *ad hoc* speculation about the how the mechanistic model variables map to values of fractional cell fate by instead using Gaussian process regression to learn which variables (and functions) map onto the measured fractional cell death values. The mechanistic model calibration, using this approach, produced accurate predictions underlying cell death dynamics and identified, in the cell death dynamics, predictive features of cell fate that are consistent with published observations. We expect our approach to improve our understanding of the data-to-knowledge relationship in biological processes, leading to a probabilistic understanding of biochemical mechanisms, and accelerated identification of systems-level interactions that drive biological network dynamics.

## Results

### Contributions and biases from different data types to mechanistic models

We first explored how experimental data measurements are used to constrain mathematical models of cellular processes. Mechanistic models typically employ physical chemistry formalisms comprised of reaction rates and chemical species concentrations to represent networks of biochemical reactions. Direct quantitative measurement of all chemical reactions and species would provide needed model parameters to carry out simulations and *in silico* experiments. However, these measurements are typically not available and likely untenable for real systems, thus leading to indirect measurements used to infer model parameter values using an objective function (Eq. 7) or a likelihood function (Eq. 8). When these functions are optimized, the resulting mathematical model can provide valuable new predictions and insights about the cellular process. Measurements from cell biology experiments comprise four broad types, namely, nominal, ordinal, semiquantitative (indirect and direct), and quantitative (Figure 1); each data type reveals different insights about the cellular process. In apoptosis signaling, for instance, nominal observations supported early research where it helped identify key components in the apoptosis signaling pathway^20^. Apoptosis and survival outcomes – as indicated by nominal nuclear fragmentation data (Figure 1 *top row*) – helped determine two parallel signaling arcs that proceed following initiator caspase activation: mitochondria-dependent and -independent pathways^20^. These pathways trigger apoptosis by activating effector caspases^20^. We built an abridged Extrinsic Apoptosis Reaction Model (aEARM)^21^, which represents these extrinsic apoptosis execution mechanisms as biomolecular reactions (Figure 2A). Nominal observations do not provide a definitive estimation of their quantity of interest (i.e., their *measurand*) and instead, encode weak constraints on the measurand values (Eq. 1). They can guide mechanistic modeling by revealing salient structural elements of a cellular process but provide limited insight into the dynamics and complex regulatory cues of apoptosis signaling. Ordinal measurements have featured prominently in works investigating apoptosis signaling. They have uncovered clues about the dynamics and complex regulatory mechanisms of apoptosis. For instance, ordinal measurements of DISC (i.e., a ligand-dependent membrane bound ‘death inducing signaling complex’) components, initiator- and effector-caspases (Figure 1 *second row*), Bcl-2 family proteins (e.g., Bid), etc. revealed how cells resist apoptosis by limiting (but not totally eliminating) pro-apoptotic cues^22^; the sub-maximal pro-apoptotic signaling presents as delay in the dynamics of caspase activation^23^. To better understand caspase activation dynamics and its effect on apoptosis and survival, we need mathematical models of the apoptosis signaling dynamics. Ordinal measurements, however, do not readily support a mathematical description of apoptosis signaling dynamics. Emerging work has leveraged ordinal and nominal measurements in the development of mathematical models of biological signaling but the weak constraints encoded by these measurements (Eq. 1 and Eq. 2) add uncertainty and bias to the modeling process. Indirect semi-quantitative (e.g., fractional cell fate) measurements report quantitative values but lack a definitive mapping between those values and the measurand. While these measurements can encode tighter constraints on the measurand (Eq. 3), but the indirectness of the measurement provides little, or no, information about the function *f*_*M*_ that could describe these constraints. Indirect semi-quantitative measurements therefore occur in the early investigative stages and offer similar insights as nominal and ordinal measurements.

**Figure 1:**
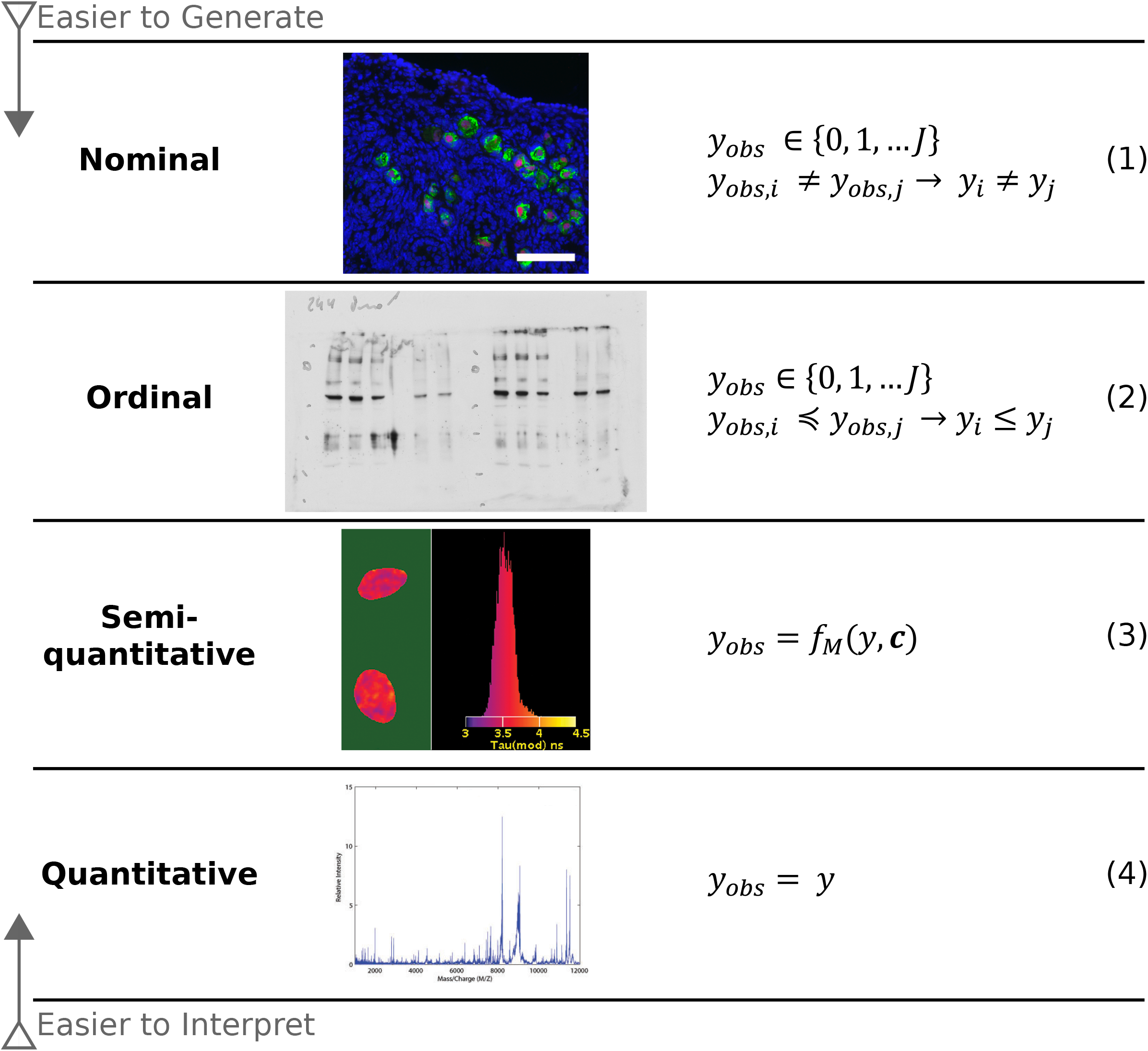
Measurements encountered in cell biology. *Nominal* measurements (top) can help understand intracellular signaling activity as it relates to broader cellular and physiological behaviors. With cellular phenotype markers or drivers, we can attribute different nominal observations to distinct (intra)cellular states. This is often modeled as in Eq. 1, where each observable measurement (*y*_*obs*_) corresponds to a given state. *Ordinal* measurements (second row) can be graded cellular phenotype observations (e.g., cell state transitions in cellular differentiation) or measurements of intracellular contents where noise can obscure intervals between values (e.g., Western Blots). Ordinal measurements imply a relative ordering of quantities along an axis but not their relative distance; i.e. we may know *y*_*i*_ ≤ *y*_*j*_ without knowing *y*_*i*_ − *y*_*j*_ (Eq. 2). Semi-quantitative measurements (third row) typically arise when an investigation has progress toward a more quantitative understanding of the intracellular signaling. Cell fate is a nominal observation while fractional cell fate measurements are semi-quantitative. We considere fractional cell fate measurements as *Indirect Semi-Quantitative* as they are indirect process measurements that lack a clear connection between mechanistic model and data. This presents as a less certainty formulation of *f*_*M*_. The contribution of indirect semi-quantitative measurements therefore is comparable to that of nominal and ordinal measurements. Direct semi-quantitative measurements (e.g., fluorescent intracellular markers) imply a quantitative relationship but a scaling function is necessary for true quantitation (Eq. 3). True quantitative measurements (bottom row) do not imply assumptions and the quantity measured can be used directly in the model (Eq. 4), such as mass-spectrometry protein concentration measurements. As shown schematically on the left triangle schematic, ordinal and nominal measurements are more abundant in biology due to their ease of production but are more difficult to interpret, whereas semiquantitative and quantitative measurements are less common but have a more straightforward interpretation.

**Figure 2.**
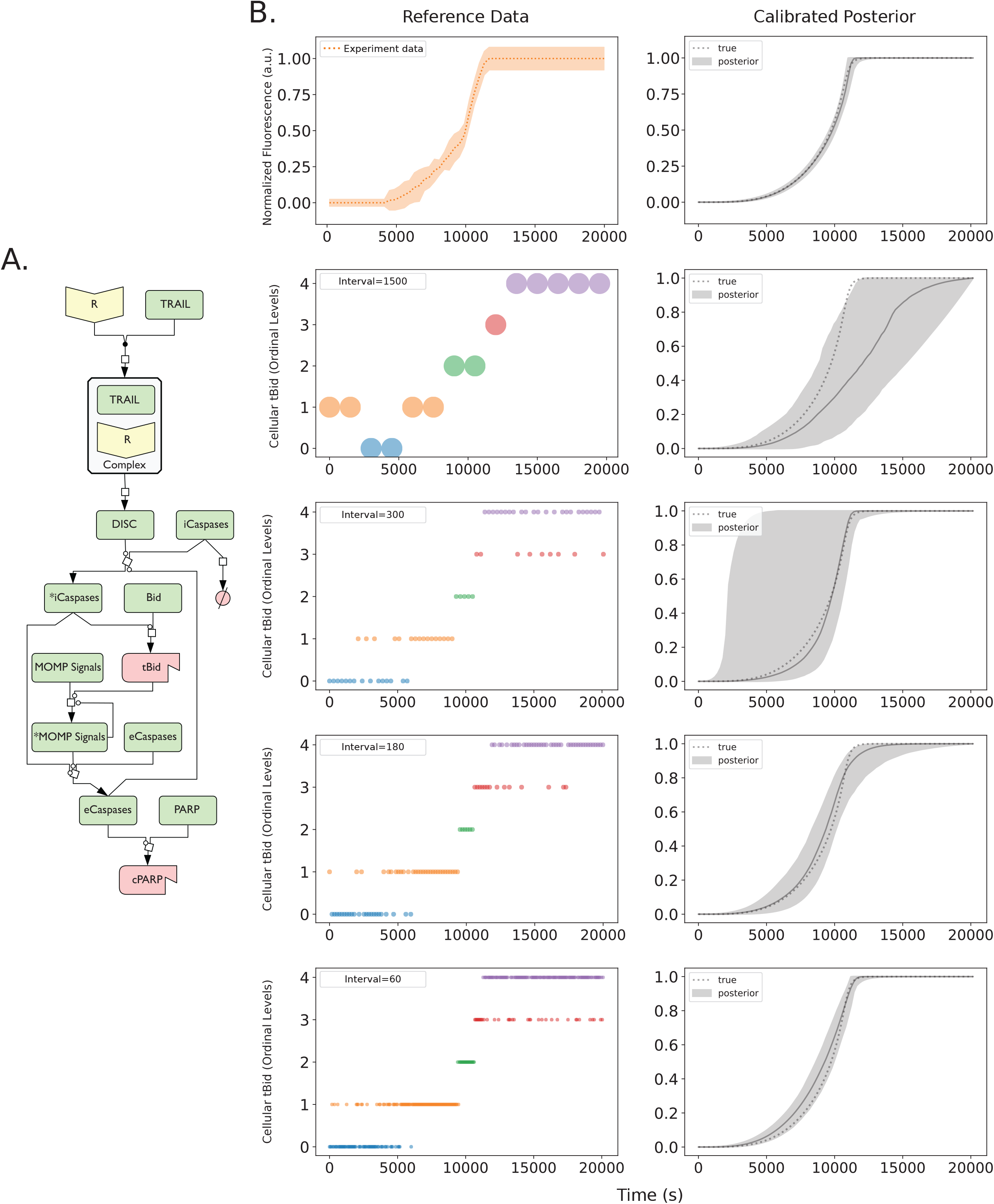
Predicted Bid truncation dynamics of aEARM trained to different sized ordinal datasets. Multiple Bayesian optimizations were run on the A.) abridged Extrinsic Apoptosis Reaction Model (aEARM) using different sized ordinal dataset to probe how dataset size influenced certainty of aEARM predictions. B.) Initiator caspase reporter (IC-RP) fluorescence time-course measurements (at 180s intervals) were measured (top left) as a proxy for truncated tBid (data from Albeck et al^29^). The plot shows the mean (dotted line) +/- 1 standard deviation (shaded region) for each time point. The 95% credible region (top right) of posterior predictions (shaded region) for tBID concentration in aEARM, calibrated to fluorescence measurements of IC-RP and EC-RP (See also supplemental figure 3). The median prediction (solid-line) and true (dotted line) tBID concentration trajectories are shown. In the next four rows (from top to bottom), Ordinal measurements of tBID (left) at every 1500, 300, 180 and 60s interval, respectively. The 95% credible region of predictions (shaded region), median prediction (solid line) and true (dotted line) tBID dynamics for aEARM calibrated to ordinal measurements of tBID and cPARP occurring at every 1500, 300, 180 and 60s timepoint are plotted in plots on the right. The plots for cPARP ordinal measurements and predictions are found in Supplemental Figure 3.

Technical challenges confine our quantitative and direct semi-quantitative measurement to just a few apoptotic signaling proteins. Fluorescence indicators of caspase activity^19^ (and by proxy, caspase substrate cleavage) enabled time course measurements of Bid and PARP cleavage dynamics (Figure 2B *top row*)^18^. They revealed pro-apoptotic activation of Bid and PARP, in TRAIL induced apoptotic HeLa cells, follows sigmoidal dynamics with delays and switch times that are sensitive to various regulatory factors. These measurements provide the details necessary for a mathematical description of apoptosis signaling dynamics and complexity. Our mathematical model aEARM captures the events from initial death ligand cue, initiator caspase activation, BID truncation (tBID), mitochondrial outer membrane permeabilization (MOMP) and eventual PARP cleavage (cPARP), as shown schematically on Figure 2^24-28^. The model was calibrated to above fluorescence data, as described in *Methods*^29^. Direct semi-quantitative measurements like fluorescence, though more defined than indirect semi-quantitative measurements, still lack a definitive estimation of the measurand because their interpretation requires mathematical manipulation, typically through a scaling function (Eq. 3), which can also add uncertainty and bias. Quantitative measurements can be used directly in a model without further modifications (Figure 1, *fourth row*) thus minimizing the uncertainty and bias introduced in the model from measurement interpretation. Therefore, the specific type of measurement and its interpretation could add significant uncertainty and bias to the mechanistic explanation of a given process.

To study the bias and uncertainty originating from different types of measurements, we introduce a concept from statistics, and social sciences: the *measurement model* (Box 1)^30^. Briefly, a measurement model is a function (Eq. 6) that describes the relationship between the measurement and its measurand. This function maps variables from the mechanistic model ***x*** to the values expressed in the data *ŷ*. This function is often assumed or implied, particularly for direct semi-quantitative data that can more readily be applied to the model calibration. However, the application of nominal, ordinal and indirect measurement types to mechanistic models is not straightforward, because their interpretation (as we show in the following sections) can significantly bias model-derived insights. Consequently, modeling efforts have relied almost exclusively on quantitative and direct semi-quantitative data. By contrast, the much more abundant non-quantitative datatypes are often ignored or used inappropriately.

Early modeling efforts interpreted nonquantitative data as a series of arbitrary surrogate quantities for the ordinal or nominal values in a corresponding dataset^14^. More recently, discrete boundaries on the values of the measurand were imposed along with a distance metric to describe how well the mechanistic model satisfies nominal or ordinal constraints in the non-quantitative data^10-13^. These approaches reveal the value of nonquantitative data for mechanistic model calibration, but the often-ad hoc nature of these constraint-based measurement models has been an overlooked source of model bias. Similarly, models of indirect semi-quantitative measurements use *ad hoc* functions whose formulation introduces model biases. To minimize biases from the interpretation of non-quantitative datatypes and apply Bayesian inference methods for model calibration, we developed a data-driven probabilistic measurement model (Box 2). Our measurement model is data-driven in that it possesses free parameters that are calibrated to match data; this lets us replace *a priori* assumptions about the measurement with a data-driven parametrization, and thereby calibrate mechanistic models whose accuracy and precision better reflect the information contained in the data. Our measurement model is probabilistic as it replaces discrete boundary-based measurement models and distance metrics with a *probability* (Box 2, Eq. 9) of the ordinal or nominal value, which enables easy formulation of a likelihood function and application of Bayesian optimization methods that utilize MCMC sampling. In our approach, the measurement model is a mathematical construct that represents the measurand through machine learning probabilistic classification and regression models, whose free parameters are estimated simultaneously with the free parameters of the mechanistic model during calibration (Box 2). As a probabilistic model, the measurement model effectively describes the probability of the categories encoded in the non-quantitative data given values of the measurand (Eq. 9). The measurand, in our case, is encoded in the mechanistic model. For example, the measurement model (Eq. 9, Box 2) can use ordinal logistic classifiers to model the probability of a categorical value as a function of variable(s) encoded by the mechanistic model. Similarly, logistic classifiers can model the probability of cell death or survival observations given specific states of the mechanistic model. We also considered a probabilistic regression model of indirect semi-quantitative data (e.g., fractional cell fate data). The corresponding likelihood function for that measurement model is detailed in the methods section. In the calibration process, the measurement model is an explicit intermediate step between simulation of the mechanistic model dynamics and calculation of the likelihood (Box 2). As described in the *Methods* section, this approach uses the Python based PySB models-as-programs framework and PyDREAM, a Python implementation of the DREAM_(ZS)_ algorithm to sample posterior values of models’ free parameters. However, other model-building and parameter sampling (or optimization) algorithms could be employed by the user. In what follows, we examine the impact of different measurement modalities and interpretations on mechanistic model constraints in apoptosis execution. This work motivates an approach that could be generalized to any mathematical model to rigorously integrate quantitative and nonquantitative data types.

#### Box 2: Model calibration with the data-driven probabilistic measurement model.

A.) The measurement model is an intermediate step between the mechanistic model and likelihood function of the measurement/observations. It receives variables from the mechanistic model and transforms for use in the likelihood function. This probabilistic machine-learning measurement model estimates *probabilities* of class membership as a function of the mechanistic model variables (Eq. 9). This measurement model is data-driven in that it contains free-parameters that are evaluated via the likelihood function (Eq. 10). B.) The measurement model uses values of e.g. tBID (grey curve) to estimate the probability of membership in an ordinal category (dotted data). C.) Plots a posterior ensemble of estimates of the probability of membership into 5 ordinal categories (x-axis) as a function of normalized tBID concentration (y-axis). The plot shows the median (solid line) and 95% credible region (shaded region) of the predictions (Category colors match data plotted in B). **Algorithm)** The mechanistic model and measurement model are calibrated simultaneously using Bayesian sampling methods through stepwise operations as described in each numeral.

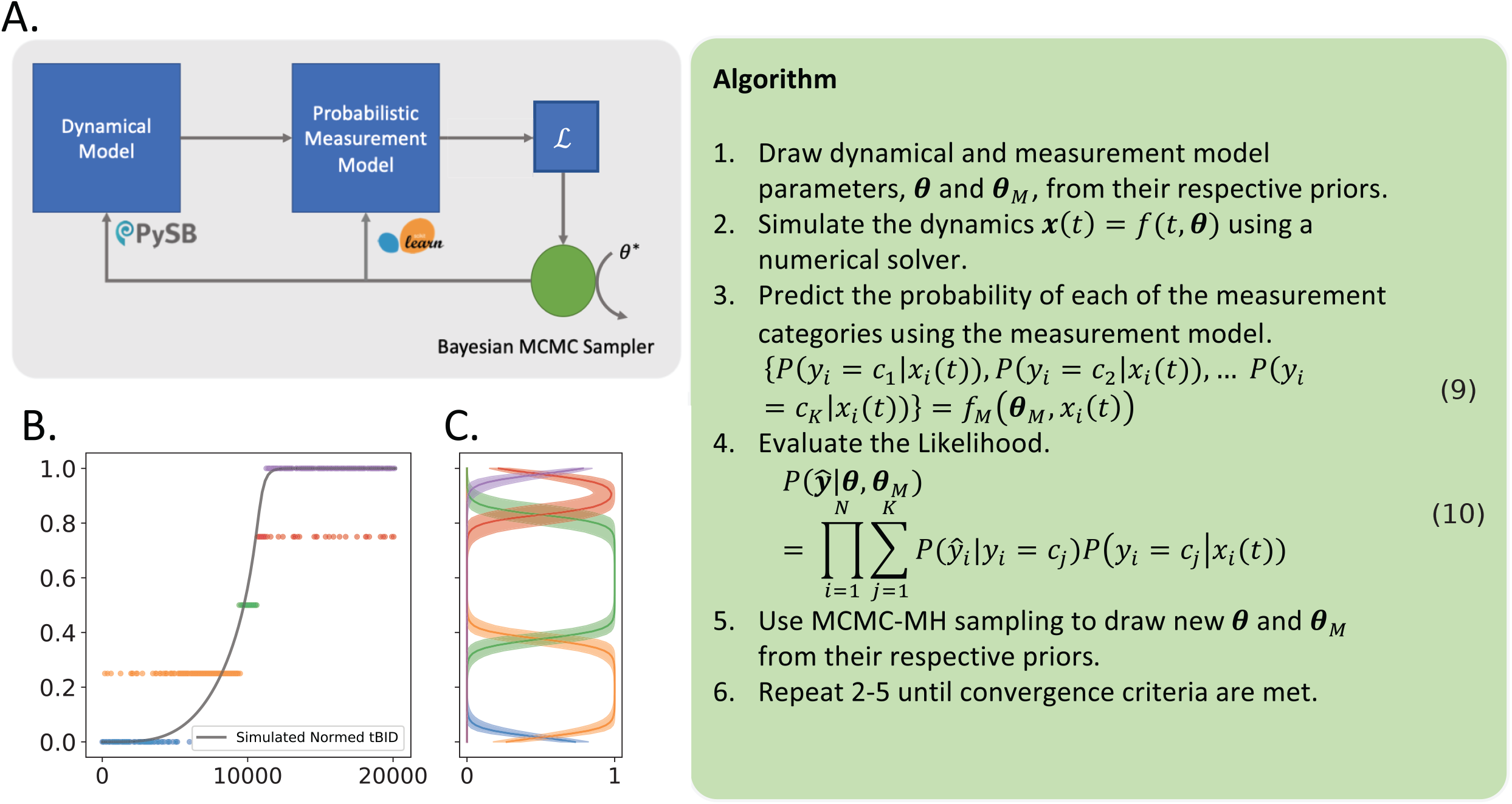

### Uncertainty associated with different data types in model calibration

To date, molecular biology investigations of intracellular signaling processes and their mechanisms predominantly report nonquantitative measurements. However, it is unclear exactly how well these measurements support the development of mechanistic models. We therefore asked how various measurement datatypes impact the certainty and accuracy of model calibrations. Specifically, we explored how to adjust the size and make-up of nonquantitative datasets to better support mechanistic inferences. The resulting posterior predictive region for tBID dynamics of aEARM calibrated to (direct semi-quantitative) fluorescence data is shown in Figure 2 (B, top row). As expected, the data can effectively constrain the model and the 95% credible region of posterior predictions for tBid dynamics falls within the data uncertainty region. We then extracted a parameter vector from the fluorescence optimized data and used it as a baseline (reference) to generate ordinal datasets for tBID and other aEARM variables as described in *Methods* and shown Figures 2 (B, bottom four rows). These synthetic datasets could be considered as numerical representations of a time-course western blot dataset. We then calibrated aEARM kinetic rate and measurement model parameters to the ordinal and nominal datasets.

As shown in Figure 2B, ordinal datasets accurately predicted quantitative predictions of “ground truth” dynamics for tBID. The 95% credible region of posterior predictions of tBID dynamics of aEARM trained to these ordinal datasets each contained “ground truth” dynamics for tBID. We also use the area bounded by the 95% credible region of posterior predictions of tBID as a measure of model certainty; with a smaller area indicating higher certainty. The ordinal dataset containing measurements at every 25-minute interval (i.e., typical of time-dependent western blot datasets), however, did not significantly constrain the posterior predictive regions of these dynamics (Figure 2D). Increasing the number of measurements, however, increases the certainty of the posterior predictions of tBID dynamics; this certainty approaches that of the typical semi-quantitative (fluorescence) dataset, which has an area of 2.7, when then the number of ordinal measurements is increased threefold, which had an area of 6.2. The areas bounded by the 95% credible region for each ordinal time-course dataset is described in the Figure 2B (Bottom two rows).

To explore the impact of nominal data on model optimization, we again extracted a parameter vector from the fluorescence optimized data and used it as a baseline (reference) to generate nominal datasets akin to an apoptosis execution observation as described in Methods. Previous work has described how features of apoptosis signaling dynamics can predict cell death vs survival^19^. The generated nominal dataset describes binary cell-fate outcomes that emerge because of extrinsic apoptosis signaling dynamics. We encode this information in a nominal measurement model as described in *Methods*. Parameters of aEARM and the free-parameters encoded in the measurement model were jointly calibrated to a synthetically generated dataset of 400 survival vs death outcomes as shown in Figure 3A (left). As shown in Figure 3A (right), the binary cell-fate data minimally constrain the posterior predictive region of tBID dynamics relative to the prior constraints on the model. This is expected as the binary cell-fate data-type essentially condenses complex apoptotic signaling dynamics to a single categorical value. *Supplemental Fig 20* shows the 95% credible region of posterior predictions of normalized tBID dynamics of aEARM trained to published fractional cell death data. These data differ from synthetic ordinal and nominal datasets in that they lack a known reference to “ground truth” dynamics and should therefore not be compared to the plots in Figures 2 and 3.

**Figure 3.**
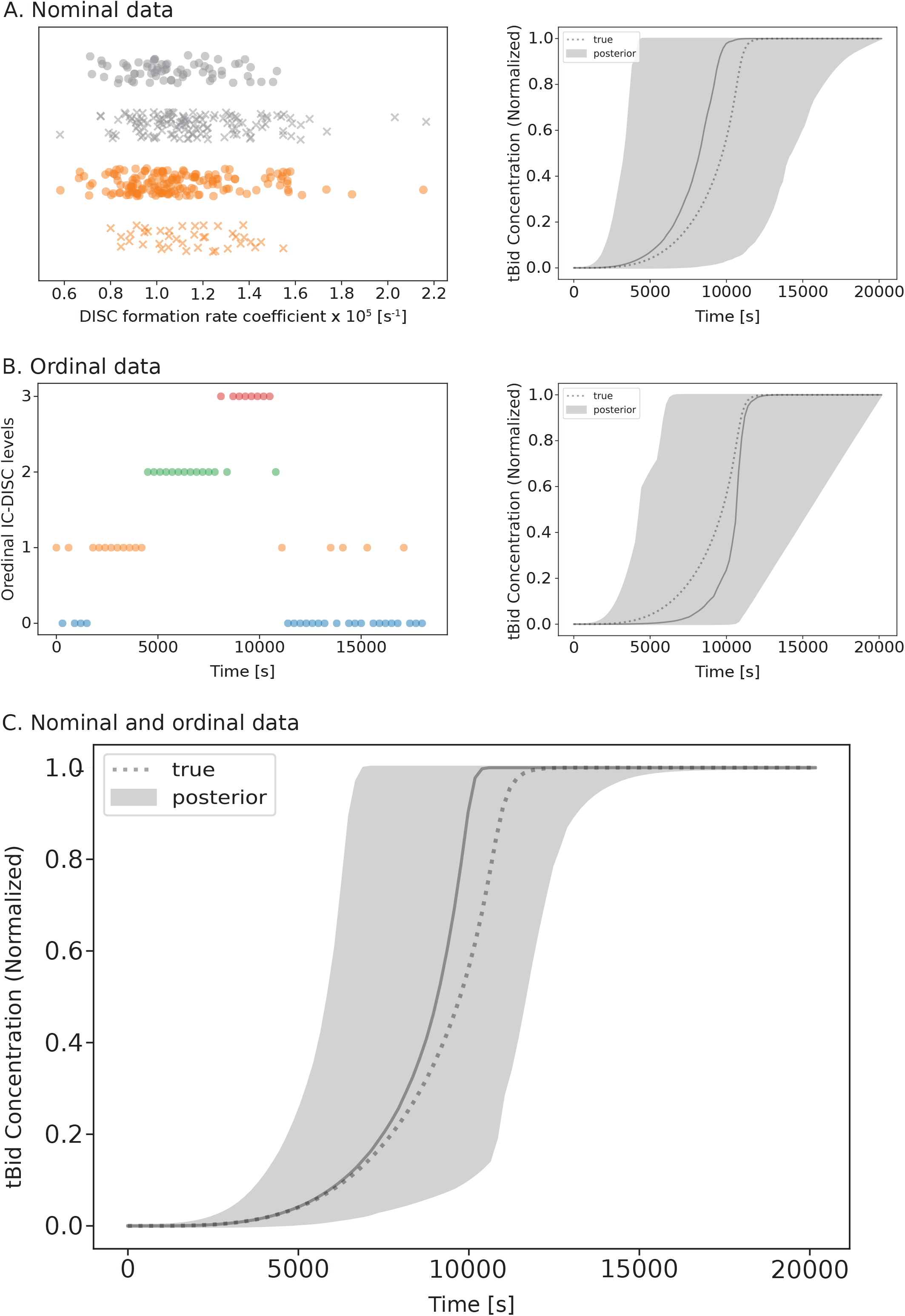
Predicted Bid truncation dynamics of aEARM trained to nominal and ordinal datasets. A.) Nominal cell death (x) vs survival (o) outcomes data for cells treated with 10ng/mL (orange) and 50ng/mL (grey) of TRAIL and with known relative values of DISC formation (x-axis). The 95% credible region (shaded region) of posterior predictions of tBID dynamics of aEARM calibrated to nominal data (right plot). The median prediction (solid-line) and true (dotted line) are also plotted. B.) Ordinal measurements for initiator caspase-DISC colocalization (IC-DISC) at 300s intervals (left plot). The 95% credible region (shaded region) of posterior predictions of tBID dynamics of aEARM calibrated to ordinal IC-DISC data (right plot), and C.) of aEARM calibrated to nominal *and* ordinal IC-DISC data. The median prediction (solid-line) and true (dotted line) were also plotted. The fit to IC-DISC data are shown in Supplemental Figure 9.

In lieu of its limited ability to constrain mechanistic models, modeling efforts understandably disregard nominal data. However, we hypothesized that combining nonquantitative datatypes and covering multiple variables in the model could improve model certainty. To explore the effect of multiple data type combinations on model calibration, we again optimized the aEARM model parameters, but this time to a dataset containing nominal and ordinal measurements. As described in *Methods*, we added a synthetic dataset containing 61 ordinal time-course measurements for the DISC complex to the nominal dataset described above (Figure 3B (left)). We modeled the likelihood of this combined dataset as the product of the likelihoods of the individual constituent datasets (see *Methods* for details). In Figure 3A and 3B (right), we see the nominal and ordinal datasets yields larger 95% credible regions for the posterior predictions of tBID dynamics. However, (in Figure 3C) the combined dataset better constrained the posterior predictions of normalized tBID dynamics than either dataset alone, with a 95% credible region area of 26.5 (compared to 55.0 and 56.4 for the ordinal and nominal datasets alone). Therefore, the model uncertainty stemming from only using tBID nominal data was decreased by including more detailed upstream measurements. However, the contribution of DISC ordinal data alone was comparable to that of the tBID nominal data in isolation (Figure 3B (right)). This data suggests that distributed measurements across multiple variables in a pathway yield synergistic effects on calibrated model accuracy and certainty.

### Data-driven measurement model as an indicator of model bias

Traditionally, applying quantitative or semi-quantitative data to a mechanistic model has been relatively straightforward as they typically follow a well-establish and simple relationship between the measurement and the measurand. However, for non-quantitative data, measurement uncertainty can prompt researchers to make assumptions about the relationship between measurement and measurand, which may negatively impact in the resulting mechanistic model. We therefore asked how the encoding of assumptions into our models of non-quantitative measurements could impact mechanistic model calibrations. To attain this goal, we calibrated aEARM kinetic rate parameters to the ordinal dataset, but this time we replaced the free parameters in the measurement model fixed *a priori* parameterizations or we encoded our assumptions as priors on the measurement model’s free-parameters. We tested four situations: (*i*) fixed parameters, a case where the measurement model is pre-parameterized by the user, presumably reflecting full confidence in their assumptions about the measurement; (*ii*) strong prior knowledge, a case where there is strong belief in the assumed values of the measurement model parameters; (*iii*) weak prior knowledge, a case where there is only weak belief in the assumed values of the measurement model parameters; and (*iv*) no prior knowledge, that is no constraints on the measurement model parameters.

Figures 4A and 4B show the ordinal class probabilities for tBID as modeled by (i) two distinct pre-parameterized measurement models. In case 1, lowest and highest categories correspond to a narrow range of tBID values, while the three internal categories each account for roughly 1/3^rd^ of the tBID range. This parameterization might aim to account for effects of sensitivity and saturation on the measurement. In case 2, all five ordinal categories each account for 1/5^th^ of the range of tBID values. The right panels in Figures 4A and 4B show the assumed relationship between tBID concentration and probability of each ordinal category. Figures 4A and 4B (left plots) also show posterior predictions of tBID dynamics by aEARM calibrated to the ordinal dataset using these fixed pre-parameterized measurement models. The different measurement model pre-parameterization produced markedly different posterior predictions of tBID dynamics by the resulting aEARM calibrations. This raises potential concerns that assumptions in our interpretation of the measurement can artificially influence our interpretation of the mechanism.

**Figure 4:**
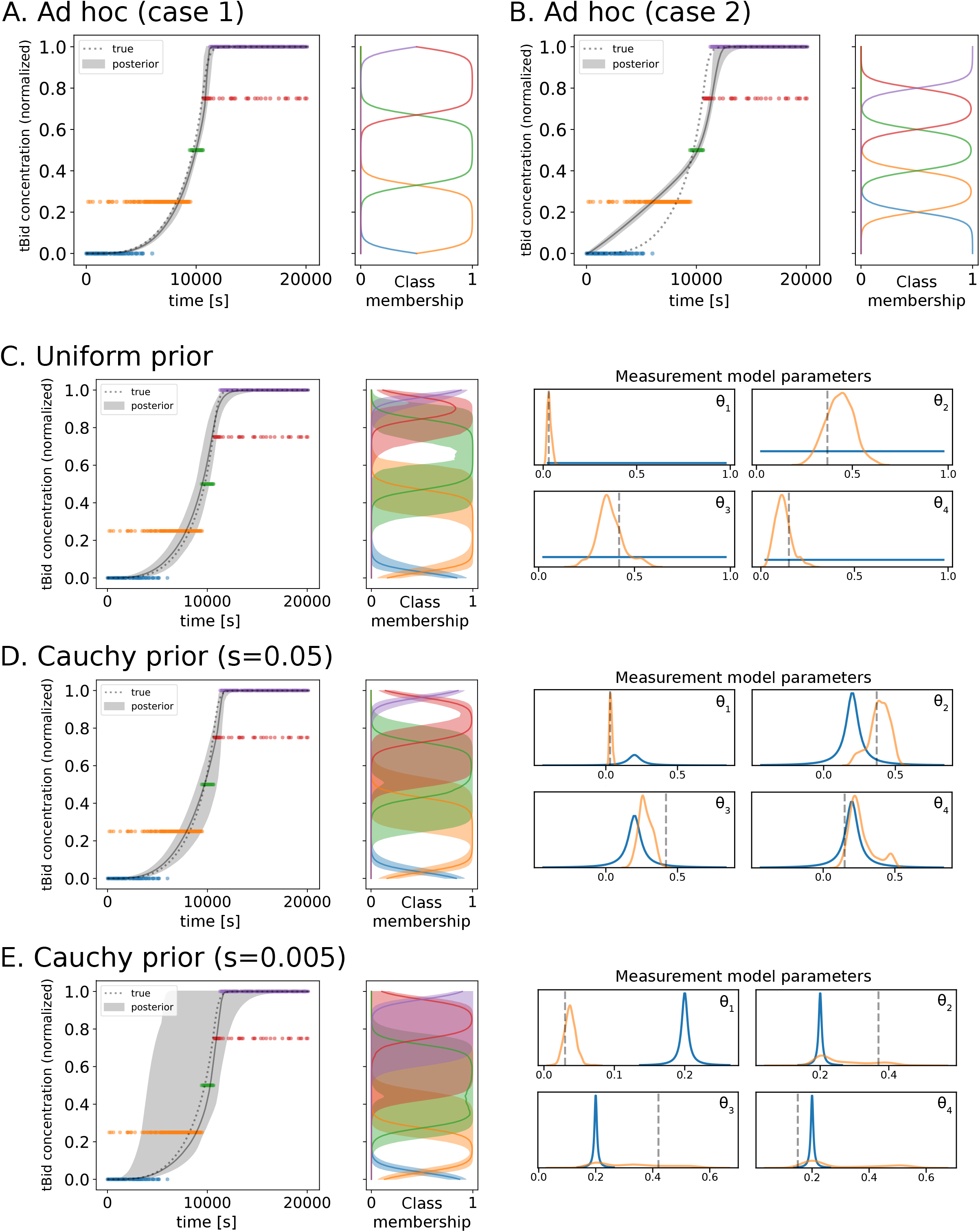
Predicted Bid truncation dynamics of aEARM trained to ordinal data using different measurement model parameterizations. A.) and B.) The 95% credible region of posterior predictions (shaded region) of tBID dynamics for aEARM calibrated to ordinal measurements two fixed parameterizations for the measurement model (see Supplemental Table 3). The adjacent panels plot the measurement models predicted probability of class membership (x-axis) as a function of normalized tBID concentration (y-axis). C.) D.) and E.) The 95% credible region of posterior predictions (shaded region) of tBID dynamics of aEARM calibrated to ordinal measurements uniform, Cauchy (scale=0.05) and Cauchy (scale=0.005) prior distributions for the parameterizations for the measurement model, respectively. In each, the median prediction (solid line) and true (dotted line) tBID dynamics are also shown. The adjacent panels give the 95% credible region of posterior predictions of the probability of class membership (x-axis) as a function of normalized tBID concentration (y-axis). Four accompanying plots show the prior (blue), posterior (orange) and true (dashed line) values of measurement model parameters.

In Figure 4B, the 50% probability boundary between adjacent categories occurs at every 0.2 interval; dividing the [0,1] range of tBID values into five equally spaced ordinal categories. Shown in Figure 4C-E, we represent this as a *flexible* assumption by encoding it in our priors (ii – iv) i.e., Cauchy distributions centered at every 0.2 interval (as detailed in *Methods*). The smaller the scale – more narrowly focused – prior distributions reflect less flexibility in the free-parameter and a stronger belief in our prior assumptions. Figure 4C-E shows the posterior predictions of tBID dynamics of aEARM calibrated to the ordinal dataset using increasingly more constrained priors on the measurement model parameters. The resulting posterior predictions of tBID dynamics were all less constrained than that of aEARM calibrated using fixed pre-parameterized measurement models (Figure 4A and 4B) but, they were more accurate as they contained the “ground-truth” tBID dynamics. Strongly constrained priors on the measurement model parameters (ii) produced a less certain mechanistic model; as indicated by its wider 95% credible region of posterior predictions of tBID dynamics (Figure 4E). The posterior distributions of measurement model parameters were spread out enough to give significant support of both the “ground truth” and the *a priori* assumed parameter values. This uncertainty in the measurement model parameter distributions translated into a less certain measurement model and less certain predictions of tBID dynamics. Weaker constraints on the measurement model parameters were encoded via larger scale prior distribution (Figure 4D). In Figure 4D we see these prior distributions, while centered on our *a priori* assumptions, includes the “ground truth” parameters. The posterior distributions of the measurement model parameters were therefore more constrained; likewise, the measurement model and posterior predictions of tBID dynamics had more certainty. This is also observed, in Figure 4C, the case where no prior assumptions were applied to the measurement model parameters (iv). The accuracy of the predictions of tBID dynamics comes from the flexibility of the data-driven measurement models’ parameters. This flexibility enables optimization (or prediction) of key properties of the measurement given the data. Figure 4 (right panels) shows the predicted probabilities of the ordinal categories (as a function predicted cellular tBID content); these predictions are accurate in that they contain “ground truth” probabilities. Using this approach, we calibrated more accurate models of mechanism by simultaneously learning a more accurate model of the measurement. This motivates us to further explore the data-driven measurement model as a potential new avenue for insights.

### Mechanistic insights from data-driven measurement models

We have shown thus far how a machine learning measurement model can reduce uncertainty and increase accuracy in model calibration. Through mechanistic model calibration to categorical data, we effectively employ machine learning classifiers to constrain mechanistic model dynamics to a corresponding categorical phenotype. We can then employ the measurement model in reverse, to better understand how properties of a biological mechanism predict, drive, and define a particular phenotype. This kind of knowledge would be essential for model-driven experimental data acquisition and model-guided validation.

To demonstrate this concept, we calibrated aEARM to synthetic nominal cell survival vs death data using a measurement model that estimated the contribution of variables in aEARM to the cell survival vs. death predictions. The survival vs death dataset was synthesized based on maximum log-rate of change of tBID and the time at which the rate of change maximized; these features were encoded into the measurement model, but their contribution was represented as a free parameter. In addition, the measurement model also considered the potential contribution of an unrelated variable (i.e., concentration of a reactant in reactions that occurred independently of the cell death ligand). Jointly calibrating aEARM and this measurement model to cell survival vs death data allowed data-driven predictions of how variables encoded in aEARM relate to cell survival vs death. Figure 5 shows posterior predictions of the values of potential predictors of cell survival vs death. The shaded region marks the 95% credible interval for the line marking 50% cell survival probability. Figure 5 (bottom row) provides the posterior distribution of weight coefficients for each the features encoded in the measurement model. (Larger absolute values of the weight coefficient indicate greater importance of the feature.) The calibrated measurement model correctly identified time at maximum Bid truncation as the most important predictor of our synthetic cell survival vs death data; and the unrelated variable as the least important predictor. Calibration of aEARM to the mixed dataset, described in the previous section, yielded a measurement model that equivalently predicted identified time at maximum Bid truncation as the most important predictor of cell survival; and the unrelated variable as the least important predictor. Calibration of a mechanistic model to categorical phenotype data, using data-driven measurement models, enabled correct identification of predictors (and potentially drivers or markers) of categorical phenotypes. The data-driven probabilistic measurement model we propose in this research was essential to this finding.

**Figure 5:**
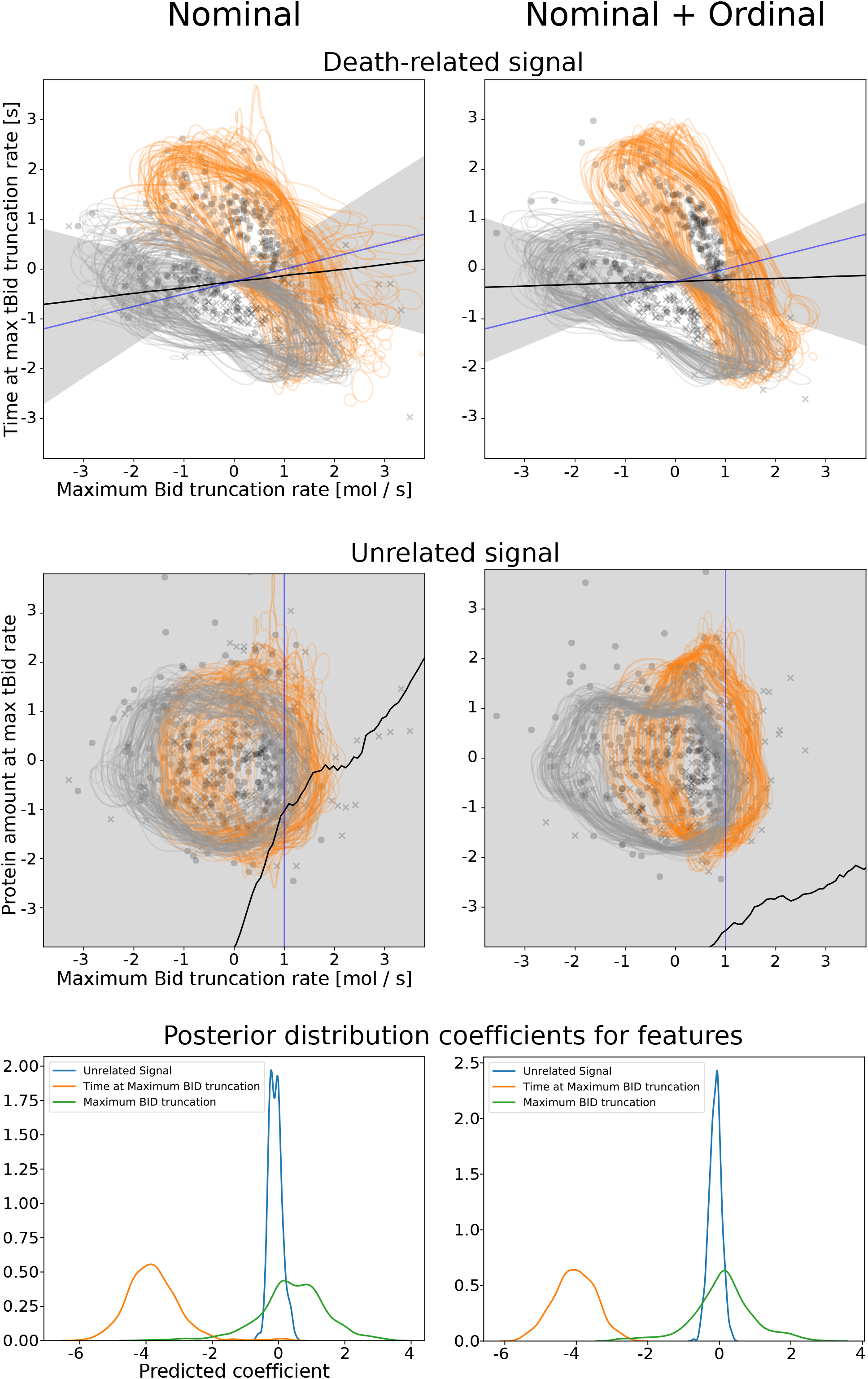
Measurement model predicts features of cell death vs. survival using aEARM calibrated to cell death datasets. Normalized predicted values of the features used in the cell death vs. survival measurement model – the x-axis is the maximum Bid truncation rate, and the y-axis is the time at maximum Bid truncation rate (top row) or an unrelated non-apoptotic signal (middle row) – for corresponding to observed cell death (x) and survival (o) outcomes. These feature values are modeled by aEARM parameterized by 100 parameter vectors randomly drawn from the posterior; for each parameterization, 5 out of the total simulated population of 400 cells were plotted. The grey and orange curves, in these plots, are 0.05 contours for the estimated density of simulated cell populations produced for each of the 100 parameter vectors – grey and orange correspond to 50 and 10ng/ml TRAIL treatments, respectively. The measurement model predicts a probability of cell death vs survival based on simulated values of the above features. The lower right region of the plots in the top row. (i.e., early maximization of Bid truncation and higher maximal Bid truncation rates) is associated with higher probability of cell death. The shaded region is the 95% credible region of the posterior prediction of the line marking 50% probability of cell death or survival. The black and blue lines are the median predicted and true 50% probability lines, respectively. The bottom row plots the posterior distributions of the weight for each feature (i.e., the product of the slope term and feature coefficient encoded in the measurement model): maximum Bid truncation rate (green), time at maximum Bid truncation (orange) and unrelated non-apoptotic signal (blue). Plots in the left column are predictions of aEARM calibrated to the cell death vs. survival dataset. Plots right column were those of aEARM calibrated to the cell death vs survival + ordinal IC-DISC combined dataset.

### Application of Data-Driven Measurement Model to Published Fractional Cell Death Data

To explore a practical application of the data-driven measurement model paradigm. We calibrated the aEARM parameters to published fractional cell death measurements (Figure 6A) by Wajant et al^31^. These measurements were taken in HeLa cell in which apoptosis was inhibited through graded expression of a dominant negative FADD. Wildtype, low- and high-dominant negative FADD genotypes were modeled (see *Methods*) via free parameters, *δ*_*low*_ and *δ*_*high*_, which represent diminished rates of DISC formation relative to wildtype *kc*_0_ in aEARM. These free parameters were calibrated along with the parameters of aEARM (and supporting the measurement model). *Supplemental Figure 21* shows the posterior distribution of these parameters.

**Figure 6:**
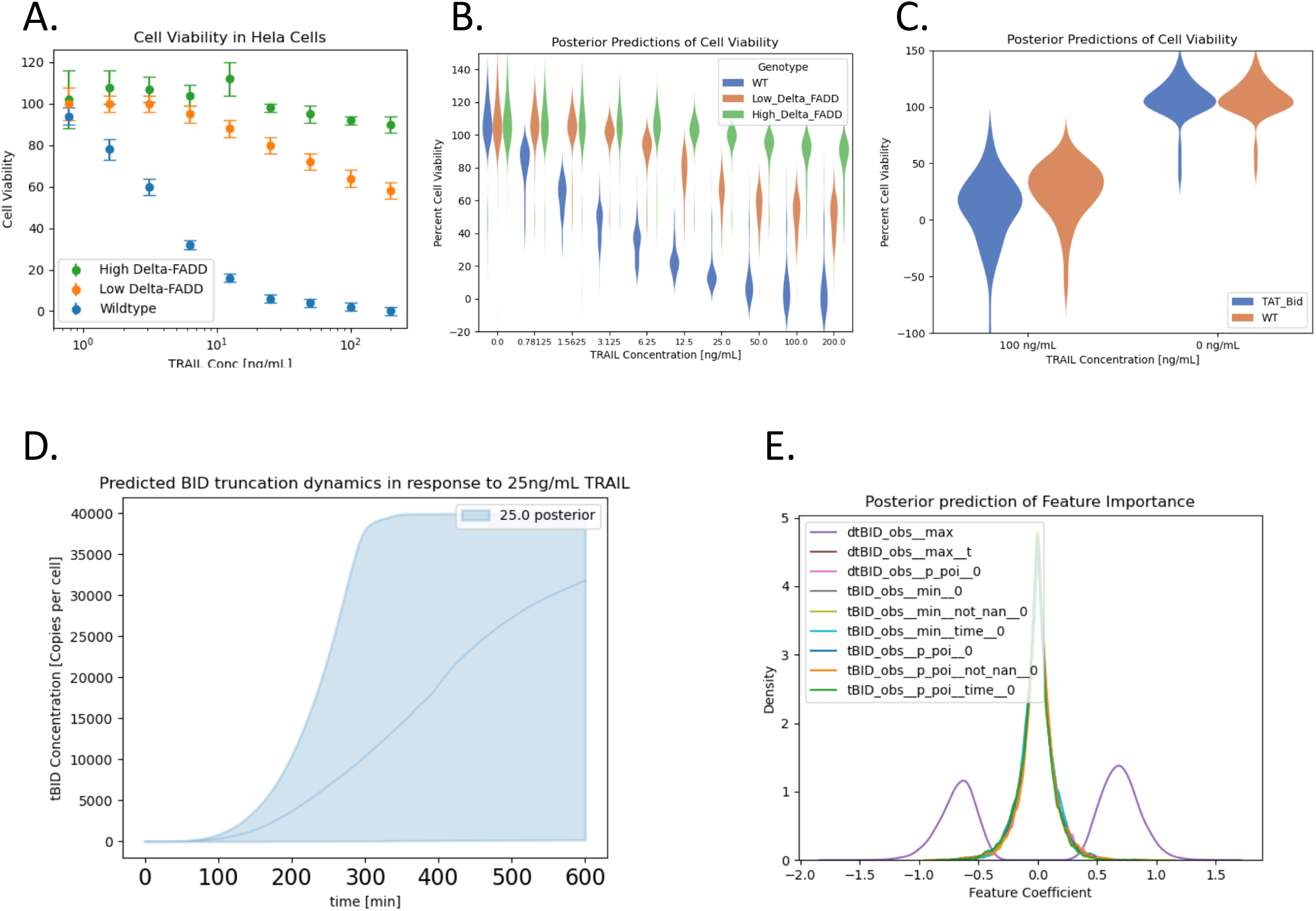
Posterior predictions of aEARM trained to published fractional cell death data. A. Fractional cell death in WT (blue) and high- and low-expression of dominant negative FADD (green and orange respectively) in HeLa cells treated with 0 to 200ng/mL TRAIL. These data come from Wajant et al. 1998^31^. The aEARM and accompanying measurement model were calibrated to these data. B. The posterior predictions of the Guassian process modeled mean fractional cell death values for WT and high- and low-expression of dominant negative FADD in HeLa cell treated with 0 to 200ng/mL TRAIL. C. Posterior predictions of the Guassian process modeled mean fractional cell death values for WT and BID overexpressed (TAT-Bid) HeLa cells treated with and without 100ng/mL TRAIL. Fractional cell death predictions for these experimental conditions, which were excluded from our training dataset, correspond to fractional cell death measurements by Orzechowska et al^32^. The 95% credible region of the posterior prediction (D.) of tBID dynamics in cells treated with 25g/mL TRAIL. (E.) Posterior distributions of the weight for each feature extracted from tBID dynamics.

Fractional cell death measurements are indirect semi-quantitative measurements since lack a clear mapping between the measured values and underlying cell death signaling dynamics. Specifically, there exists limited information about which features of the cell death signaling *dynamics* predict fractional cell death, and no established function to map those features into fractional cell death values. We therefore used Gaussian process (GP) regression to model the fractional cell death measurements. GP regression does not require we commit an assumed functional form for our measurement. Instead, it learns the posterior distribution over all functions that fit the data. This lets us flexibly apply Bayesian data-driven methods to our model of fractional cell death. In this way, GP model is a data-driven probabilistic measurement model like those demonstrated in previous sections. Our GP measurement model *y*(***x***)∼*GP*(*m*(***x***), *κ*(***x, x***′)) maps features of aEARM modeled tBID dynamics ***x*** onto fractional cell death values *y*(***x***). We used a radial-basis kernel (Eq. 14), for *κ*(***x, x***′)), with free parameters for the coefficient 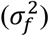 and variance matrix (*M*) and calibrated those parameters along with the rate parameters of aEARM. The variance matrix in this kernel (*M*) is the diagonal *β*^−2^, where *β* is the weight coefficient for the features extracted from tBID dynamics predicted by aEARM. The dynamics of tBID (as modeled by aEARM and others) are smooth, differentiable trajectories that can be accurately summarized via a small number of critical points (maximum, minimum, point of inflection, etc.). This low-dimensional critical points representation of tBID dynamics includes those features explored in Roux et al^19^. A single GP model was used for all data in the training set (see *Methods* for more information). The resulting likelihood function (i.e., the GP marginal likelihood Eq. 16) was used, with PyDREAM, to calibrate aEARM to fractional cell death measurements.

The posterior predictions of fractional cell death of the calibrated aEARM (Figure 6. B.) aligns with the training data (Figure 6. A.). Further, the calibrated model also accurately predicts fractional cell death observations outside of the training set: fractional cell death data in TAT-BID mutant HeLa cells (Figure 6. C.) as observed in Figure 6. G. of Orzechowska et al^32^. These predictions, however, had much less certainty which might reflect limited information provided in fractional cell death data. The 95% credible region of the posterior predictions of tBID dynamics (in response to treatment with 25 ng/mL TRAIL) of the aEARM trained to published fractional cell death measurements (Figure 6. D.) similarly occupies a large (or uncertain) range of values. However, this range is comparable to the range of trajectories of a caspase activity fluorescent indicator in HeLa cells treated with 25 ng/mL TRAIL, observed in Roux et al (Reprinted in *Supplemental Figure 22*)^19^. For instance, the predicted maximal tBID dynamics exhibit points of inflection between 200 and 300 minutes, which accurately reflects the observed maximal tBID trajectories (in HeLa cells treated with 25ng/mL TRAIL) by Roux et al^19^.

The fractional cell death measurement model considered critical point extracted from aEARM dynamics as predictive features of fractional cell death. Figure 6.E. shows the posterior distributions of feature coefficient weight *β*. The features with greater displacement from zero indicate increased predictive value. The fractional cell death measurement model therefore identified the maximum Bid truncation rate as the most important predictor of fractional cell death. This is consistent with observations by Roux et al. that changes in the maximum Fluorescence indicated caspase activity (proxy for BID truncation rate) imply sharp changes in observed probability of cell death (Reprinted in *Supplemental Figure 22*.)^19^. This result underscores the capacity of the data-driven measurement model to reveal insights into the potential predictors and drivers of cellular phenotype.

## Discussion

We used data-driven probabilistic measurement models to calibrate, using Bayesian methods, a dynamical model of biological mechanism to quantitative and nonquantitative data. Our approach allowed us to estimate posterior predictive regions for the calibrated models and to observe how the size of a dataset, its different measurement types, and our assumptions about the measurements affect model accuracy and certainty. Our findings support results from previous studies that suggest nonquantitative data are valuable for mechanistic modeling efforts^10-13^. For instance, a sufficiently large ordinal dataset can constrain the posterior predictions of a mechanistic model as much as quantitative dataset. However, far more nonquantitative data than is typically generated would be necessary for nonquantitative assays to match the information content of quantitative assays. In Figure 2B (second row), fourteen ordinal measurements of tBID – typical of common immunoblot measurements of intracellular biology – did not constrain the model around an accurate prediction of tBID dynamics. Instead, it took 24x as many ordinal measurements of tBID (336 measurements) to constrain the mechanistic model of apoptosis as well as the fluorescence dataset (112 measurements). We also found that datasets that combined categorical measurements of multiple variables in aEARM out-perform the datasets with measurements of an individual variable. These findings suggest one could overcome challenges posed by a dearth of quantitative data by devising experiments that, while nonquantitative, produce a larger number of diverse measurements that can cover multiple variables.

We also found the posterior predictions of our mechanistic model were sensitive to the assumptions, we encode in the measurement model, about the relationship between measurement and measurand. All measurements possess uncertain (or unknown) properties, but this uncertainty has a pronounced presence in nonquantitative measurements. The limitations of nonquantitative data exist because they impose less informative constraints on models, and this leaves room for biasing assumptions and/or uncertainty. Uncertainty in nonquantitative measurements drives the, often unacknowledged and implicit, assumptions about the relationship between measurement and measurand (i.e., between data and model). With the proposed Bayesian calibration framework, we could observe how assumptions about measurement affected the uncertainty and accuracy of the posterior predictions, in essence providing a measurable quality of how well the model can make mechanistic predictions. We found that inaccurate *ad hoc* assumptions about the measurement could produce models that suggested, with a higher degree certainty, an inaccurate prediction (Figure 4B). This finding suggests that *ad hoc* assumptions about measurements can lull practitioners into a false sense of confidence about the model and the data. This concern also motivated Schmiester and co-workers to avoid certain *ad hoc* assumption in their model calibration approach^11^.

Having a measurement model whose attributes are determined by data creates an opportunity to learn new details about the relationship between a measurement and its measurand(s). For instance, could a model of biological mechanism plus cell phenotype observations data enable identification of cell phenotype predictors? To explore this, we encoded a small number of suspected cell-fate predictors into our measurement model and let the data (and the mechanistic model) determine, through model calibration, their respective contribution to phenotype. In doing so, model calibration using our data-driven measurement model performed feature selection to correctly identify the most important predictor of cell death. We extend the data-driven measurement model to a practical example wherein published fractional cell death data were used to calibrate our mechanistic model of cell death. Here, we find calibration using these data produces a model that accurately predicts underlying cell death dynamics and correctly identifies features of those dynamics which best predict cell death. The high uncertainty of the model predictions, however, reflects the limited information contained in nominal cell fate observations and fractional cell death measurements. Leveraging larger datasets (of multiple datatypes) would likely improve the certainty of the mechanistic model and measurement model predictions. The measurements models deliberately use simple supervised machine learning models that capture the salient features of the measurement while maintaining tractability of the model calibrations. These characteristics lets us map cellular dynamics to diverse cellular phenotypes without requiring prohibitively large datasets (and computing resources). The measurement model can readily extend to mappings from intracellular dynamics to extracellular processes (e.g., tumor growth). We restricted our modeling to published observations of TRAIL induced apoptosis in HeLa cells where experimental treatments produced a definitive impact on the proteins modeled in aEARM. However, models of other cell types experimental systems etc. may supported by a larger set of available data sources. In general, this kind of measurement model, which relates mechanism to cellular phenotype, can be used to predict phenotype outcomes and identify potentially informative experimental conditions from *in silico* perturbation experiments.

### Conclusions

The present work presents an analysis and a proof-of-concept that can be improved upon in future work. We chose linear logistic classifiers, as they enable easy formulation of a likelihood function and application of Bayesian calibration methods, but other probabilistic classifiers could be used. We constrained our measurement representation to small number of potential features to avoid complications of high dimensionality to our machine learned measurement model. However, dimensionality reduction and feature learning (e.g., PCA) can, in theory, be integrated into the measurement model’s preprocessing and/or model calibration workflow. Possibilities for integrating more complex machine learning into models of measurement will depend on dataset size, computational power, and modeling goals.

Our work introduces the concept of measurement models to the mechanistic modeling paradigm. Measurement models have their origin in social sciences and statistics^30^. They also appear in more quantitative applications; some recent examples include management, manufacturing, and computer vision^33-35^. These measurement models can take on more complexity than the examples we provided, depending on the unique needs of the problems in these areas. The use of measurement models in these areas is motivated by a desire to define and quantify observations of nuanced and/or subjective phenomena; and connect those observations to an underlying theory. Biology, being “harder” than social sciences, but arguably “softer” than physics will straddle the technical domains of both. As a field, we face the same challenge as these social sciences given that our mechanistic models are situated within a larger context of explaining nuanced and subjective biological phenomena (e.g., cell-fate, morphology, physiology and overall health vs. pathology). As practitioners, we never encode *everything* into our mechanistic models; instead, there is always some aspect of the model (or its interpretation) that aims to connect back to these relevant biological phenomena. This fact ultimately motivates our application of data-driven probabilistic measurement models in our mechanistic models of intracellular biology.

## Acknowledgements

The authors would like to thank Dr. Alexander Lubbock, Dr. Leonard Harris, and Dr. Vito Quaranta, for insightful conversations and critical feedback on this work. This work was supported by the following funding sources: MWI was supported by the National Institutes of Health (NIH)[T32-GM139800]; CFL was supported by the National Science Foundation (NSF) CAREER Award [MCB 1942255]; and the National Institutes of Health (NIH) [U54-CA217450 and U01-CA215845].

## Methods

### Extrinsic Apoptosis Reaction Model

We built an abridged extrinsic apoptosis reaction model (aEARM) and trained it using PyDREAM to normalized fluorescence time-course data^21^. We built this abridged version of EARM to simplify convergence of Bayesian calibration algorithms and thus make feasible probability-based predictions on the model-data relationship^21^. The aEARM abstracts detailed mitochondrial reactions from the original model as two sequential mitochondrial outer membrane pore (MOMP) “signal” activation steps. In addition, apoptosome formation and effector caspase activation reactions take place in a single activation step. The aEARM does capture key dynamic characteristics, such as the snap-action delay dynamics of apoptotic effector molecules that is observed empirically^29^. For this work, three additional non-apoptotic species were encoded and linked via feedback activation and inactivation loops to test whether our data-driven measurement model could discriminate between drivers and non-drivers of apoptosis. (Supplemental Table 2). These additional species and reactions do not interact with any species or reaction in the aEARM model. The aEARM was encoded using rule-based modeling python package PySB^35^.

The aEARM parameters – initial conditions and rate coefficients – were adapted from the previously developed EARM and/or calibrated to fit available fluorescence data. Initial conditions parameters were lifted from the previously developed EARM (Supplemental Table 1). Previous work characterized extrinsic heterogeneity in the expression of proteins and its effect on apoptosis. To model extrinsic heterogeneity in apoptosis signaling, initial values of certain species (marked in table 1) were sampled from a log-normal distribution such that its mean equaled that in Supplemental Table 1 and coefficient of variation was 0.20. Rate coefficients were calibrated (described below) to fit normalized fluorescence time-course measurements of initiator and effector caspase reporter proteins (IC-RP and EC-RP respectively).

### Integrating aEARM Dynamics

Snap-action delay dynamics present challenges for Ordinary Differential Equation (ODE)-based models, as they feature rapid non-stiff to stiff transitions during integration. For this work we employed the LSODA integrator (from scipy, via the PySB solver suite), suitable for non-stiff/stiff systems^37^. However, we found that particularly poorly behaved parameter vectors could prolong integration evaluations in LSODA. Integrator settings were adjusted for efficiency and accuracy of integration as follows: mxstep (2^20), atol (1e-6 default), rtol (1e-3 default). The aEARM was integrated over a linear space of 100 time-points spanning 0 to 20160 seconds, in direct correspondence with the fluorescence time-course data^29^. Additional time-points in the data were obtained via linear interpolation.

### Measurement Models and Likelihood Functions

Likelihood formulations incorporated a measurement model and resulting distance metric for each datatype in the study: fluorescence time-course data, synthetic ordinal time-course data, and synthetic survival vs death binary data for a sample of 400 initial conditions. These likelihood functions were used to calibrate the models to each dataset. In addition to their use in the likelihood formulation, the measurement models, were also used to generate synthetic non-quantitative datasets.

We first trained the aEARM to normalized fluorescence time-course data for IC-RP and EC-RP, i.e. fluorescent proxies for substrates of initiator and effector caspase, respectively (i.e. Bid and PARP, respectively). Consistent with previous work, we defined a likelihood that assume an i.i.d. Gaussian-noise component *ϵ*∼*N*(0, *σ*^2^) on normalized tBID and cPARP predictions of the aEARM; where *σ*^2^ assumedly equals the variance of the data^21,38^. This yields a log-likelihood function (Eq. 11) where data the, *ŷ*, and normalized aEARM predictions, *y*, are compared for each time-point, *t*, and observable, *i* (i.e. tBID/IC-RP and cPARP/EC-RP). The aEARM trained to these fluorescence data served as the starting point in the synthesis of ordinal, nominal, mixed, etc. datasets, below.

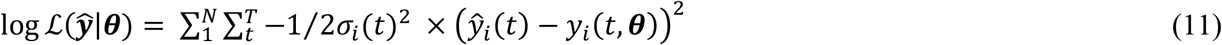

To train the aEARM to synthetic ordinal time-course data, a measurement model (i.e. that models the probability of each ordinal category as a function of an aEARM variable) was defined and applied in the formulation of a likelihood function^39^. The ordinal logistic regression python package, MORD, applies empirical ordering constraints to Scikit-Learn’s logistic regression class; this class then calculates a probability for each ordinal category^39^. The ordinal logistic model, encoded in MORD, defines ordinal constraints as a linear function of predicted values of an aEARM variable (e.g. *p*(*y*_*tBID*_ ≥ *c*_*j*_|*x*_*tBID*_) = *φ*(*αx*_*tBID*_ + *β*_*j*_) for aEARM variable, *x*_*tBID*_) where each ordinal constraint, *j*, is a logistic function *φ*(*z*) with a different offset coefficient, *β*_*j*_, but shared slope coefficient, *α*, for each of the ordinal categories. Each ordinal constraint function is combined, using the *sequential model* (i.e. the product of the logistic functions), to give a probability of each ordinal category, *P*(*y*_*i*_(*t*) = *c*_*j*_|*x*_*i*_(*t*, ***θ***), *α*_*i*_, *β*_*i,j*_)^40, 41^. These offset and slope coefficients are additional free parameters to be inferred in the model calibration. For example, a measurement model with *K* categories can be defined using *K* − 1 ordinal constraints and will therefore add a total of *K* free parameters (i.e. *K* − 1 offset coefficients and 1 shared slope coefficient) to the model. We also encoded error in our synthetic ordinal data by defining a 5% misclassification probability; i.e. we assume 95% probability the reported ordinal category, *c*_*j*_ = *ŷ*, and 2.5% probability of adjacent categories, *c*_*j*_ = *ŷ* ± 1, (5% for adjacent terminal categories). We model this by the marginal probability that the observation classified into the category predicted by the model: 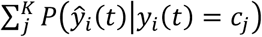^41^. Together, this yields a log-likelihood function (Eq. 12) where the probability of each category *c*_*j*_ is calculated for each time-point, *t*, and observable, *i*; and applied toward a likelihood of the data *ŷ* given the model. Where noted, we also trained the aEARM using measurement models with preset fixed parameters (Supplemental Table 3).

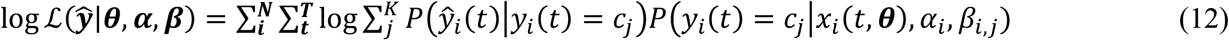

We trained aEARM to synthetic binary (survival vs death) data by incorporating a measurement model (i.e. logistic model of the probability of each categorical outcome) similar to that used for the ordinal data. We used the Scikit-Learn logistic regression class to model the probability of a cell-death outcome, *y* = *c*_1_, as a linear function of features, *x*_*l*_, derived from the aEARM simulation: 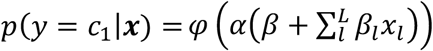, where *α* is a slope term, *β* is an intercept and *β*_*l*_ are weight coefficients for each of the *L* features^42^. Previous studies used *a priori* knowledge and assumptions about which features of a cell-fate marker’s dynamics to associate with the binary outcome. For instance, recent work delineates necrotic and survival cell fate outcomes using a threshold in the concentration of a known necroptosis marker (this assumption enabled models of necroptosis in the absence of an established relationship between the dynamics of the marker and commitment to necroptosis)^43^. Roux et al, investigated an empirical relationship between initiator caspase reporter protein (IC-RP), a fluorescent indicator of caspase activity or proxy for caspase substrate cleavage, and apoptosis in TRAIL stimulated HeLa cells^19^. They found, instead of concentration, the maximum rate of change in IC-RP and the time when that rate of change maximized better predicted the apoptosis-survival decision^19^. The features we use in our study are based on findings by Roux et al^19^. The features are derived from aEARM simulated tBID dynamics, *x*_*tBID*_ (*t*, ***θ***): time at maximum rate of change, and log-maximum rate of change. To test the measurement model’s ability to discriminate between predictors and non-predictors of cell death, we encoded an additional feature: the concentration of an unrelated non-apoptotic species (USM2 in Table 2) when bid truncation maximizes. Together this totals three features. We interpret each observation in the dataset as an independent Bernoulli random variable. Each cell death vs survival observation is compared with these three features, *x*_*l,m*_, extracted from an aEARM trajectory that was simulated from a unique vector of initial conditions. There were 400 observations; 2 sets of 200 observations corresponding to 10 and 50ng/mL initial ligand concentration. Together, this yields a log-likelihood function (Eq. 13) where each, *m*, of the *M* aEARM simulated trajectories corresponds to an observation *ŷ*_*m*_. Given the definitiveness of observed surviving vs dead outcomes, we considered the chance of misclassification to be zero (i.e., *P*(*ŷ*_*m*_|*y*_*m*_ = *c*_1_) = 0 when *ŷ*_*m*_ ≠ *c*_1_).

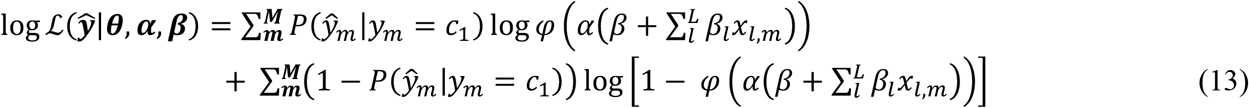

We trained aEARM to published fractional cell death data by Wajant et al^31^. We used the the Scikit-Learn Gaussian process regression ***y***(***x***)∼*GP*(*m*(***x***), *κ*(***x, x***′)) to maps variables in aEARM to values of fractional cell death in the data *y*(***x***). In this case, ***x*** is vector of features extracted from aEARM predicted tBID dynamics (described below). The kernel in our Gaussian process model, *κ*(***x, x***′), is a radial-basis kernel (Eq. 14), where *x*_*p*_ and *x*_*q*_ are features corresponding to two points in the data (*y*_*p*_ and *y*_*q*_) and 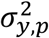 is observed standard deviation at *y*_*p*_. A single Gaussian process model was applied to all the points in the fractional cell death dataset. The coefficient 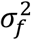 and variance term *M*, (Eq. 15) are free parameters that are estimated in the calibration. There variance term (Eq. 15) is modeled as the diagonal matrix where ***β*** are the weight coefficients for the features extracted from tBID dynamics predicted by aEARM^44^. The features we use in our model critical points (i.e., relative maxima and minima, points of inflection), the rate of change at the point of inflection, and/or half-maximal point on the aEARM modeled tBID trajectories. Different parameterizations of aEARM provide different sets of critical points. To keep a consistent parameterization of the features weights, a look-up table was employed where new critical points were appended to the table as they arose during model calibration. The look up table took a maximum of ten features. We modeled the likelihood of the data given the model (and measurement model) as the log-marginal likelihood of the Gaussian process model given the data (Eq. 16); where the kernel which is a function of the model parameters *θ* and the measurement models *θ*_*M*_.

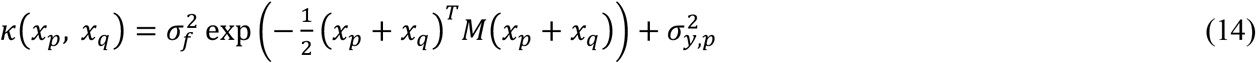

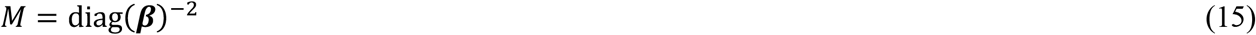

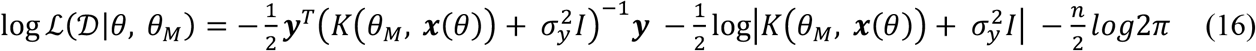

### Generating Synthetic Datasets

The calibration of aEARM to IC-RP and EC-RP fluorescence time-course data provided an optimally fit vector of rate coefficient parameters, which served as the “ground truth” parameter vector in the synthesis of the nonquantitative datasets (Supplemental Table 5). These parameters were applied to aEARM, and the resulting aEARM was used simulate time-courses for variables to be indicated in the nonquantitative data: truncated BID (tBID), initiator caspase localization to the death inducing signaling complex (IC-DISC), and cleaved PARP (cPARP).

These time-courses were converted to ordinal time-course datasets. The effective bit resolution of a measurement technology dictates how many unique values it can distinguish^45^. The total number of ordinal categories, *K*, was set such that resulting dataset had less than 70% of the effective bit resolution, *EBR*, (Eq. 14) of the IC-RP of EC-RP data. The signal to noise ratio, *SNR*, (Eq. 15) assumes the data, *d*, were subject to Gaussian noise and a 0.10 misclassification rate between adjacent values; modeled as the 0.95 quantile of a unit normal distribution^45^. Therefore, the number of ordinal categories were 5 and 4 for tBID and cPARP, respectively. The number of ordinal categories for IC-DISC were arbitrarily set to 4. Arbitrary values of slope and offset coefficients (Supplemental Table 3) were designated “ground truth” and applied to ordinal measurement models (described above). The resulting measurement models map the values in the aEARM simulated time-courses to probabilities of each ordinal category. These probabilities were used to simulate random class assignments for synthetic ordinal datasets (see Fig 2). The aEARM was trained to time-course ordinal values of tBID and cPARP or time-course ordinal values of IC-DISC and nominal data described below.

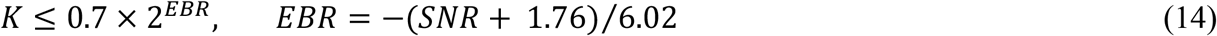

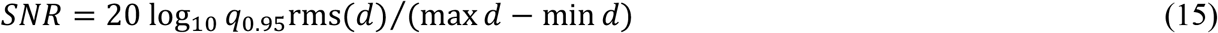

To generate synthetic nominal (binary cell survival vs death) data, two heterogeneous populations of 200 aEARM tBID (and an unrelated non-apoptotic species, USM2) trajectories were simulated from ground truth parameters. The populations had distinct initial ligand concentrations (10 or 50 ng/mL). Heterogeneity was modeled by a log-normal random sample of certain initial conditions (described above). These time-courses were preprocessed to yield values of the features encoded in nominal measurement model, above. This measurement model (which was encoded with preset “ground truth” values of slope, intercept and weight coefficients – See Supplemental Table 4) maps these features to probabilities of the binary outcomes. These probabilities were used to simulate random class assignments for synthetic nominal datasets (Fig 3b).

To generate a synthetic distribution of times at which Bid truncation was half-maximal, two heterogenous populations of 200 aEARM tBID time-courses, corresponding to 10 and 50ng/mL initial ligand concentrations, were simulated from ground truth parameters (as above). Time at half-maximal tBID was calculated via linear interpolation and rounded to the nearest 3-minute time-point (i.e. to reflect temporal resolution of common time-series intracellular experiments) (Fig 3a).

### Fractional Cell Death Dataset

We calibrated the aEARM parameters to published fractional cell death measurements (Figure 6. A.) taken Wajant et al via visual inspection of the published figure^31^. These measurements were taken of HeLa cell in which apoptosis was inhibited through graded expression of a dominant negative FADD. A test set of fractional cell death data was taken from Orzechowska, et al.,)via visual inspection of the published figure^32^. The test data were measurement of HeLa cells in which Bid expression was overexpressed (i.e., TAT Bid HeLa cells). Bid expression in TAT Bid HeLa was modeled as 20% increased Bid (which is consistent with the observations of Orzechowska, et al.)^32^.

### Model Calibration via Bayesian Inference

The aEARM was calibrated using DREAM(ZS) algorithm for all datasets^46^. Rate parameters in aEARM were given independent log-normal prior probability functions with a location equal to the ground-truth parameter vector and a scale term of 1.5. Wildtype, low- and high-dominant negative FADD genotypes were modeled via free parameters, *δ*_*low*_ and *δ*_*high*_, which represent diminished rates of DISC formation relative to wildtype *kc*_0_ in aEARM. These parameters, *δ*_*low*_ and *δ*_*high*_, were given independent truncated log-normal prior probability function with locations equal to 0 and -1 respectively. The prior probability functions were truncated at 0 and negative infinity. The nominal (cell death vs survival) dataset features a heterogeneous population of values. We modeled this heterogeneity with a random sample of initial conditions (described above). This random sample was shifted and scaled according to inferred values of the model mean and variance. The mean (if estimated) was given a log-normal distribution prior probability function with a location equal to ground-truth and a scale term of 1.5. The extrinsic noise (or variance) was given inverse gamma distribution with *a* and *b* terms such that the resulting coefficient of variation had a prior mean and standard deviation of 0.20 and 0.015 respectively.

Prior probability functions were also applied to the measurement models’ free-parameters. To in encode empirical ordering constraints on the ordinal measurement model, the slope terms, *α*, were greater than zero; they were given independent exponential distribution prior probability functions (with location of 0.0 and scale of 100.0). To insure monotonically increasing offset terms, each offset, *β*_*j*_, was defined by the distance, *θ*_*j*_, from its preceding offset term; *β*_*j*_ = *β*_*j*−1_ + *θ*_*j*_. The first offset, *β*_0_ = *θ*_0_, and subsequent distance, *θ*_*j*_, terms were given independent exponential distribution prior probability functions (with location of 0.0 and scale of 0.25). We explored the effect of increasingly biased priors on the ordinal measurement model parameters. Where noted, the slope terms, *α*, were given increasingly constrained independent prior probability functions: uniform (0.0 - 100.0 bounds), Cauchy (50.0 location and 10.0 scale) and Cauchy (50.0 location and 1.0 scale). The offset, *β*_0_, and distance, *θ*_*j*_, terms were similarly given independent uniform (0.0 - 1.0 bounds), Cauchy (0.2 location and 0.05 scale) and Cauchy (0.2 location and 0.005 scale) distribution prior probability functions. Parameters for the measurement model were given independent Laplace distribution prior probability functions with a location of 0.0 and scale of 1.0 for the slope, *α*, and 0.10 for the intercept and weighting coefficients, *β*and *β*_*l*_.

The likelihood functions were described above. Additional settings applied to the DREAM(ZS) algorithm were: number of chains (4) number of crossover points (nCR=25), adaptive gamma (TRUE), probability of gamma=1 (p_gamma_unity=0.10), gamma term resolution (gamma_levels=8). A burn-in period wherein crossover weights are adapted was set to 50,000-step burn-in for ordinal datasets and 100,000+ step burn-in. The calibration algorithm continued until it reached the stopping criterion: when the Gelman-Rubin metric (calculated on the latter 50% of the traces) was less than or equal to 1.2 for all free-parameters in the model; at which point the parameter traces were considered converged^32^. Gelman-Rubin metrics for each calibration are listed in Supplemental Table 6. Model calibrations were run on a x64 Intel with 32 total CPU threads (256GB RAM) and x64 AMD with 256 threads (1024GB RAM). Run times varied widely given the stochastic nature of the optimization algorithm but were typically one to seven days for simple model calibrations. Random samples of 1000 parameter were taken from the latter 50% of the resulting parameter-traces were used in subsequent analyses. Source code for the model calibrations as well as code for downloading the resulting parameter-traces is found at https://github.com/LoLab-VU/Opt2Q.

### Model Predictions

We simulated the equal-tailed 95% credible region of the posterior predictions of aEARM via samples of the model parameters posterior distribution. This was done by randomly generating 1000 parameter sets sub-sampled from the posterior sample of parameters generated via PyDREAM. For each parameter set, tBID time-courses (and/or cPARP, IC-DISC) were simulated from aEARM. The 95% credible region of the predictions was then determined via 0.025 and 0.975 quantile bounds on the tBID (or other variables) values for each time-point in the simulated time-course. The area bounded in the 95% posterior credible interval was determined by summing the difference between the 0.025 and 0.975 quantile bounds across 100 equally spaced time points on the trajectory. The 95% posterior credible intervals on the measurement model predictions were similarly the described by calculating 0.025 and 0.975 quantile boundaries on the predictions of the measurement model parameterized via 1000 parameter set samples from a posterior. This includes the posterior probability distributions of the feature coefficients encoded in the nominal measurement model. To model predictions of the nominal dataset, however, we randomly generated 100 parameter sets via sub-sampling of the posterior parameter distribution. For each parameter set, we simulate tBID dynamics from the set of 400 initial conditions as described above; from that we compute maximum BID truncation rate and time at maximum BID truncation rate for each of the 400 trajectories. The 0.05 contour of the KDE of the resulting 400 values of maximum BID truncation rate and time at maximum BID truncation rate was plotted for each of the 100 parameter sets.

We also simulated posterior predictions fractional cell death. This was done by randomly generating 1000 parameters sets subsampled from the posterior sample of parameters generated via PyDREAM. For each parameter set tBID time-courses were simulated from aEARM and preprocessed to extract a critical points representation of the tBID dynamics. These features were supplied to the parameterized Gaussian process model of our fractional cell death measurements, which returned a prediction for the mean fractional cell death. Violin plots of the KDE of the resulting sample of 1000 posterior predictions of fractional cell death were plotted in Figure 6.

