## Supplemental Materials for "Predictive uncertainty in mechanistic models of cellular processes calibrated to experimental data"

**Supplemental Methods Details**

Table 1 - Initial conditions:

| Species | Symbol | Value | Notes |
|---|---|---|---|
| Ligand (TRAIL) | 'L_0' | 3000 | 3000 copies per cell equates to 50ng/mL |
| Receptor | 'R_0' | 200 | |
| DISC | 'DISC_0' | 0 | The death inducing signaling complex (DISC) is initially absent |
| Initiator Caspase | 'IC_0' | 2.0e4 | Caspase 8 |
| Effector Caspase | 'EC_0' | 1.0e4 | Caspase 3 |
| MOMP signal* | 'MOMP_sig_0' | 1.0e5 | Uses the EARM initial value for Smac |
| PARP | 'PARP_0' | 1.0e6 | PARP (Caspase 3 substrate and apoptosis marker) |
| Three Unrelated* Signaling Molecules | 'USM1_0' | 1.0e3 | These signaling molecules do not interact with aEARM and are linked together via a separate set of activation and inactivation reactions (Supplemental table 2) |
| | 'USM2_0' | 1.0e3 | |
| | 'USM3_0' | 1.0e3 | |

*indicates values that were subjected log-normal extrinsic noise. Additionally, rate parameter 'kc0' (which catalyzes the formation of DISC) was subjected to extrinsic noise using the same procedure as for those noted in the table.

Table 2 – Reactions for unrelated signaling molecules

| | |
|---|---|
| USM1 catalyzes USM2 activation | $USM1^* + USM2 \rightleftharpoons USM1^*{:}USM2 \longrightarrow USM1^* + USM2^*$ |
| USM2 catalyzes USM3 activation | $USM2^* + USM3 \rightleftharpoons USM2^*{:}USM3 \longrightarrow USM2^* + USM3^*$ |
| USM3 catalyzes USM1 activation | $USM3^* + USM1 \rightleftharpoons USM3^*{:}USM1 \longrightarrow USM3^* + USM1^*$ |
| USM1 catalyzes USM3 inactivation | $USM1^* + USM3^* \rightleftharpoons USM1^*{:}USM3^* \longrightarrow USM1^* + USM3$ |
| USM2 catalyzes USM1 inactivation | $USM2^* + USM1^* \rightleftharpoons USM2^*{:}USM1^* \longrightarrow USM2^* + USM1$ |
| USM3 catalyzes USM2 inactivation | $USM3^* + USM2^* \rightleftharpoons USM3^*{:}USM2^* \longrightarrow USM3^* + USM2$ |

* indicates activated.

Monotonically increasing values would correlate highly with TRAIL dependent species in aEARM, as an artifact of the aEARM species all having nonequilibrium values at t=0. This activation and inactivation scheme can produce slow oscillations, which decrease their correlation with TRAIL dependent species in aEARM.

Table 3 – Parameterizations of Ordinal Measurement Model

| | |
|---|---|
| "Ground Truth Parameterization" | $\alpha_{tBID} = 50.0, \boldsymbol{\beta}_{tBID} = (0.03, 0.40, 0.82, 0.97)$<br>$\alpha_{cPARP} = 50.0, \boldsymbol{\beta}_{cPARP} = (0.03, 0.20, 0.97)$<br>$\alpha_{IC-DISC} = 50.0, \boldsymbol{\beta}_{IC\_DISC} = (0.05, 0.40, 0.85)$ |
| *Ad hoc* parameterization (Case 1) | $\alpha_{tBID} = 50.0, \boldsymbol{\beta}_{tBID} = (0.00, 0.33, 0.67, 1.00)$<br>$\alpha_{cPARP} = 50.0, \boldsymbol{\beta}_{cPARP} = (0.00, 0.50, 1.00)$ |
| *Ad hoc* parameterization (Case 2) | $\alpha_{tBID} = 50.0, \boldsymbol{\beta}_{tBID} = (0.20, 0.40, 0.60, 0.80)$<br>$\alpha_{cPARP} = 50.0, \boldsymbol{\beta}_{cPARP} = (0.25, 0.50, 0.75)$ |

Table 4 – Ground Truth Parameterization of Nominal Measurement Model

| | |
|---|---|
| "Ground Truth Parameterization" | $\alpha = 4.0$ (slope term)<br>$\beta = -0.25$ (intercept term)<br>$\beta_{unrelated\ signal} = 0.0$<br>$\beta_{\max tBID\ rate} = 0.25$ |

| | |
|---|---|
| | $\beta_{time\ at\ \max tBID\ rate} = -1.0$ |

Table 5 – Ground Truth Parameterization of aEARM

| | | | | | | | |
|---|---|---|---|---|---|---|---|
| kf0 | 1.15594933e-07 | kr2 | 6.87631084e-02 | kc4 | 3.39059104e-03 | kf7 | 1.98195485e-05 |
| kr0 | 1.04056134e-05 | kc2 | 1.09271827e-02 | kf5 | 8.69183082e-10 | kr7 | 1.82182416e-04 |
| kc0 | 1.12028665e-05 | kf3 | 3.82096809e-06 | kr5 | 1.15619903e-03 | kc7 | 3.90039620e-03 |
| kf1 | 1.93083645e-06 | kr3 | 1.39230000e-05 | kc5 | 8.49612026e-06 | kf8 | 1.76040074e-05 |
| kr1 | 1.71446702e-03 | kc3 | 2.95330594e-02 | kf6 | 7.69174202e-07 | kr8 | 1.34854804e-03 |
| kc1 | 3.29355599e-01 | kf4 | 1.61609297e-05 | kr6 | 7.35417590e-04 | kc8 | 1.67799299e-01 |
| kf2 | 1.48867282e-08 | kr4 | 1.71818138e-02 | kc6 | 5.18485726e-03 | kc9 | 8.28571278e-08 |

Table 6 – Gelman Rubin Values for each model calibration
Attached .xlsx file

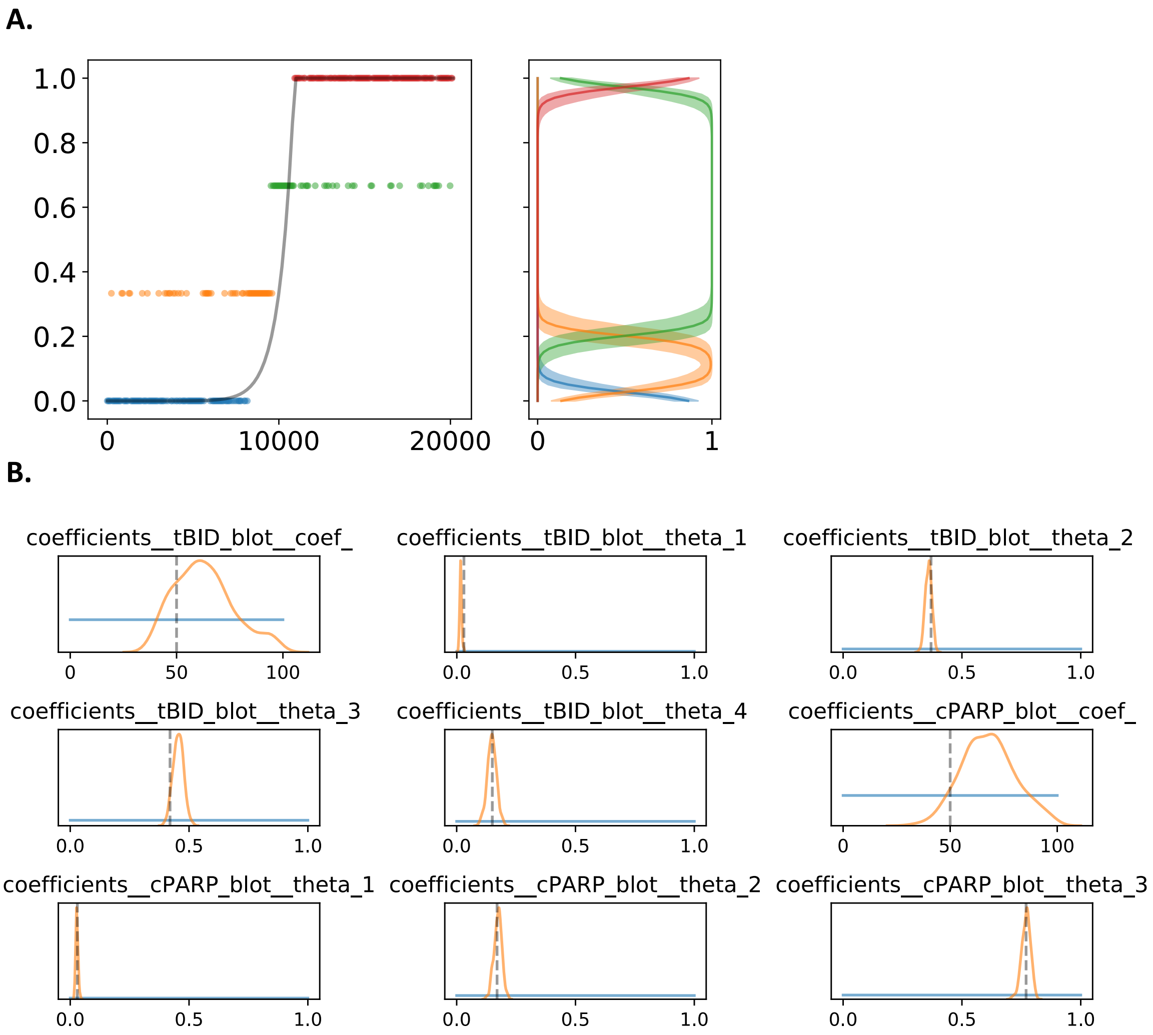

**Supplemental Figure 2: Measurement model parameters calibrated to an ordinal dataset.** The ordinal dataset contained a single ordinal value for tBID (Box 2, B.) and cPARP (A.) at every 60 second interval. Prior and posterior distributions for measurement model parameters are plotted in C. The 95% credible region of posterior predictions (shaded region) for the measurement model for tBID (Box 2, C.) and cPARP (B.) are plotted. The solid line in these plots is the median prediction for the measurement model. These plots give estimates of the probability of the ordinal value (x-axis) as a function of the normalized value of tBID or cPARP (y-axis). The ordinal categories are color coded and plotted in ascending order (for example, the ‘blue’ category is lower than the ‘orange’.

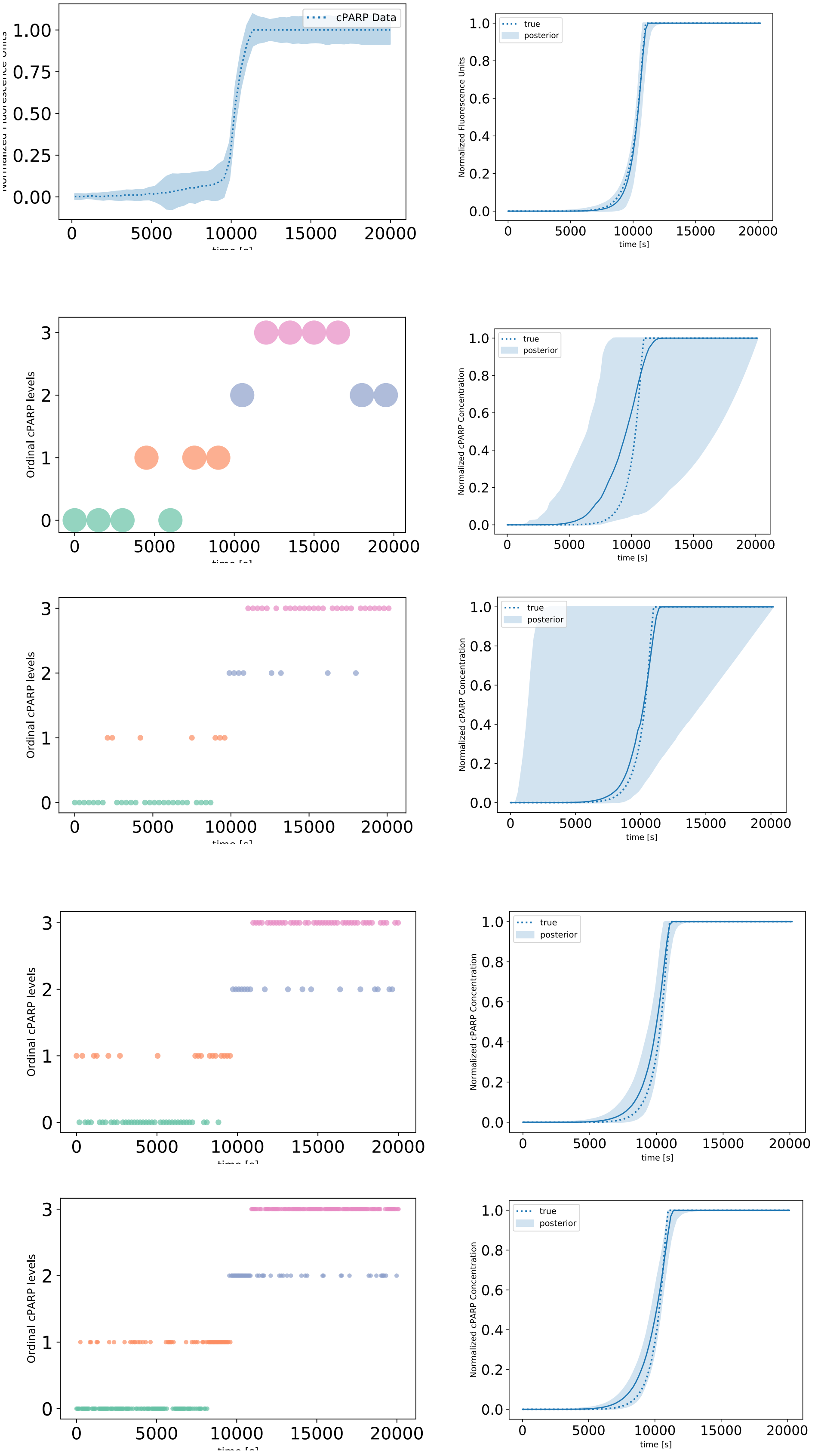

cPARP Data
0.00
0.25
0.50
0.75
1.00
0
5000
10000
15000
20000
true
posterior
Normalized Fluorescence Units
time [s]
0.0
0.2
0.4
0.6
0.8
1.0
Ordinal cPARP levels
0
1
2
3
Normalized cPARP Concentration
true
posterior
Normalized cPARP Concentration
time [s]
Ordinal cPARP levels
true
posterior
Normalized cPARP Concentration
time [s]
Ordinal cPARP levels
true
posterior
Normalized cPARP Concentration
time [s]
Ordinal cPARP levels
true
posterior
Normalized cPARP Concentration
time [s]

**Supplemental Figure 3: PARP cleavage dynamics of aEARM trained to Fluorescence EC-RP data.** Bayesian optimization of aEARM parameters trained to A.) Effector caspase reporter (EC-RP) fluorescence time-course measurements at 180s intervals (i.e. a proxy for PARP cleavage) and IC-RP fluorescence time-course (Fig 2A) data from [REF]). The plot shows the mean (dotted line) +/- 1 standard deviation (shaded region) for each time point. B.) The 95% credible region of posterior predictions (shaded region) for cPARP concentration in aEARM, calibrated to fluorescence measurements of IC-RP and EC-RP data. The median prediction (solid-line) and true (dotted line) cPARP concentration trajectories are shown. C., E., G., and I.) Ordinal measurements of cPARP at occurring at every 1500, 300, 180 and 60s timepoint, respectively. The 95% credible region of posterior predictions of cPARP dynamics for aEARM calibrated to ordinal measurements of tBID and cPARP occurring at every 1500, 300, 180 and 60s timepoint, respectively. The 95% credible region of predictions (shaded region), median prediction (solid line) and true (dotted line) cPARP dynamics for aEARM calibrated to ordinal measurements of tBID and cPARP occurring at every 1500, 300, 180 and 60s timepoint, are plotted in D., F., H., and J., respectively. The plots for tBID ordinal measurements and predictions are found in Figure 2.

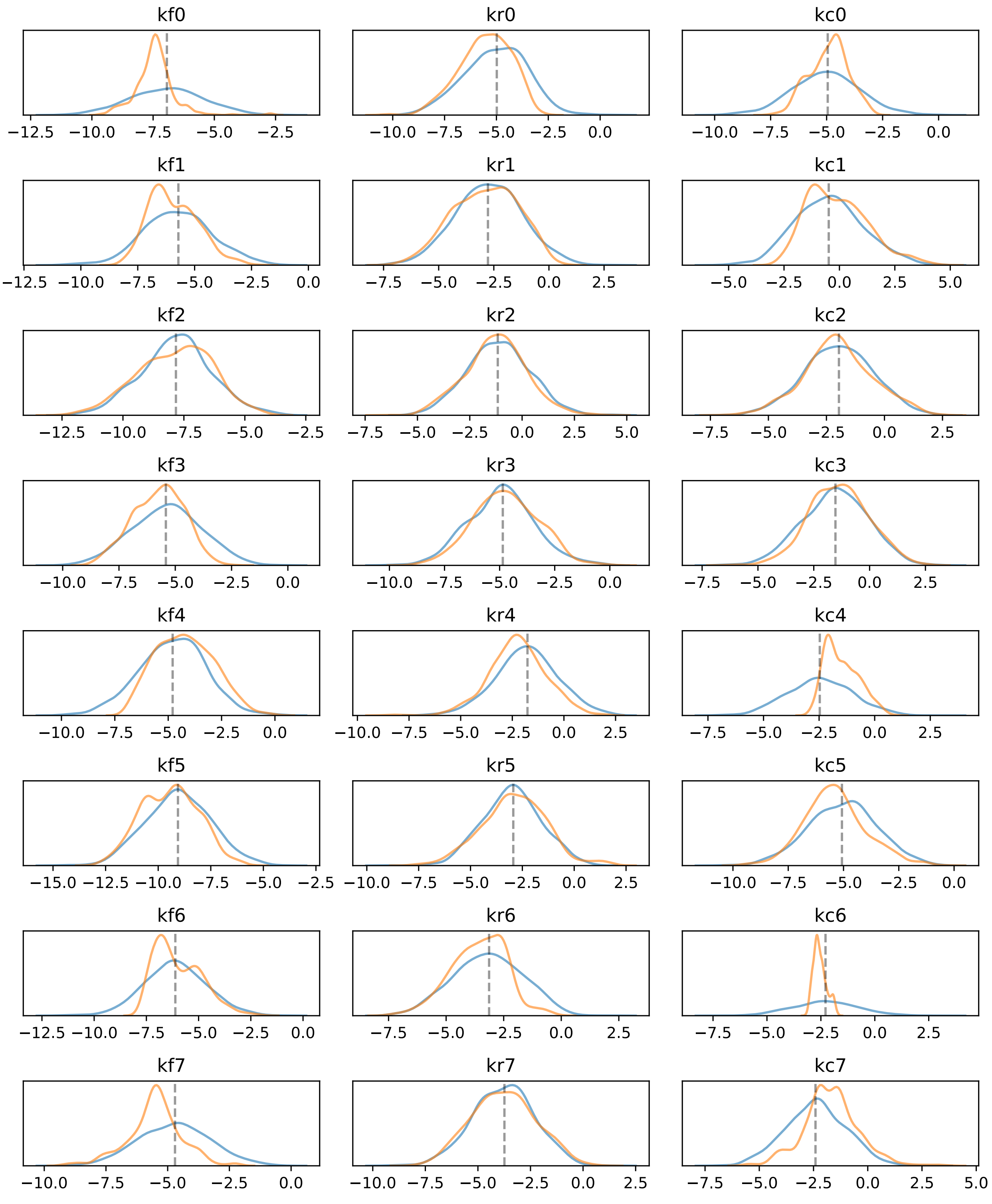

**Supplemental Figure 4A: Model parameters calibrated to a Fluorescence Dataset** Parameters for aEARM were calibrated to fluorescence time-course measurements of EC-RP and IC-RP at every 180s interval. Prior (blue) and posterior (orange) distributions $\log_{10}$ of the value of parameter are shown.

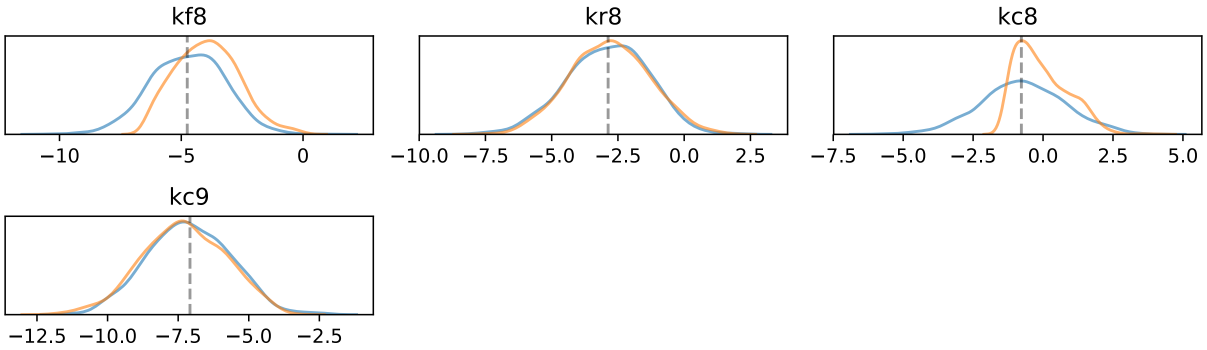

**Supplemental Figure 4B: Model parameters (remaining) calibrated to a Fluorescence Dataset**
Parameters for aEARM were calibrated to fluorescence time-course measurements of EC-RP and IC-RP at every 180s interval. Prior (blue) and posterior (orange) distributions $\log_{10}$ of the value of parameter are shown.

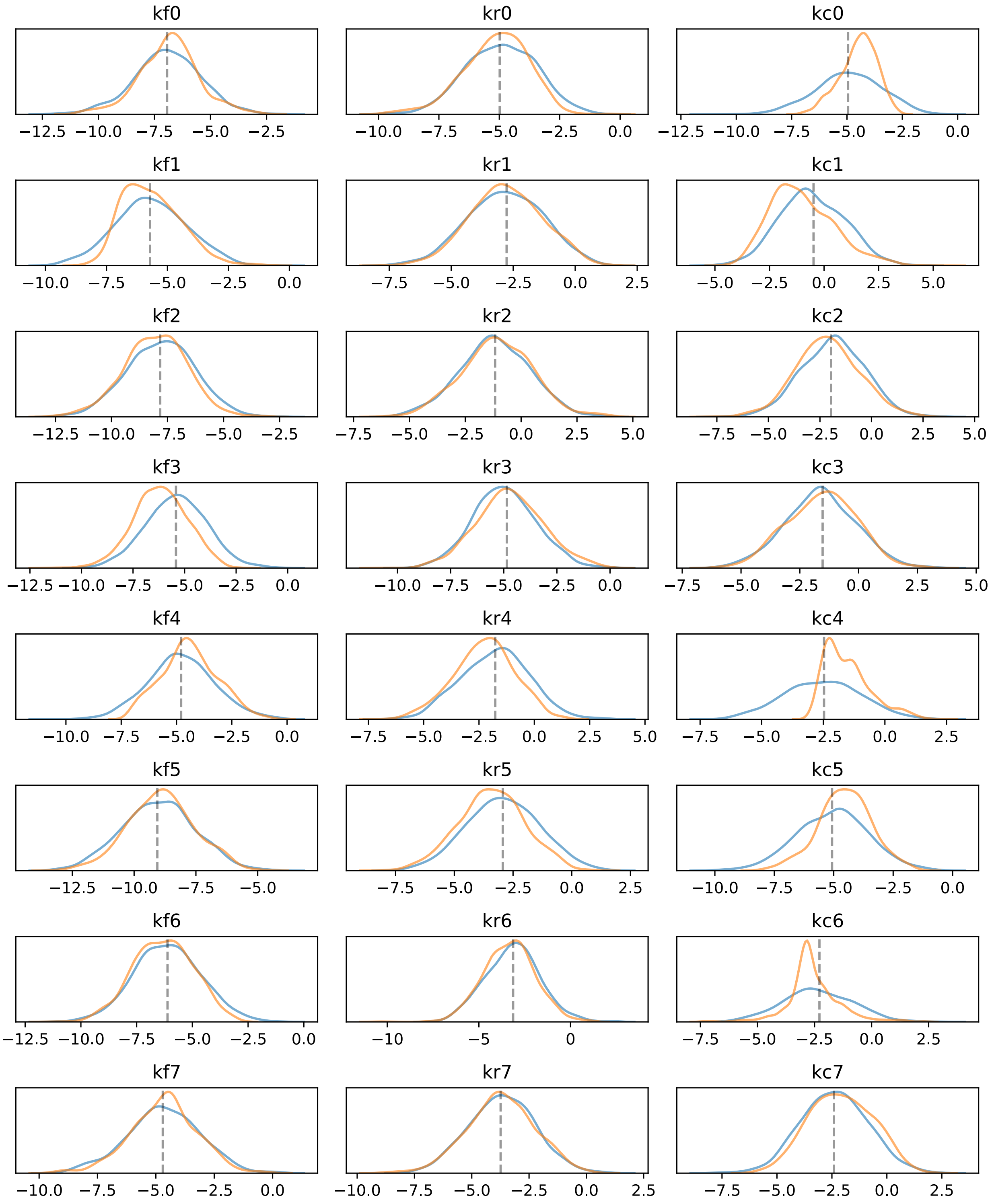

**Supplemental Figure 5A: Model parameters calibrated to an Ordinal Dataset** Parameters for aEARM were calibrated to ordinal values of tBID and cPARP abundance at every 60s interval. Prior (blue) and posterior (orange) distributions $\log_{10}$ of the value of parameter are shown.

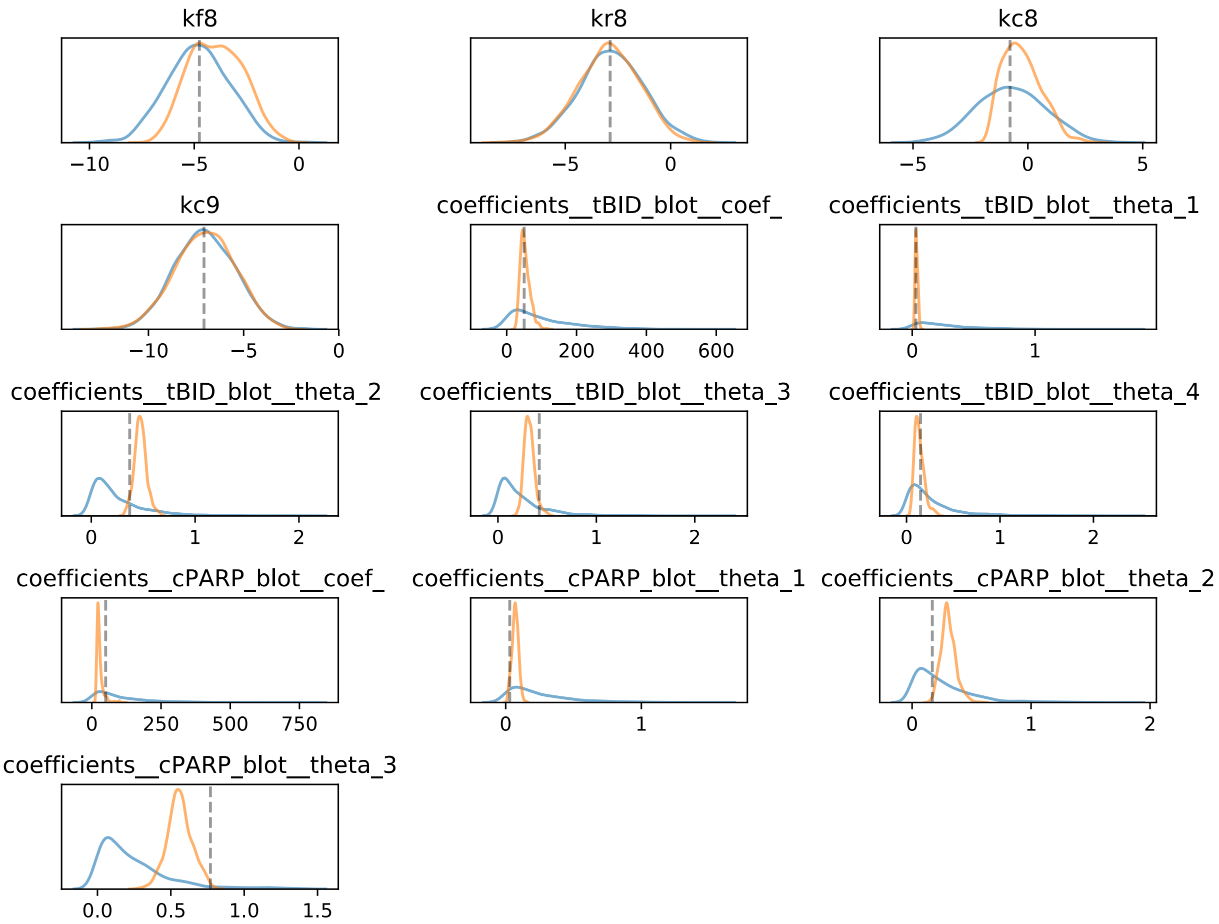

**Supplemental Figure 5B: Model parameters calibrated to an Ordinal Dataset** Parameters for aEARM were calibrated to ordinal values of tBID and cPARP abundance at every 60s interval. Prior (blue) and posterior (orange) distributions $\log_{10}$ of the value of the aEARM parameter are shown. Prior and posterior distributions of the value of measurement model coefficients are also shown (these are not log-scale).

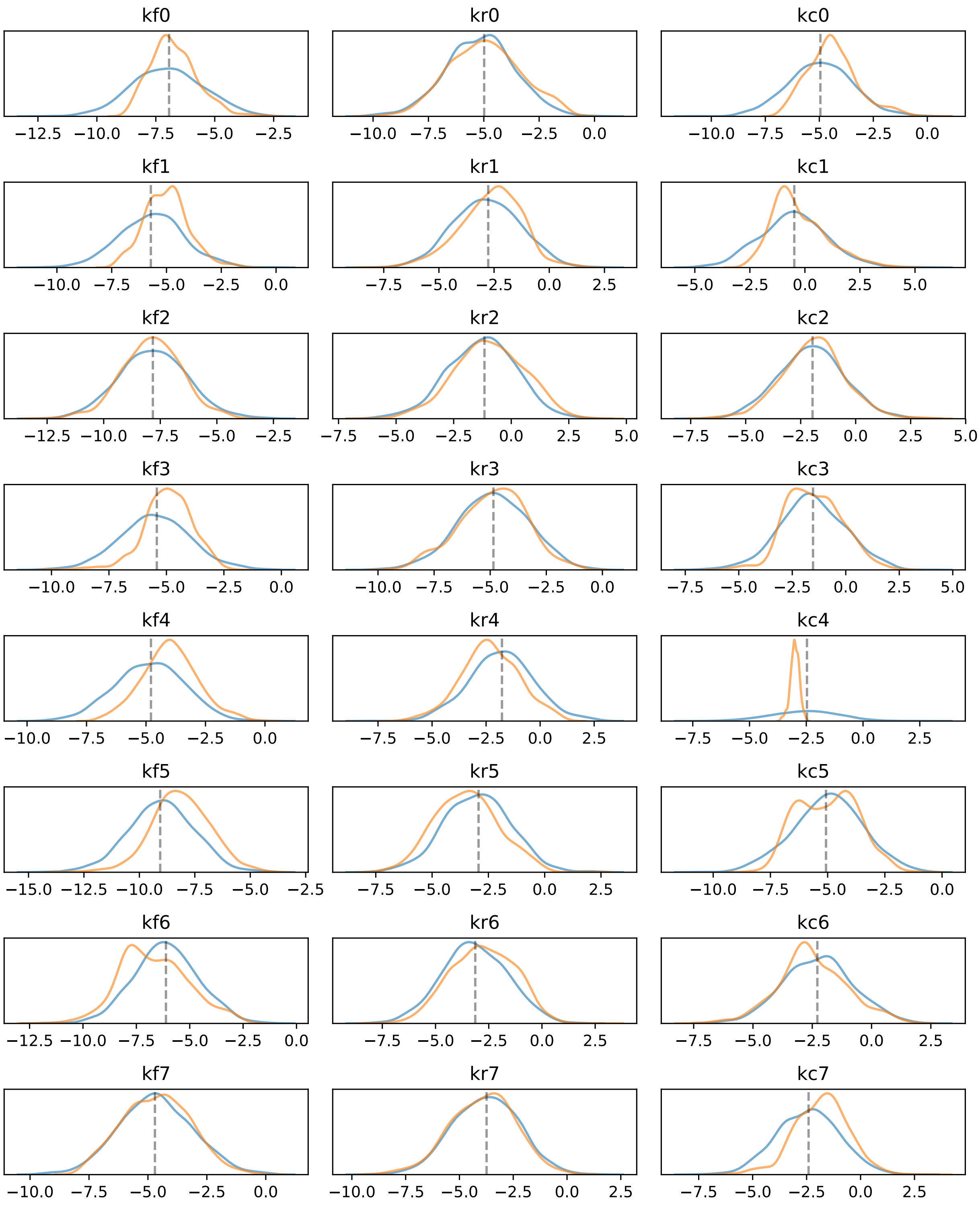

**Supplemental Figure 6A: Model parameters calibrated to an Ordinal Dataset** Parameters for aEARM were calibrated to ordinal values of tBID and cPARP abundance at every 180s interval. Prior (blue) and posterior (orange) distributions $\log_{10}$ of the value of parameter are shown.

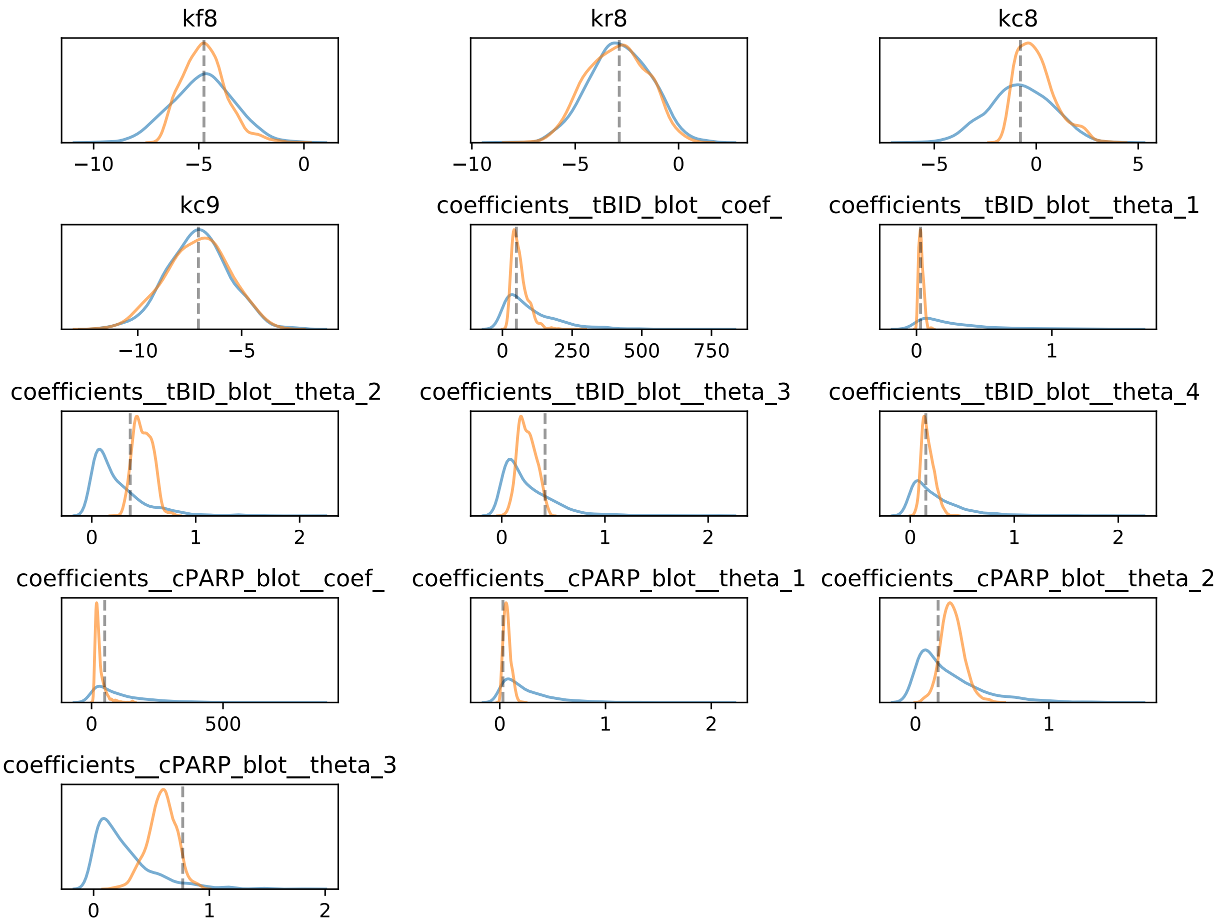

**Supplemental Figure 6B: Model parameters calibrated to an Ordinal Dataset** Parameters for aEARM were calibrated to ordinal values of tBID and cPARP abundance at every 180s interval. Prior (blue) and posterior (orange) distributions $\log_{10}$ of the value of the aEARM parameter are shown. Prior and posterior distributions of the value of measurement model coefficients are also shown (these are not log-scale).

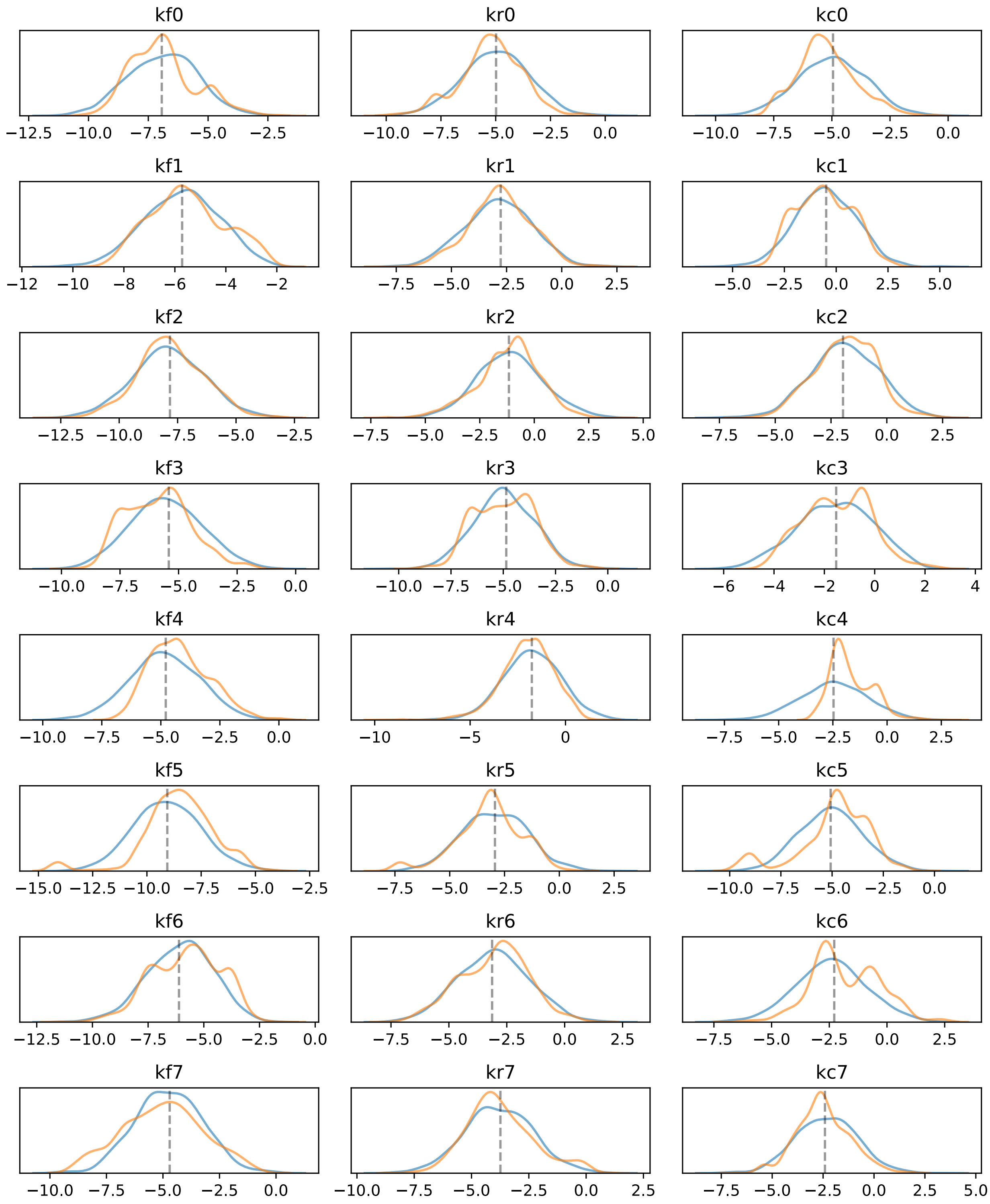

**Supplemental Figure 7A: Model parameters calibrated to an Ordinal Dataset** Parameters for aEARM were calibrated to ordinal values of tBID and cPARP abundance at every 300s interval. Prior (blue) and posterior (orange) distributions $\log_{10}$ of the value of parameter are shown.

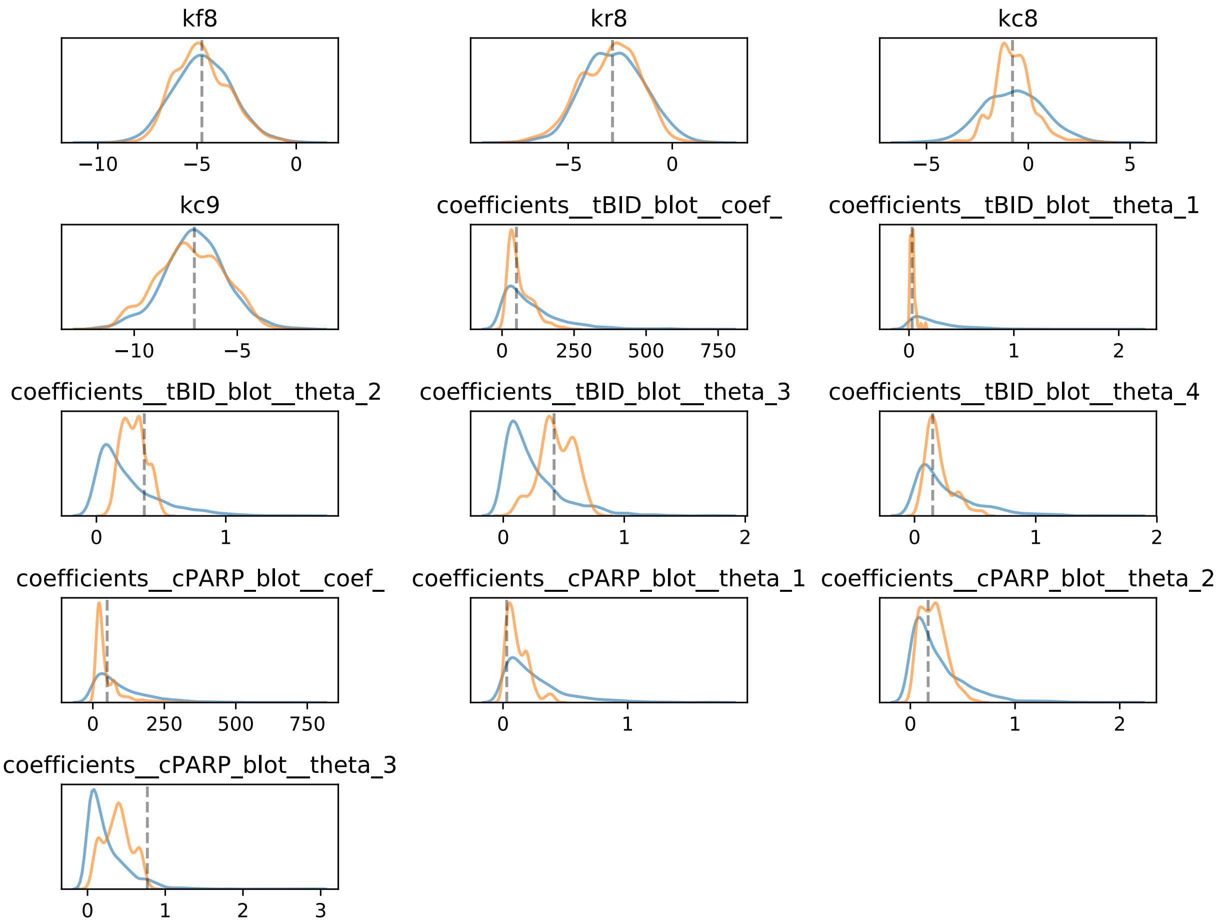

**Supplemental Figure 7B: Model parameters calibrated to an Ordinal Dataset** Parameters for aEARM were calibrated to ordinal values of tBID and cPARP abundance at every 300s interval. Prior (blue) and posterior (orange) distributions $\log_{10}$ of the value of the aEARM parameter are shown. Prior and posterior distributions of the value of measurement model coefficients are also shown (these are not log-scale).

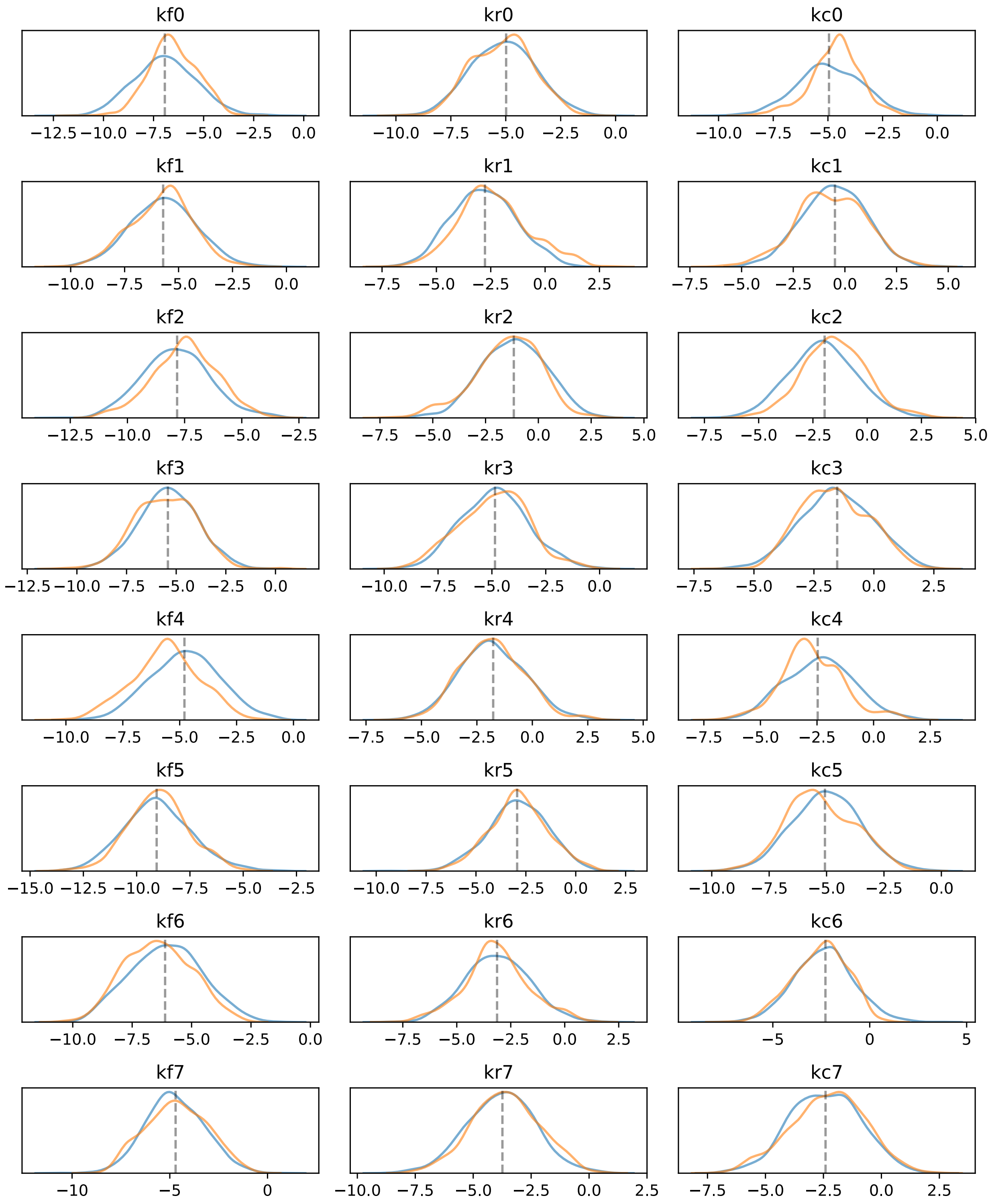

**Supplemental Figure 7A: Model parameters calibrated to an Ordinal Dataset** Parameters for aEARM were calibrated to ordinal values of tBID and cPARP abundance at every 1500s interval. Prior (blue) and posterior (orange) distributions $\log_{10}$ of the value of parameter are shown.

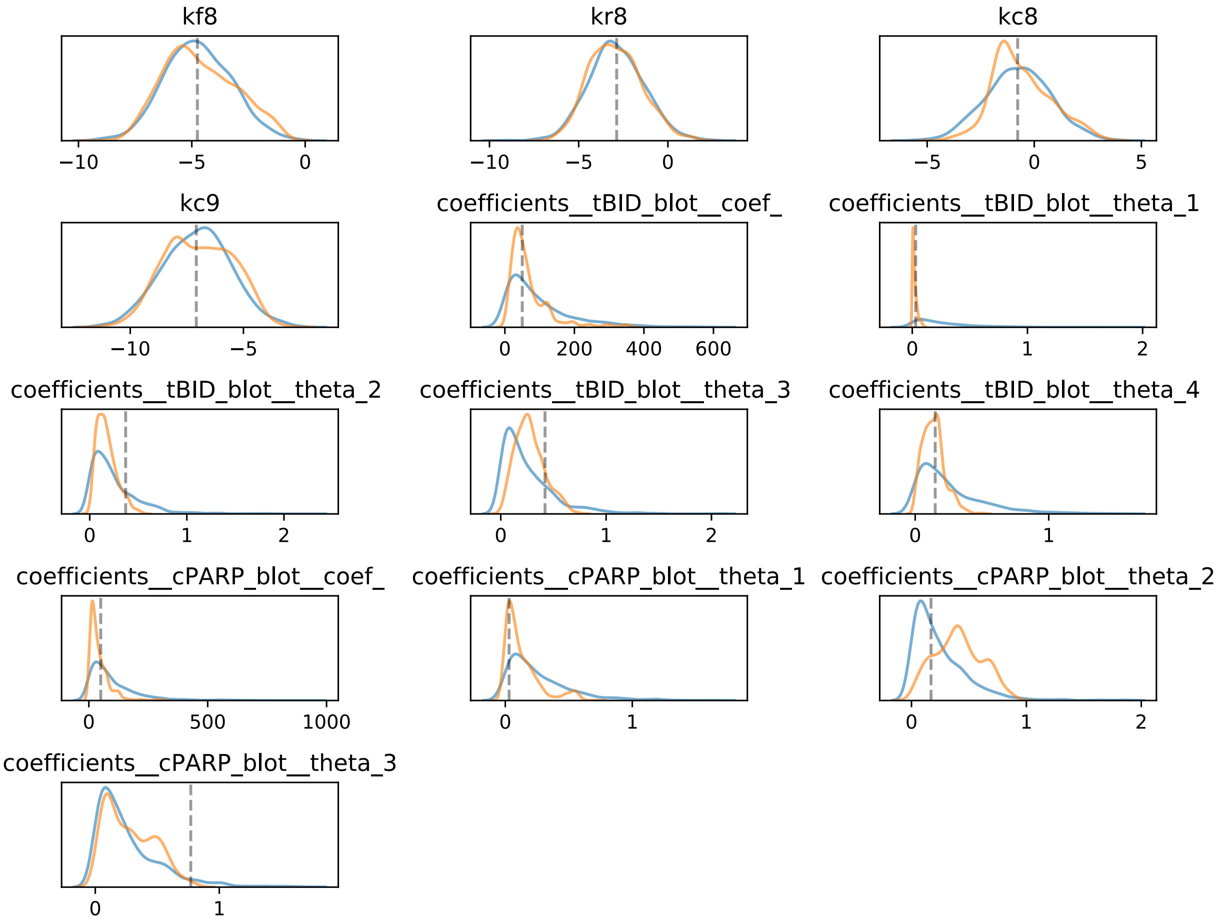

**Supplemental Figure 8B: Model parameters calibrated to an Ordinal Dataset** Parameters for aEARM were calibrated to ordinal values of tBID and cPARP abundance at every 1500s interval. Prior (blue) and posterior (orange) distributions $\log_{10}$ of the value of the aEARM parameter are shown. Prior and posterior distributions of the value of measurement model coefficients are also shown (these are not log-scale).

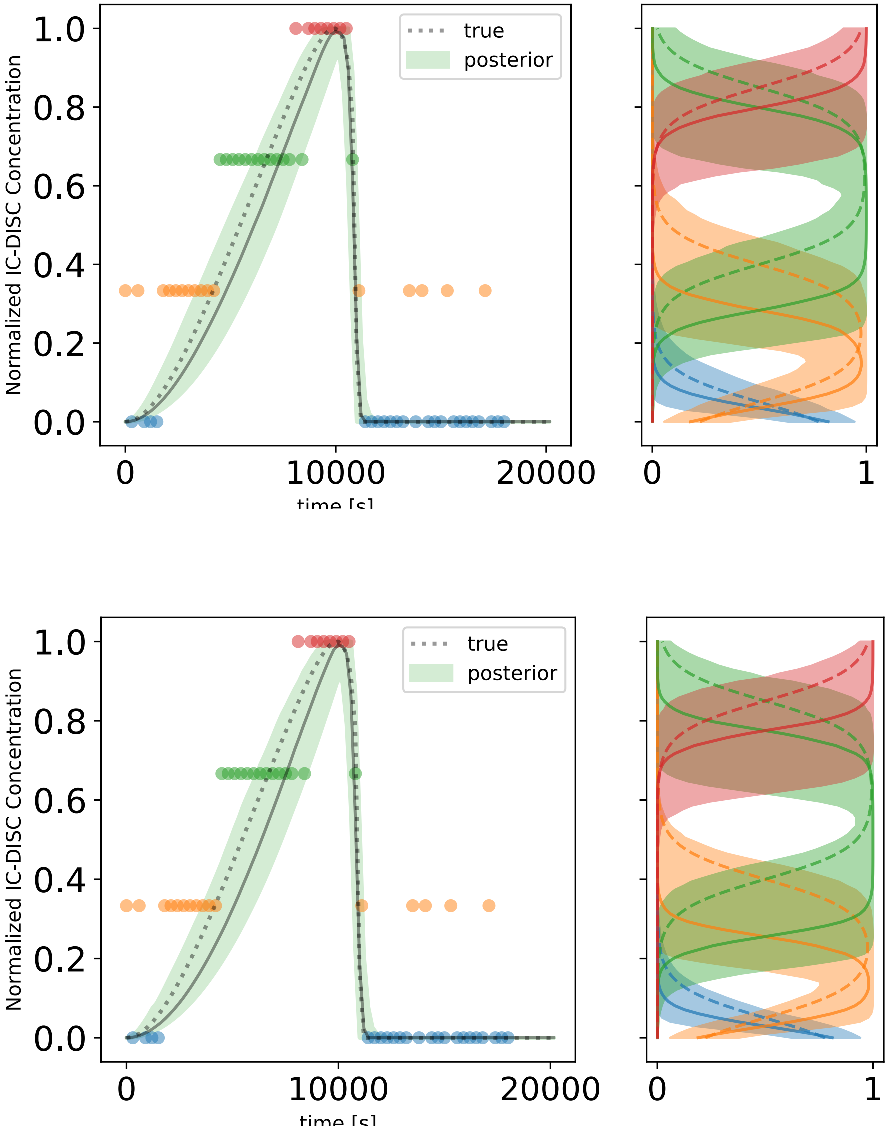

**Supplemental Figure 9. Predicted Initiator caspase and DISC colocalization dynamics of aEARM trained to ordinal and mixed ordinal/nominal datasets.**

The 95% credible region (shaded region) of posterior predictions of IC-DISC dynamics of aEARM calibrated to ordinal IC-DISC data in Figure 4C, A.) and a mixed dataset containing the nominal data in Figure 4A the ordinal IC-DISC data, C.). The median predictions (solid-line) and true (dotted line) are also plotted. The adjacent panels B.) and D.) give the 95% credible region of posterior predictions (shaded regions) for the probability of class membership (x-axis) as a function of aEARM-simulated normalized IC-DISC concentration (y-axis). The ordinal categories are color coded and plotted in ascending order.

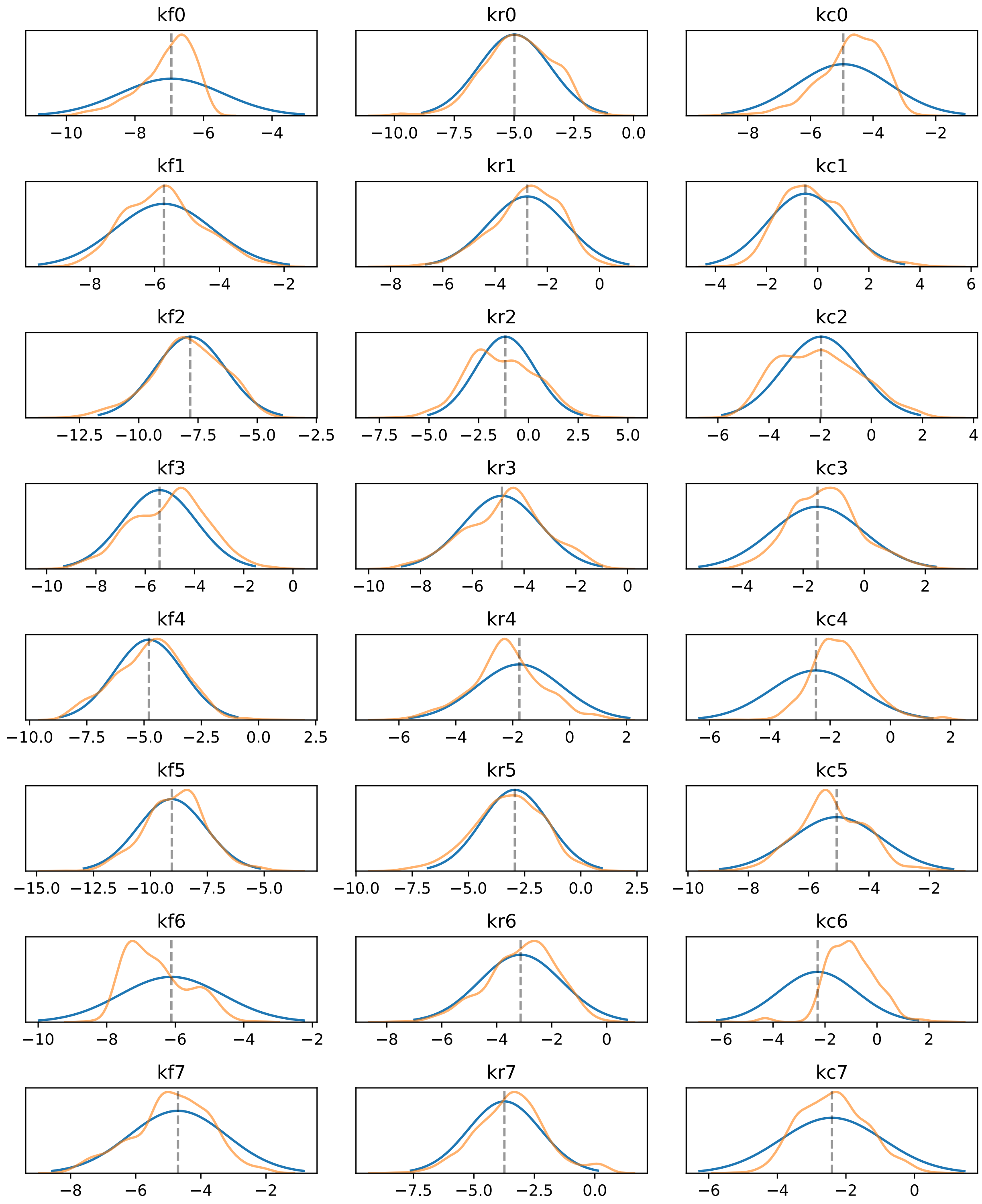

**Supplemental Figure 10A: Model parameters calibrated to a Cell Death Dataset** Parameters for aEARM were calibrated to nominal observations of cell death vs survival. Prior (blue) and posterior (orange) distributions $\log_{10}$ of the value of parameter are shown.

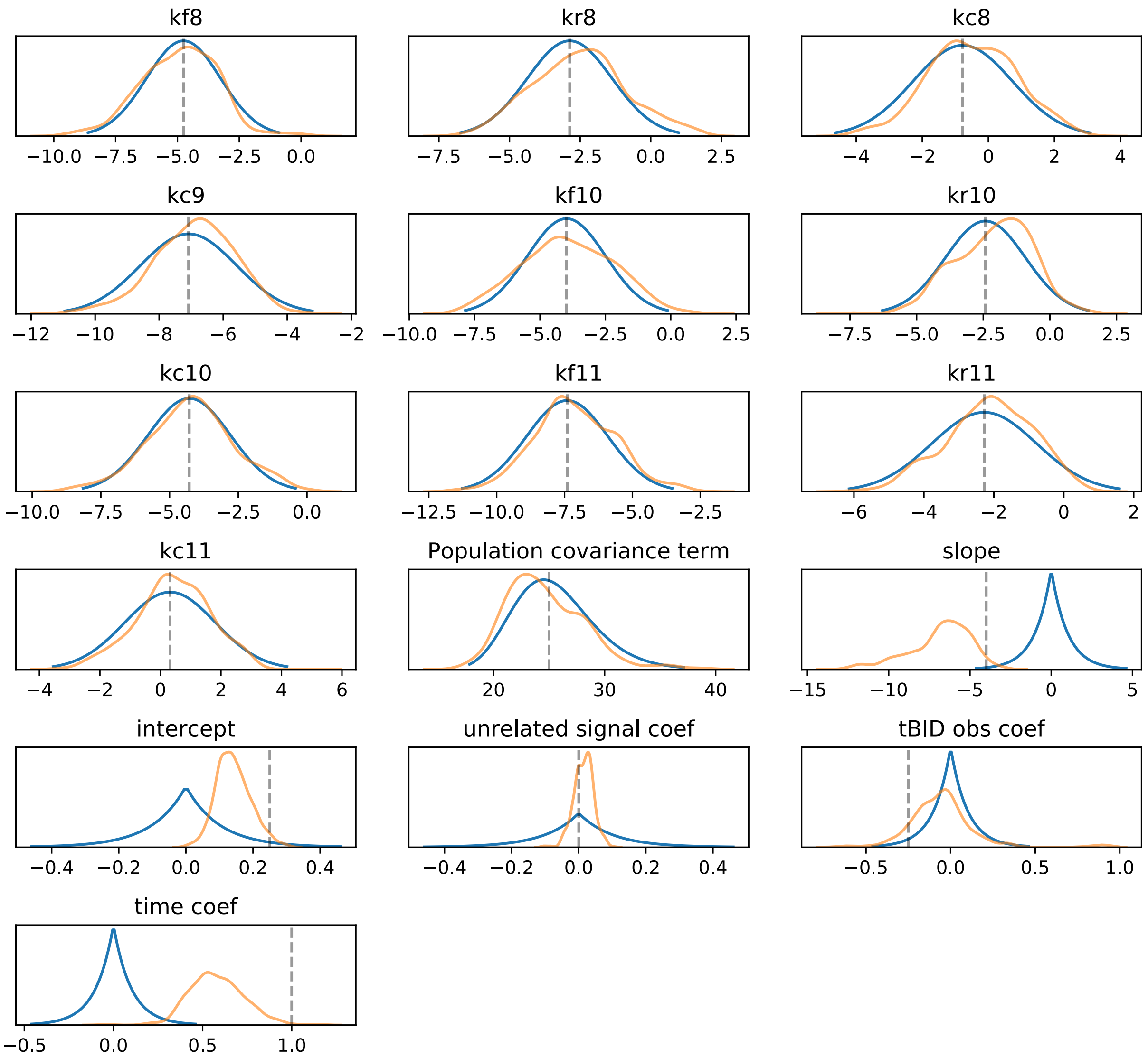

**Supplemental Figure 10B: Model parameters calibrated to a Cell Dataset** Parameters for aEARM were calibrated to nominal observations of cell death vs survival. Prior (blue) and posterior (orange) distributions $log_{10}$ of the value of the aEARM parameter are shown. Prior and posterior distributions of the value of measurement model coefficients are also shown (these are not log-scale).

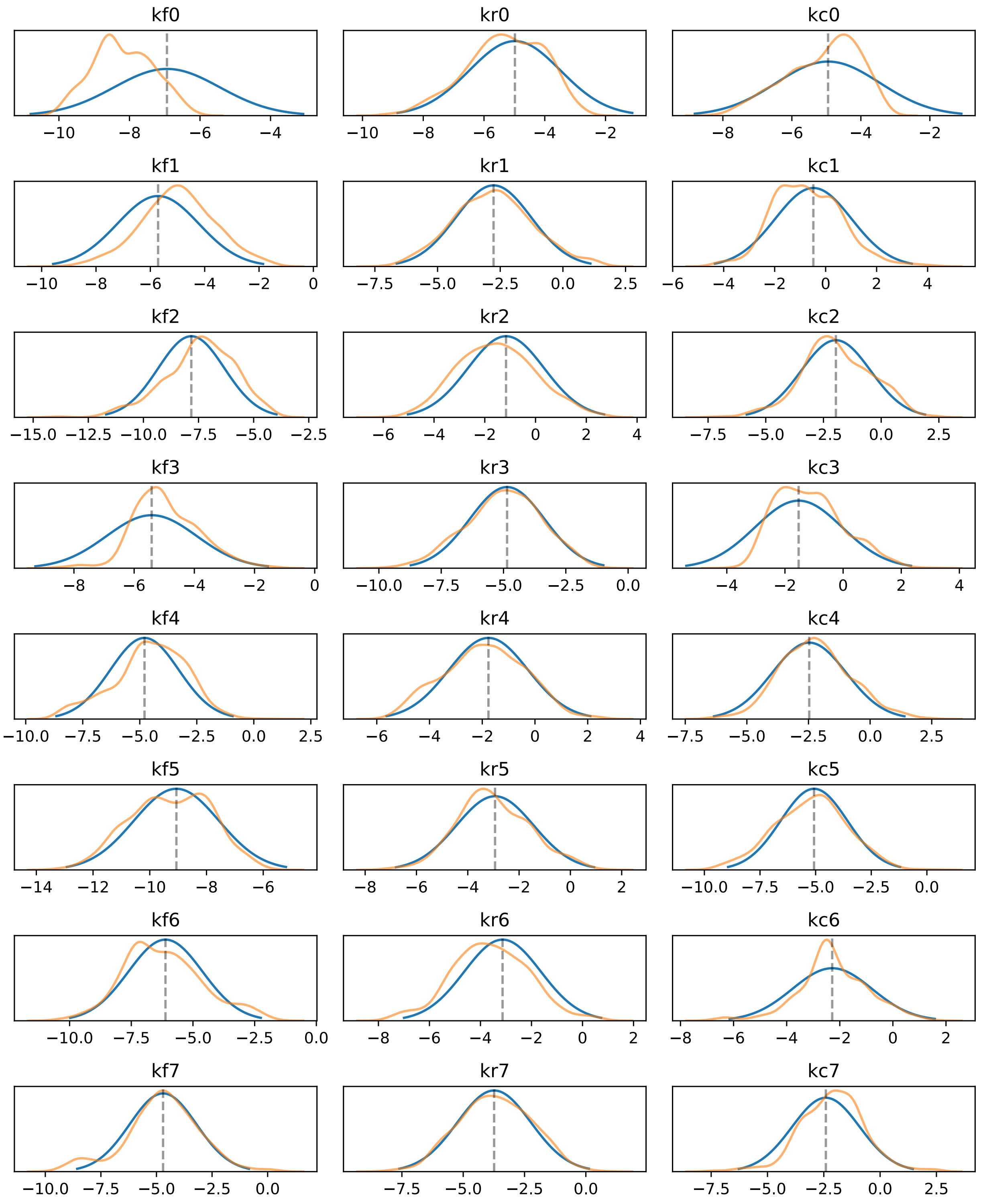

**Supplemental Figure 11A: Model parameters calibrated to Ordinal IC-DISC Dataset**
Parameters for aEARM were calibrated to ordinal values of IC-DISC abundance at every 300s interval. Prior (blue) and posterior (orange) distributions $\log_{10}$ of the value of parameter are shown.

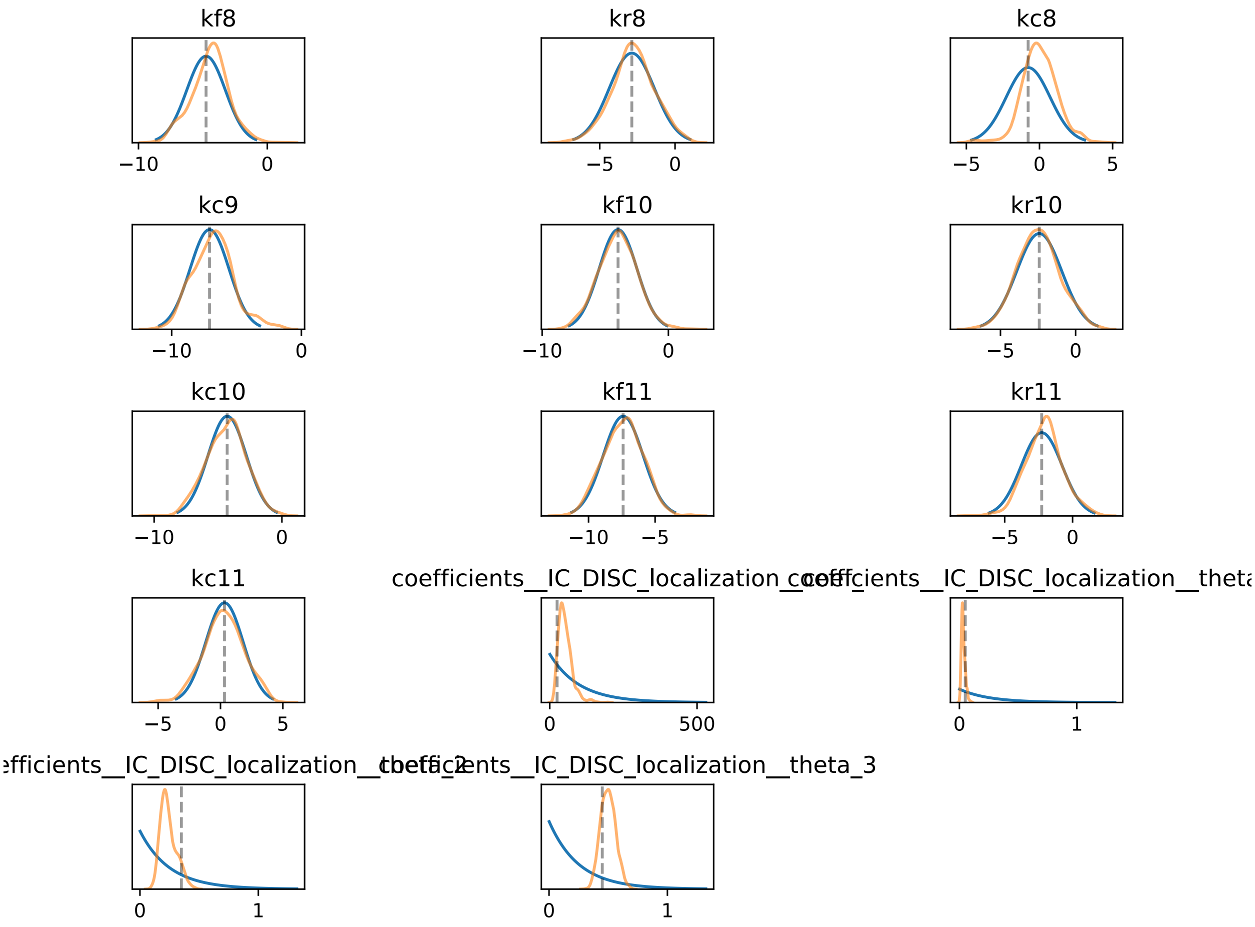

**Supplemental Figure 11B: Model parameters calibrated to Ordinal IC-DISC Dataset**
Parameters for aEARM were calibrated to ordinal values of IC-DISC abundance at every 300s interval. Prior (blue) and posterior (orange) distributions $\log_{10}$ of the value of parameter are shown. Prior and posterior distributions of the value of measurement model coefficients are also shown (these are not log-scale).

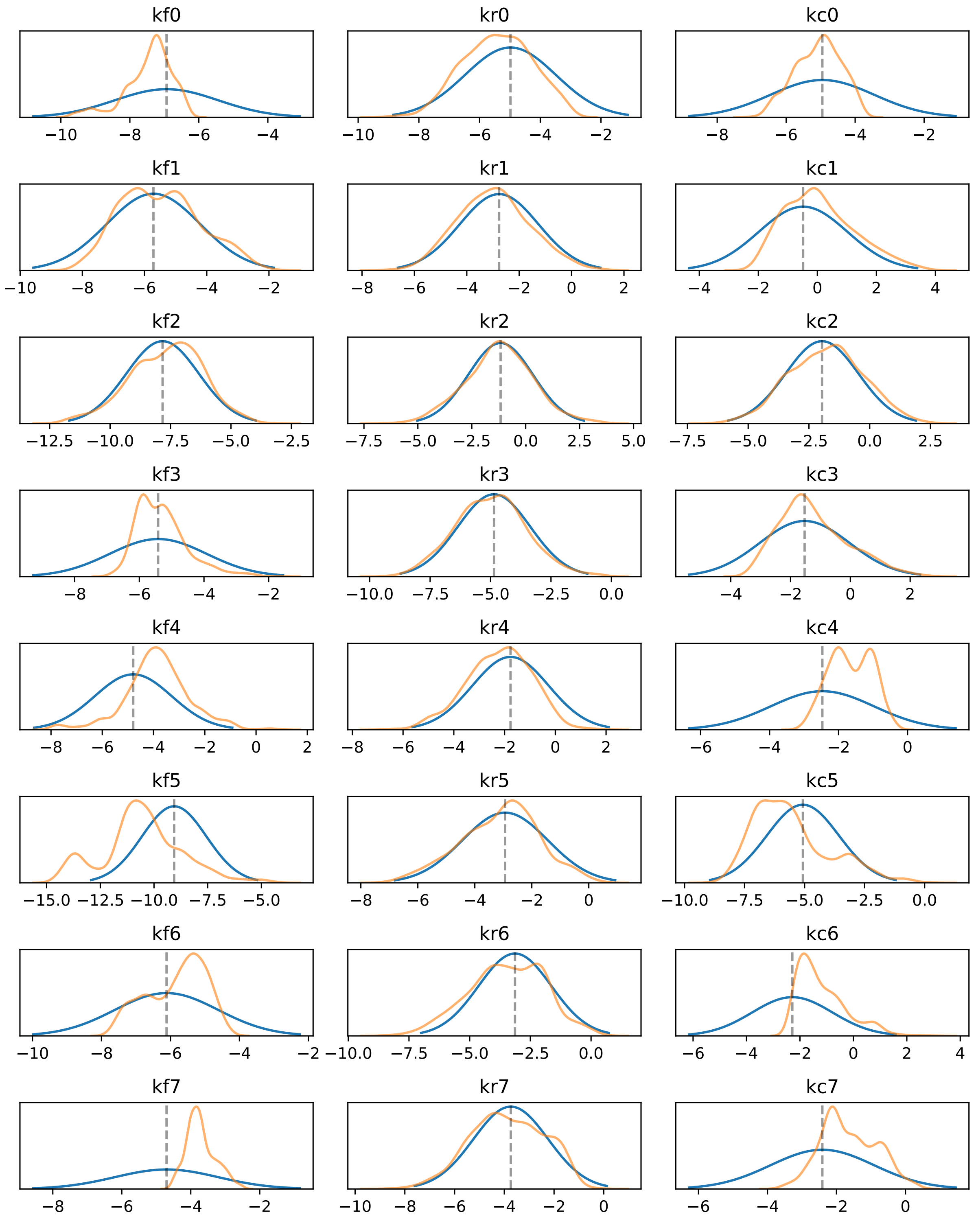

**Supplemental Figure 12A: Model parameters calibrated to Mixed Ordinal and Nominal Dataset** Parameters for aEARM were calibrated to ordinal values of IC-DISC abundance at every 300s interval and to nominal observations of cell death vs survival. Prior (blue) and posterior (orange) distributions $\log_{10}$ of the value of parameter are shown.

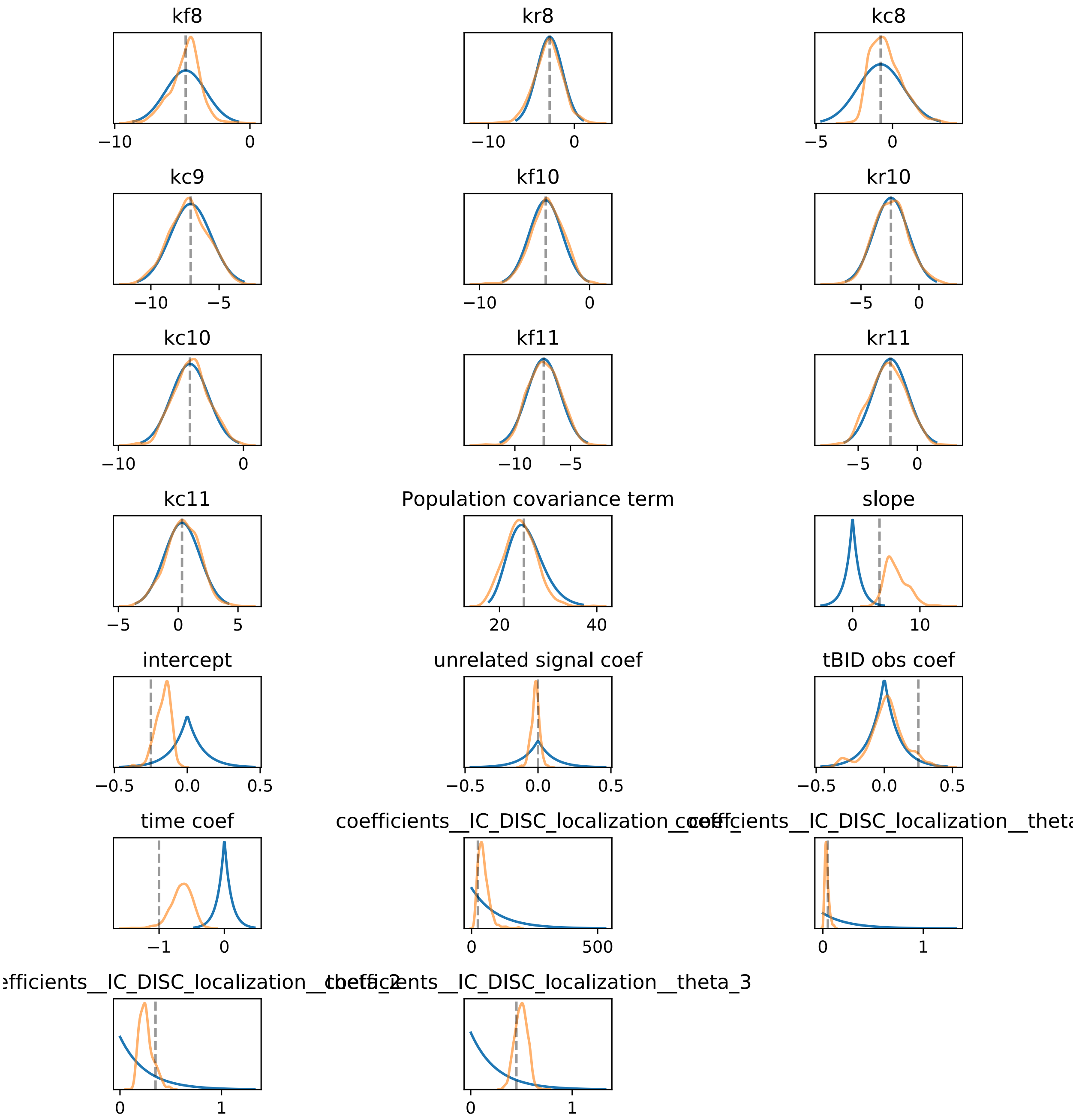

**Supplemental Figure 12B: Model parameters calibrated to Mixed Ordinal and Nominal Dataset** Parameters for aEARM were calibrated to ordinal values of IC-DISC abundance at every 300s interval and to nominal observations of cell death vs survival. Prior (blue) and posterior (orange) distributions $\log_{10}$ of the value of parameter are shown. Prior and posterior distributions of the value of measurement model coefficients are also shown (these are not log-scale).

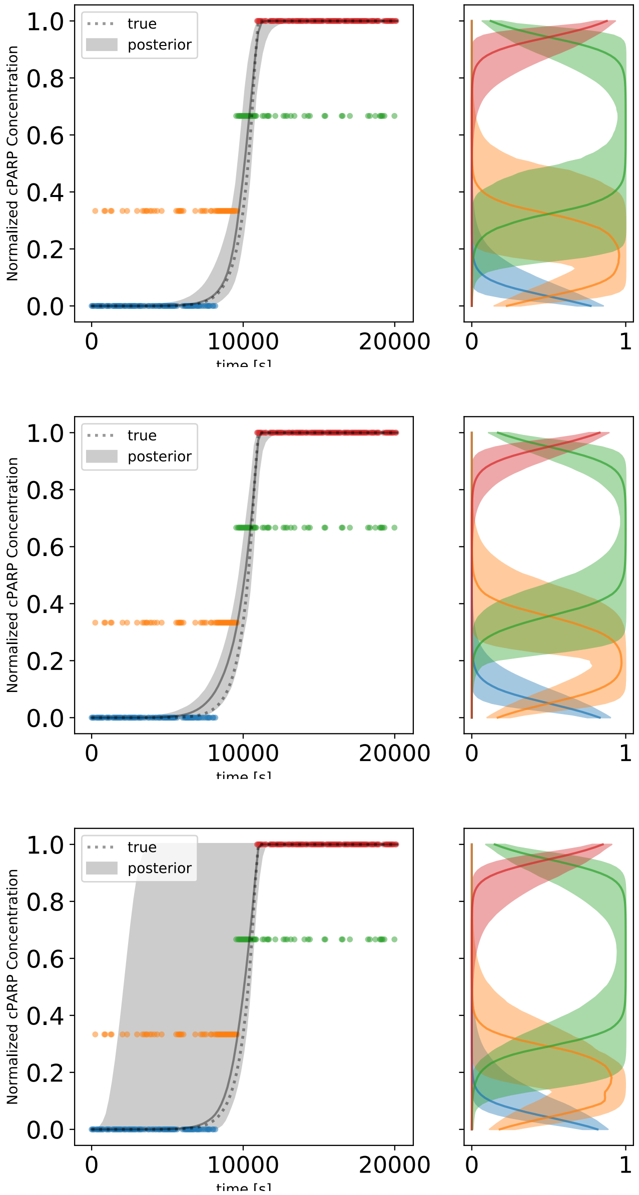

**Supplemental Figure 13: Predicted PARP cleavage dynamics of aEARM trained to ordinal data using different measurement model priors.**

The 95% credible region of posterior predictions (shaded region) of cPARP dynamics of aEARM calibrated to ordinal values of tBID and cPARP at every 60s interval. Uniform, Cauchy (scale=0.05) and Cauchy (scale=0.005) prior distributions for the parameterizations for the measurement model, respectively. In each, the median prediction (solid line) and true (dotted line) cPARP dynamic sare also shown. The adjacent panels give the 95% credible region of posterior predictions of the probability of class membership (x-axis) as a function of normalized cPARP concentration (y-axis).

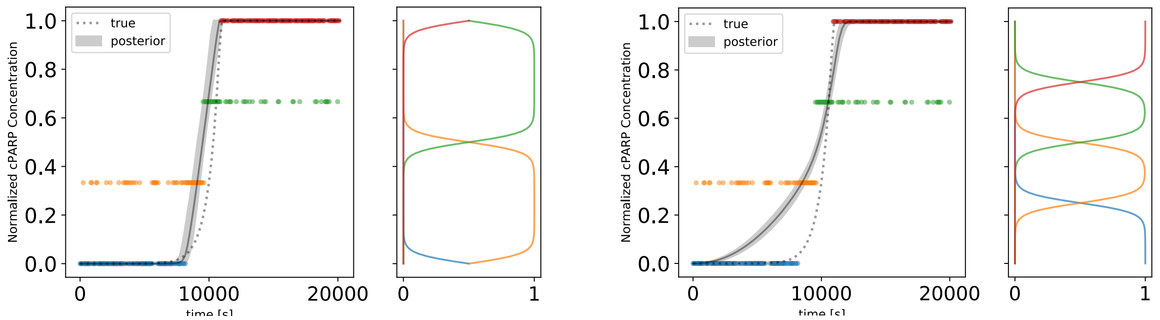

**Supplemental Figure 14: Predicted PARP cleavage dynamics of aEARM trained to ordinal data using different *ad hoc* parameterizations of the measurement model.** The 95% credible region of posterior predictions (shaded region) of cPARP dynamics for aEARM calibrated to ordinal measurements of tBID and cPARP at every 60s interval. Two fixed parameterizations for the measurement model. The adjacent panels plot the measurement models predicted probability of class membership (x-axis) as a function of normalized cPARP concentration (y-axis). The adjacent panels give the 95% credible region of posterior predictions of the probability of class membership (x-axis) as a function of normalized cPARP concentration (y-axis).

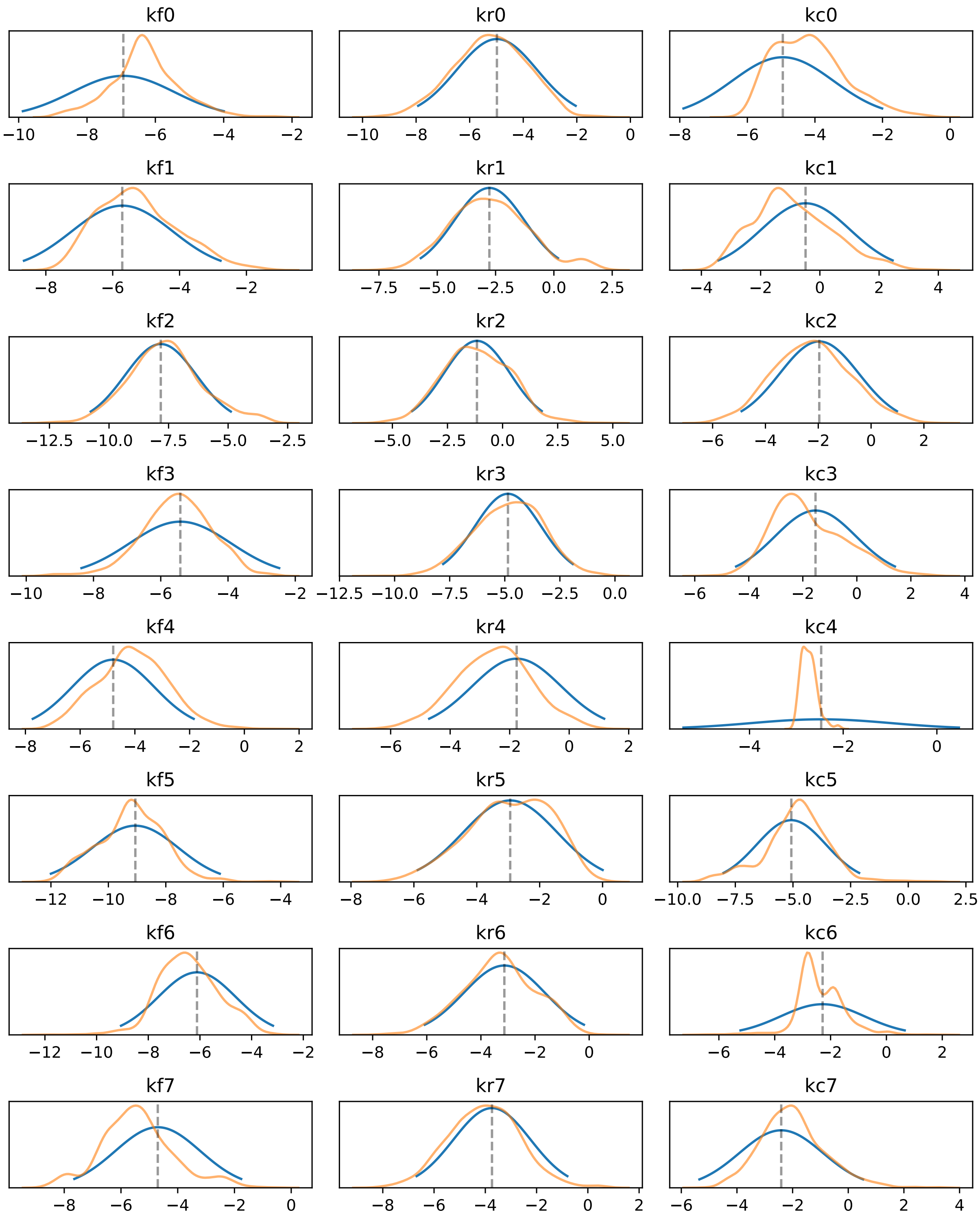

**Supplemental Figure 15A: Model parameters calibrated to Ordinal Dataset using Uniform priors on Measurement Model Parameters** Parameters for aEARM were calibrated to ordinal values of tBID and cPARP abundance at every 60s interval. Prior (blue) and posterior (orange) distributions $\log_{10}$ of the value of parameter are shown. Uniform priors were placed on the measurement model parameters (these are not log-scale).

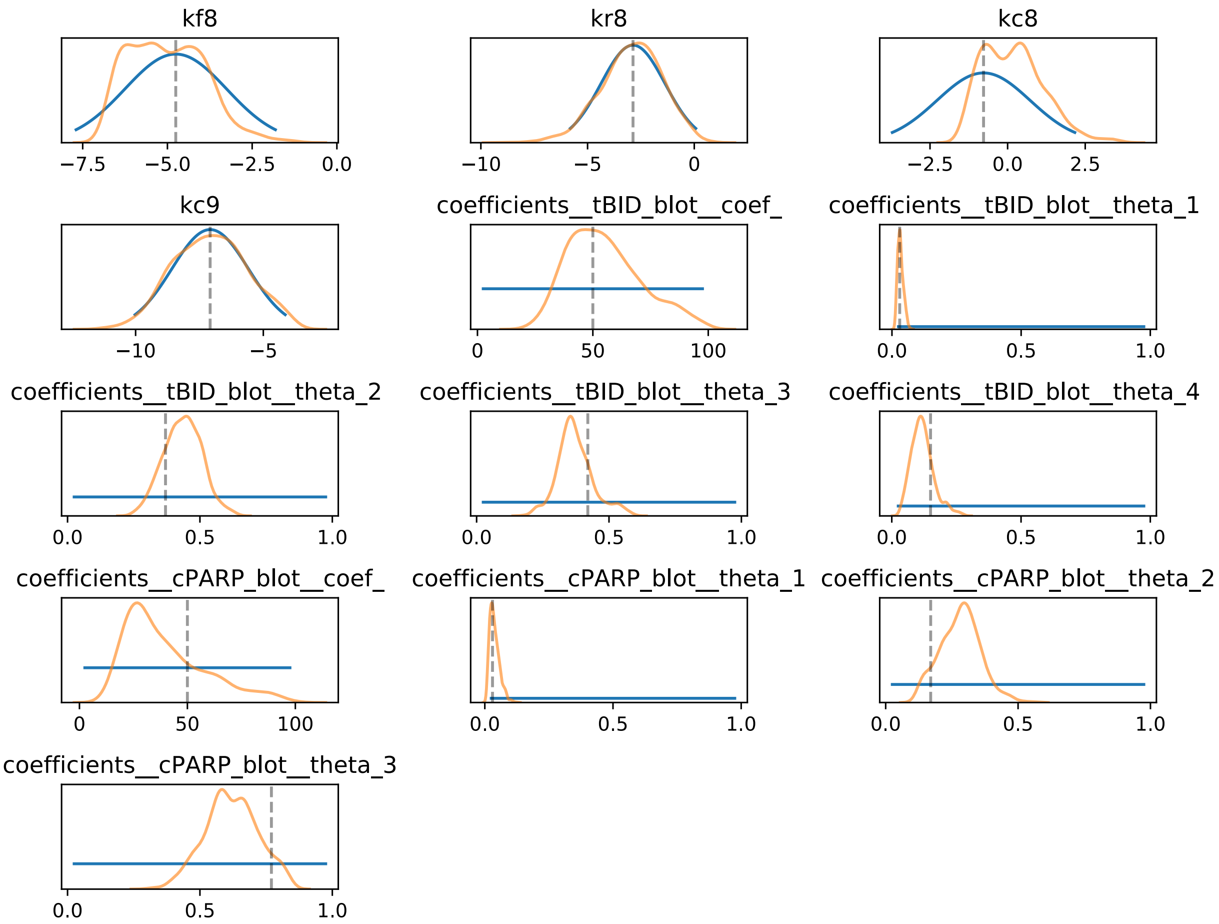

**Supplemental Figure 15B: Model parameters calibrated to Ordinal Dataset using Uniform priors on Measurement Model Parameters** Parameters for aEARM were calibrated to ordinal values of tBID and cPARP abundance at every 60s interval. Prior (blue) and posterior (orange) distributions $log_{10}$ of the value of parameter are shown. Uniform priors were placed on the measurement model parameters (these are not log-scale).

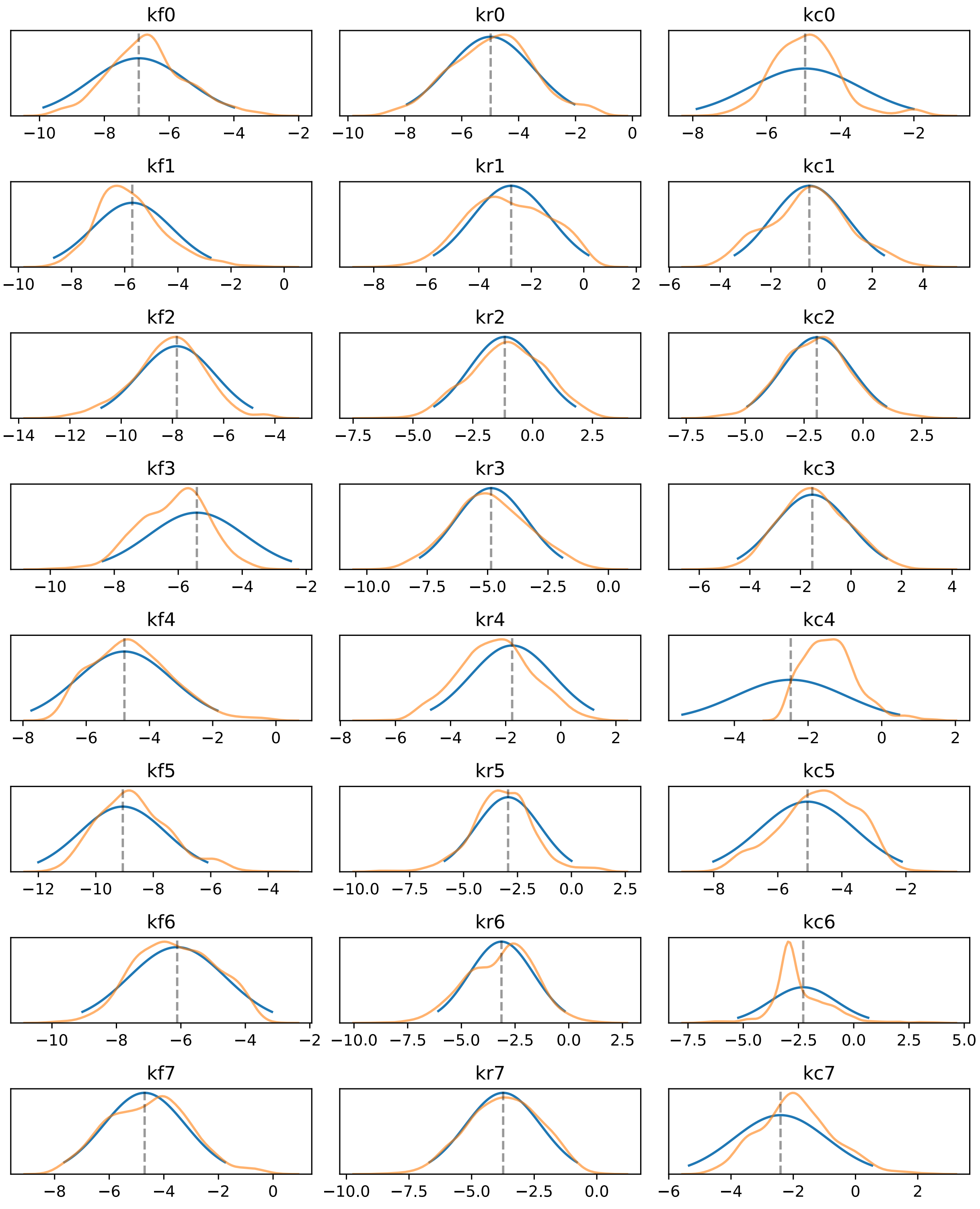

**Supplemental Figure 16A: Model parameters calibrated to Ordinal Dataset using Cauchy priors on Measurement Model Parameters.** Parameters for aEARM were calibrated to ordinal values of tBID and cPARP abundance at every 60s interval. Prior (blue) and posterior (orange) distributions $\log_{10}$ of the value of parameter are shown. Cauchy priors with a scale term of 0.05 were placed on the measurement model parameters (these are not log-scale).

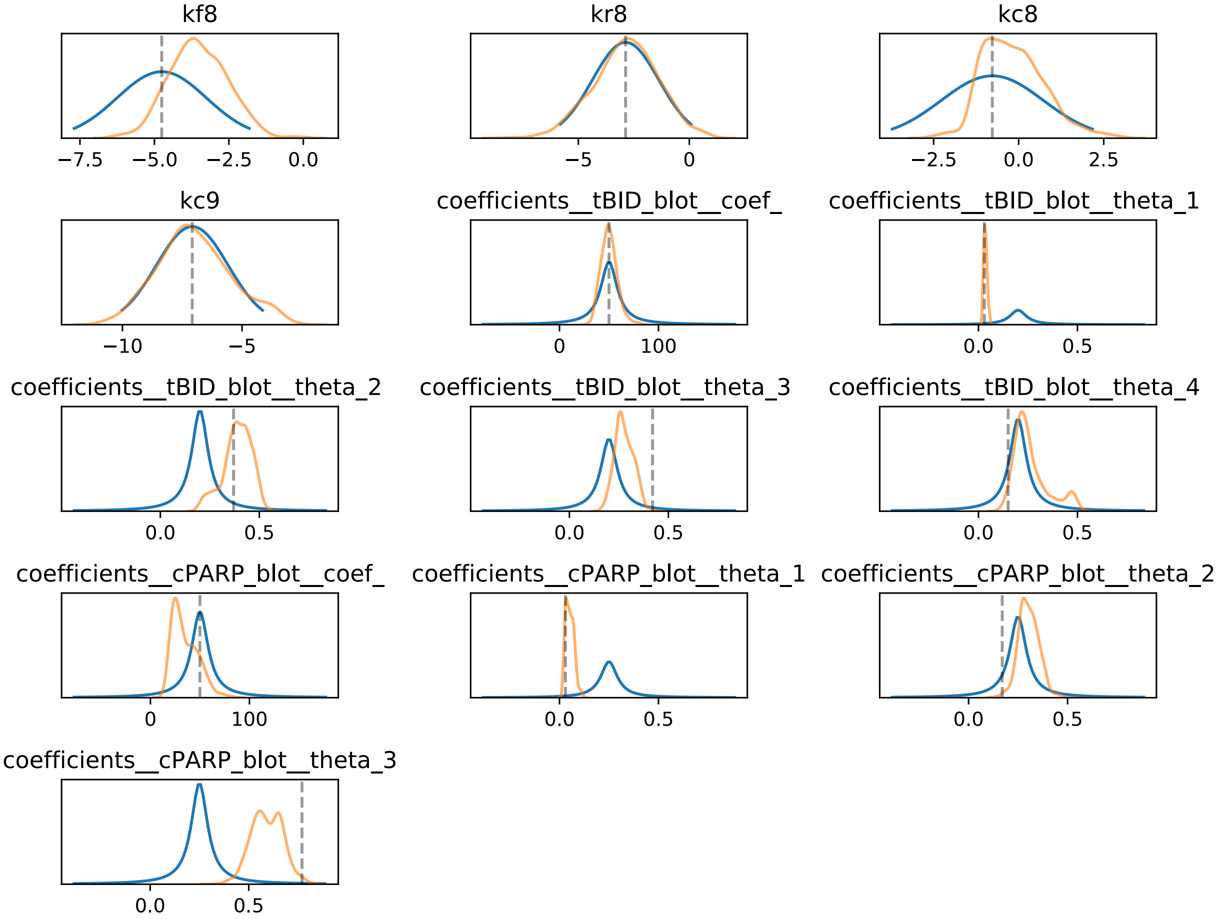

**Supplemental Figure 16B: Model parameters calibrated to Ordinal Dataset using Cauchy priors on Measurement Model Parameters.** Parameters for aEARM were calibrated to ordinal values of tBID and cPARP abundance at every 60s interval. Prior (blue) and posterior (orange) distributions $\log_{10}$ of the value of parameter are shown. Cauchy priors with a scale term of 0.05 were placed on the measurement model parameters (these are not log-scale)

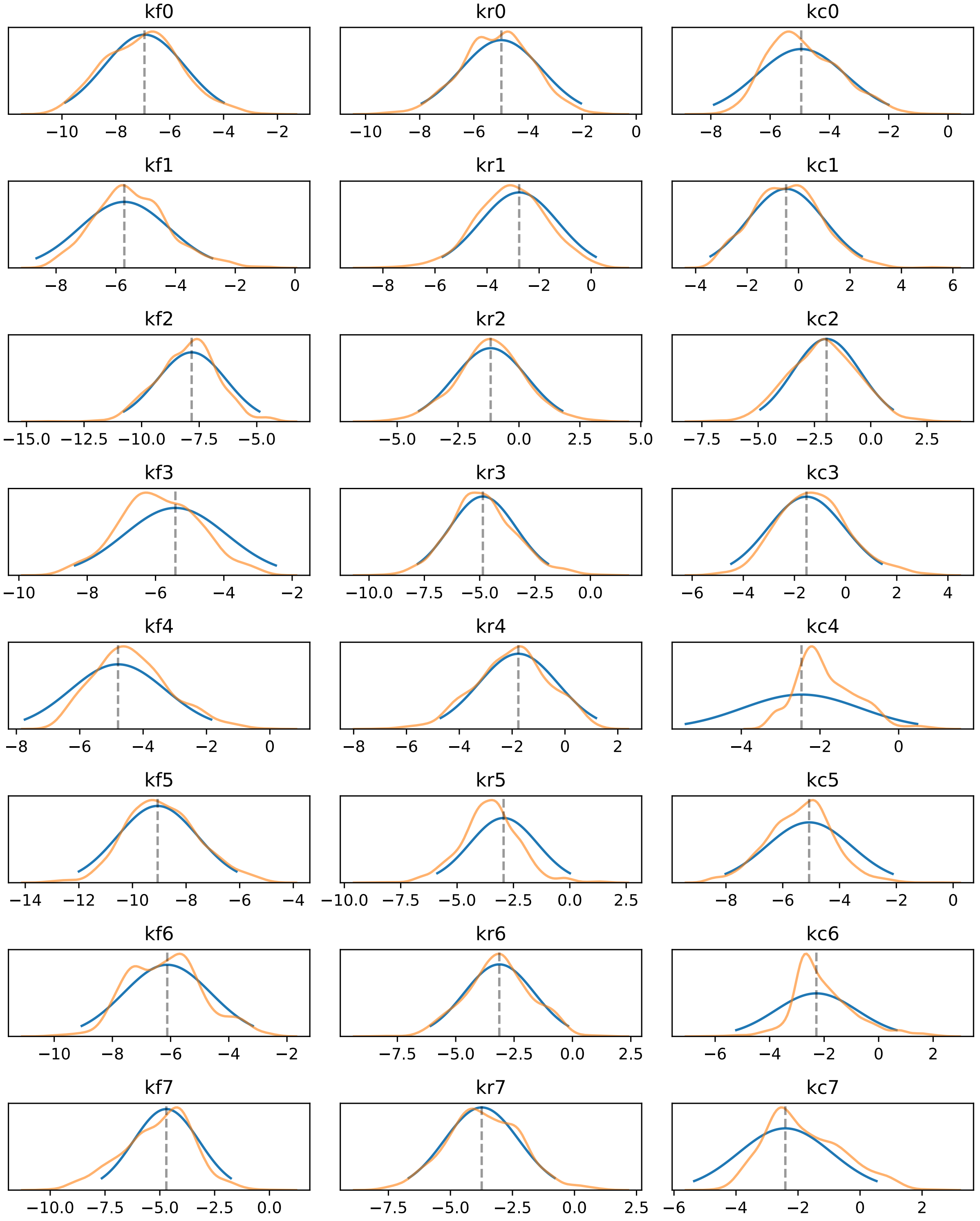

**Supplemental Figure 17A: Model parameters calibrated to Ordinal Dataset using Cauchy priors on Measurement Model Parameters.** Parameters for aEARM were calibrated to ordinal values of tBID and cPARP abundance at every 60s interval. Prior (blue) and posterior (orange) distributions $log_{10}$ of the value of parameter are shown. Cauchy priors with a scale term of

0.005 were placed on the measurement model parameters (these are not log-scale).

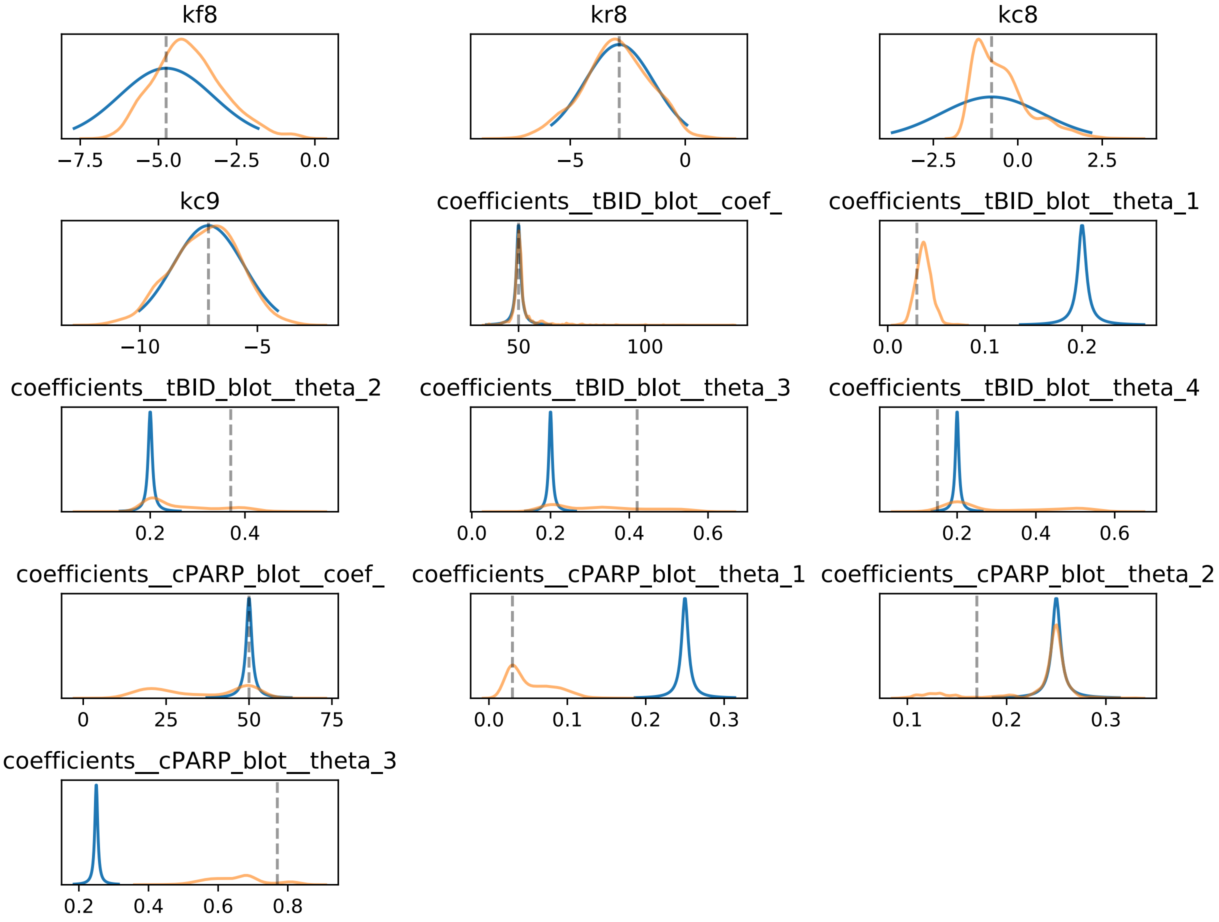

**Supplemental Figure 17B: Model parameters calibrated to Ordinal Dataset using Cauchy priors on Measurement Model Parameters.** Parameters for aEARM were calibrated to ordinal values of tBID and cPARP abundance at every 60s interval. Prior (blue) and posterior (orange) distributions $\log_{10}$ of the value of parameter are shown. Cauchy priors with a scale term of 0.005 were placed on the measurement model parameters (these are not log-scale).

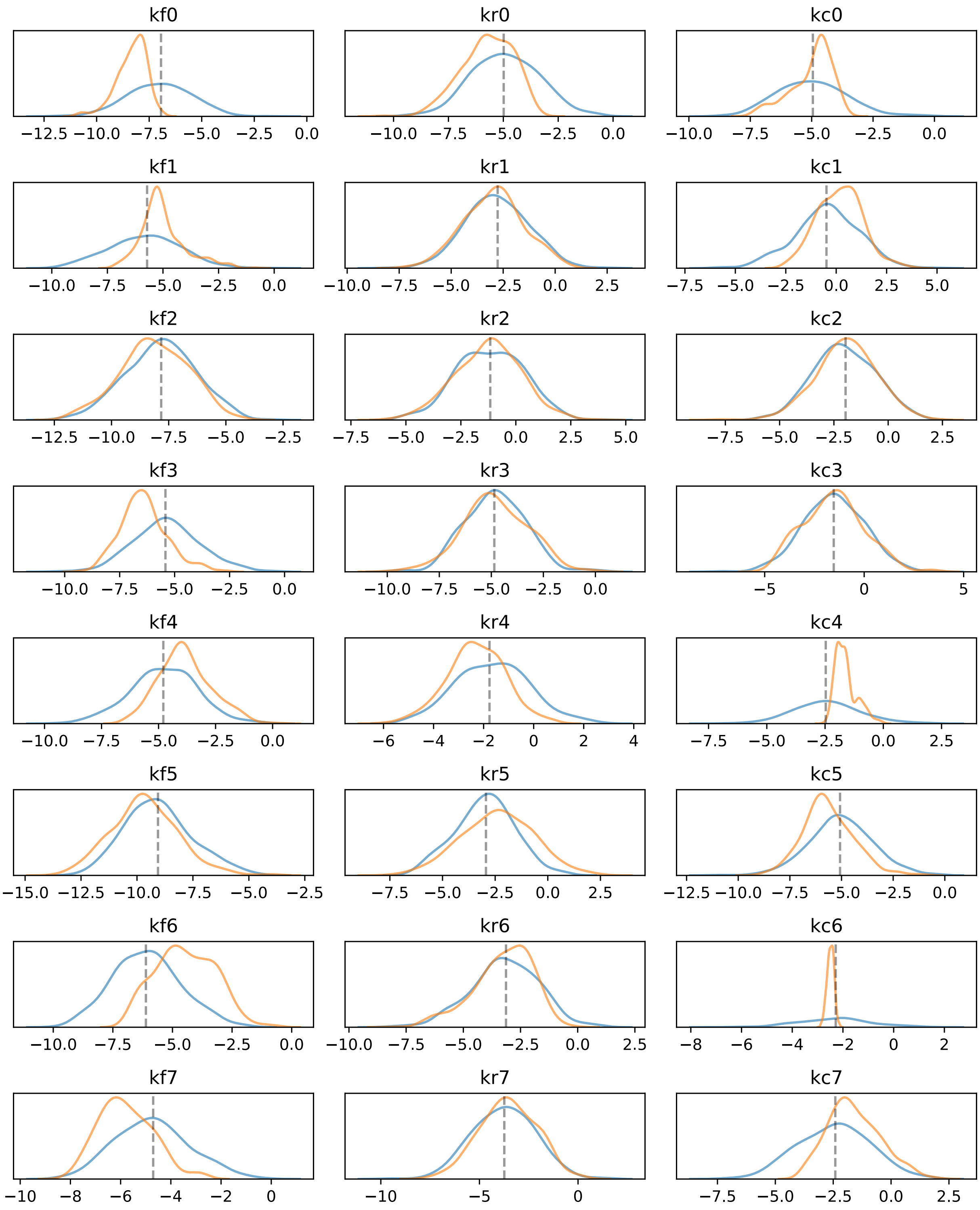

**Supplemental Figure 18A: Model parameters calibrated to Ordinal Dataset using Fixed *ad hoc* Measurement Model Parameters.** Parameters for aEARM were calibrated to ordinal values of tBID and cPARP abundance at every 60s interval. Prior (blue) and posterior (orange) distributions $\log_{10}$ of the value of parameter are shown. Fixed *ad hoc* values were applied to the measurement model parameters (see Supplemental Table 3 (Case 1))

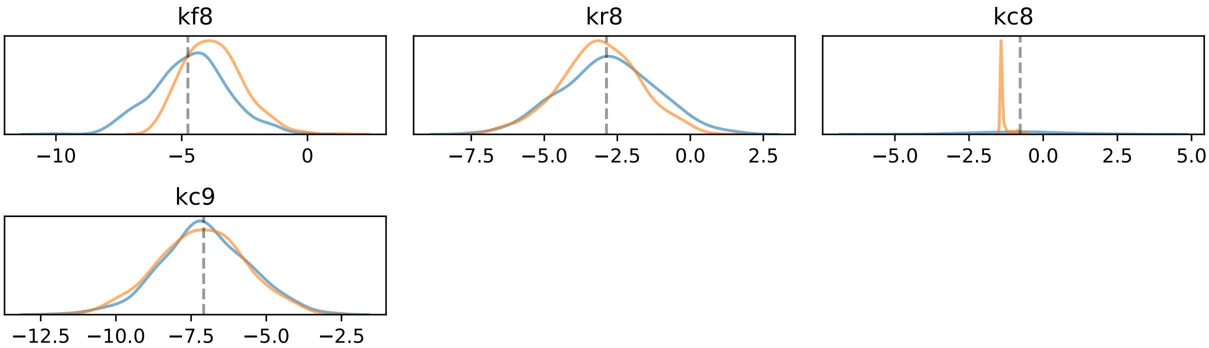

**Supplemental Figure 18B: Model parameters calibrated to Ordinal Dataset using Fixed *ad hoc* Measurement Model Parameters.** Parameters for aEARM were calibrated to ordinal values of tBID and cPARP abundance at every 60s interval. Prior (blue) and posterior (orange) distributions $\log_{10}$ of the value of parameter are shown. Fixed *ad hoc* values were applied to the measurement model parameters (see Supplemental Table 3 (Case 1))

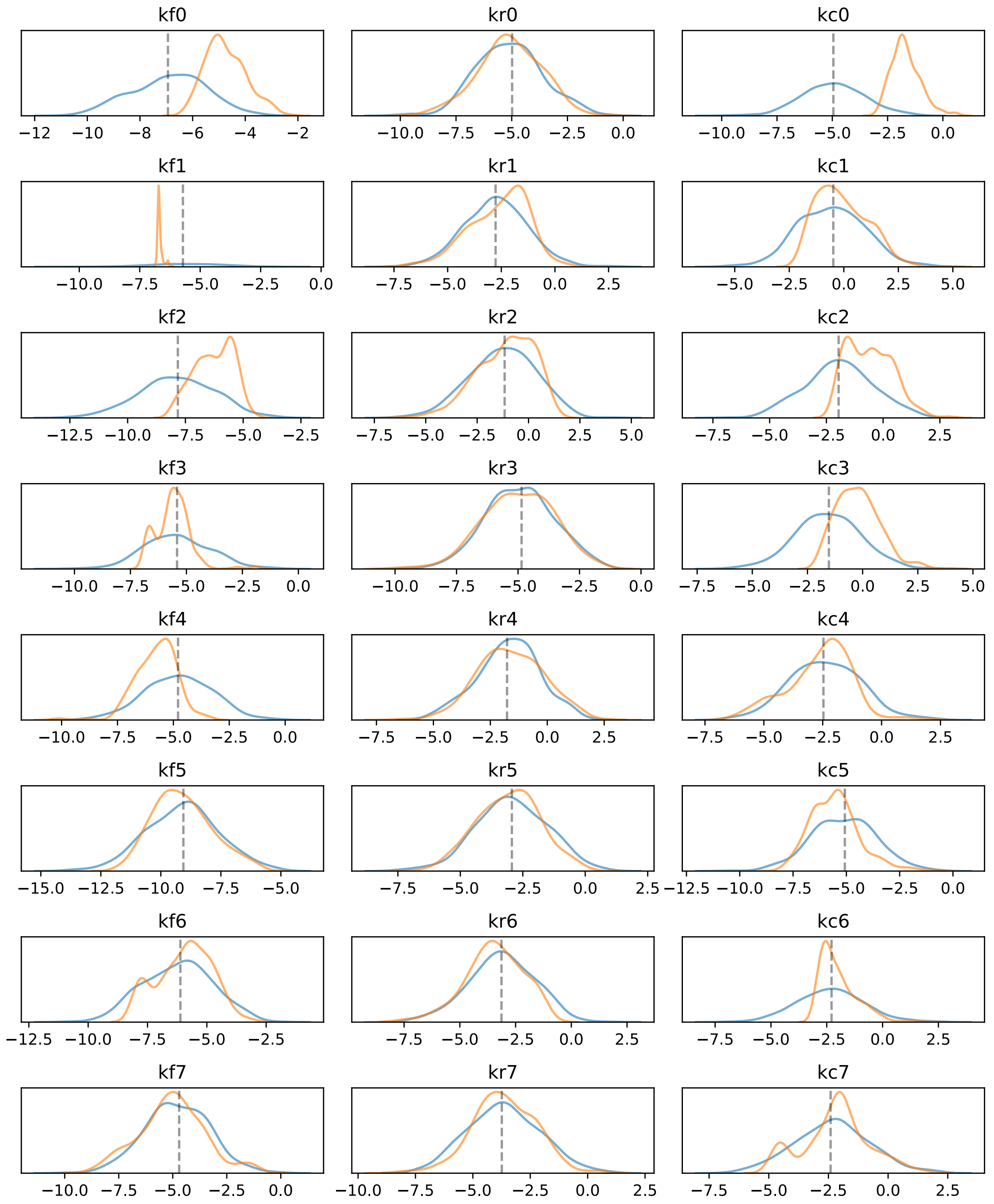

**Supplemental Figure 19A: Model parameters calibrated to Ordinal Dataset using Fixed *ad hoc* Measurement Model Parameters.** Parameters for aEARM were calibrated to ordinal values of tBID and cPARP abundance at every 60s interval. Prior (blue) and posterior (orange) distributions $\log_{10}$ of the value of parameter are shown. Fixed *ad hoc* values were applied to the measurement model parameters (see Supplemental Table 3 (Case 2))

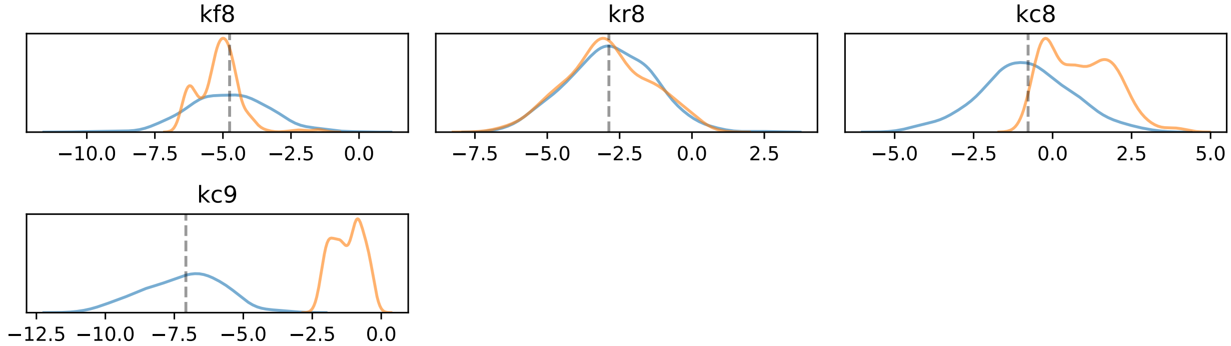

**Supplemental Figure 19B: Model parameters calibrated to Ordinal Dataset using Fixed *ad hoc* Measurement Model Parameters.** Parameters for aEARM were calibrated to ordinal values of tBID and cPARP abundance at every 60s interval. Prior (blue) and posterior (orange) distributions $\log_{10}$ of the value of parameter are shown. Fixed *ad hoc* values were applied to the measurement model parameters (see Supplemental Table 3 (Case 2))

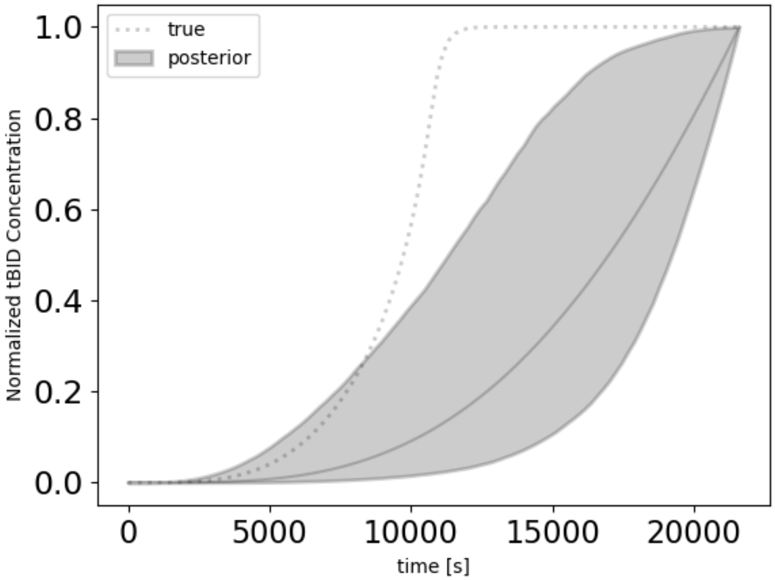

**Supplemental Figure 20: Predicted Bid truncation dynamics of aEARM trained to published fractional cell death data.** The 95% credible region of posterior predictions (shaded region) of normalized tBID dynamics for aEARM calibrated to published fractional cell death measurements by Wajant et al. (see *Methods*)[31].

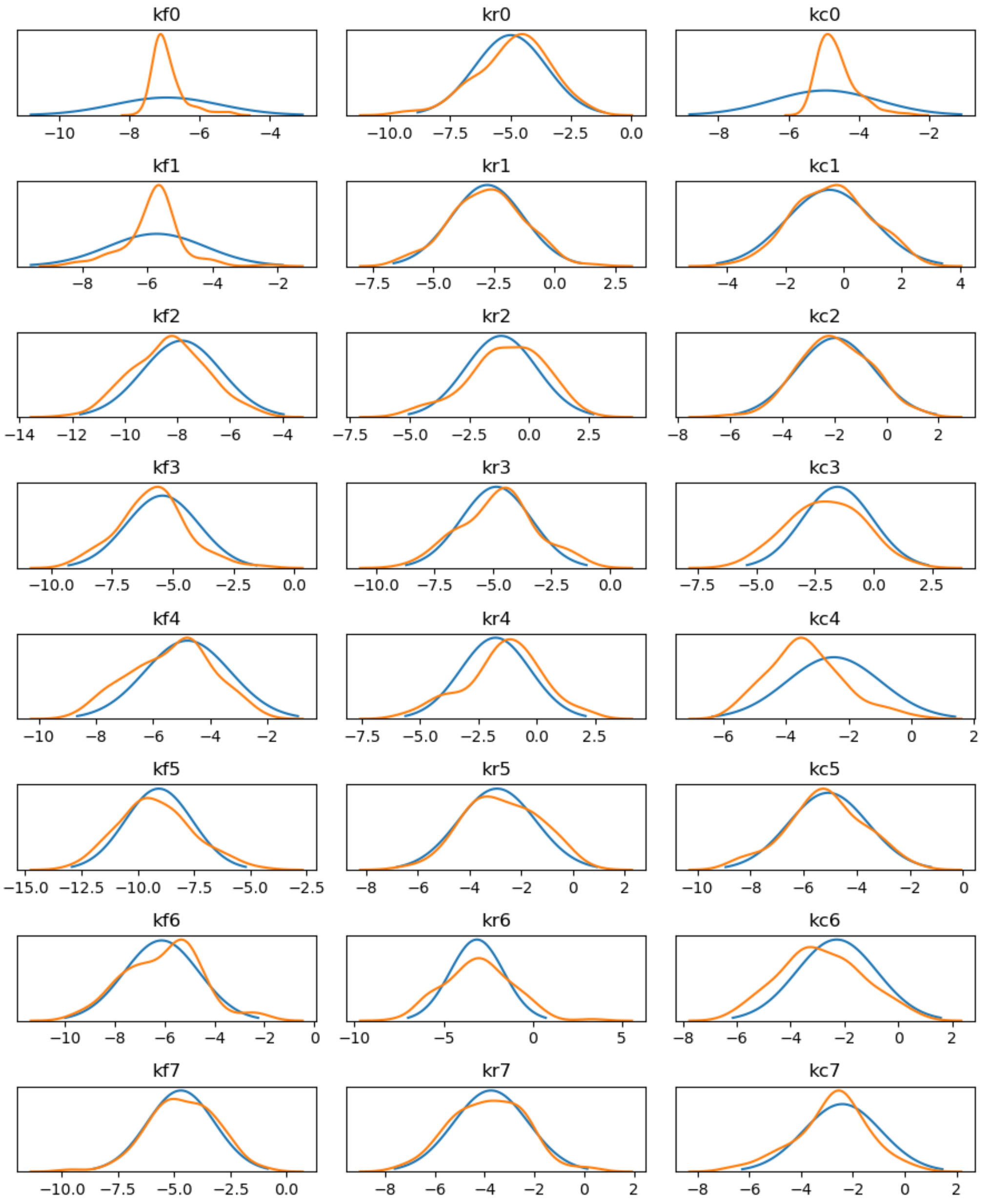

**Supplemental Figure 21A: Model parameters calibrated to published fractional cell death data.** Parameters for aEARM were calibrated to published fractional cell death measurements by Wajant et al. (see *Methods*)[31]. Prior (blue) and posterior (orange) distributions of $\log_{10}$ of the value of parameter are shown

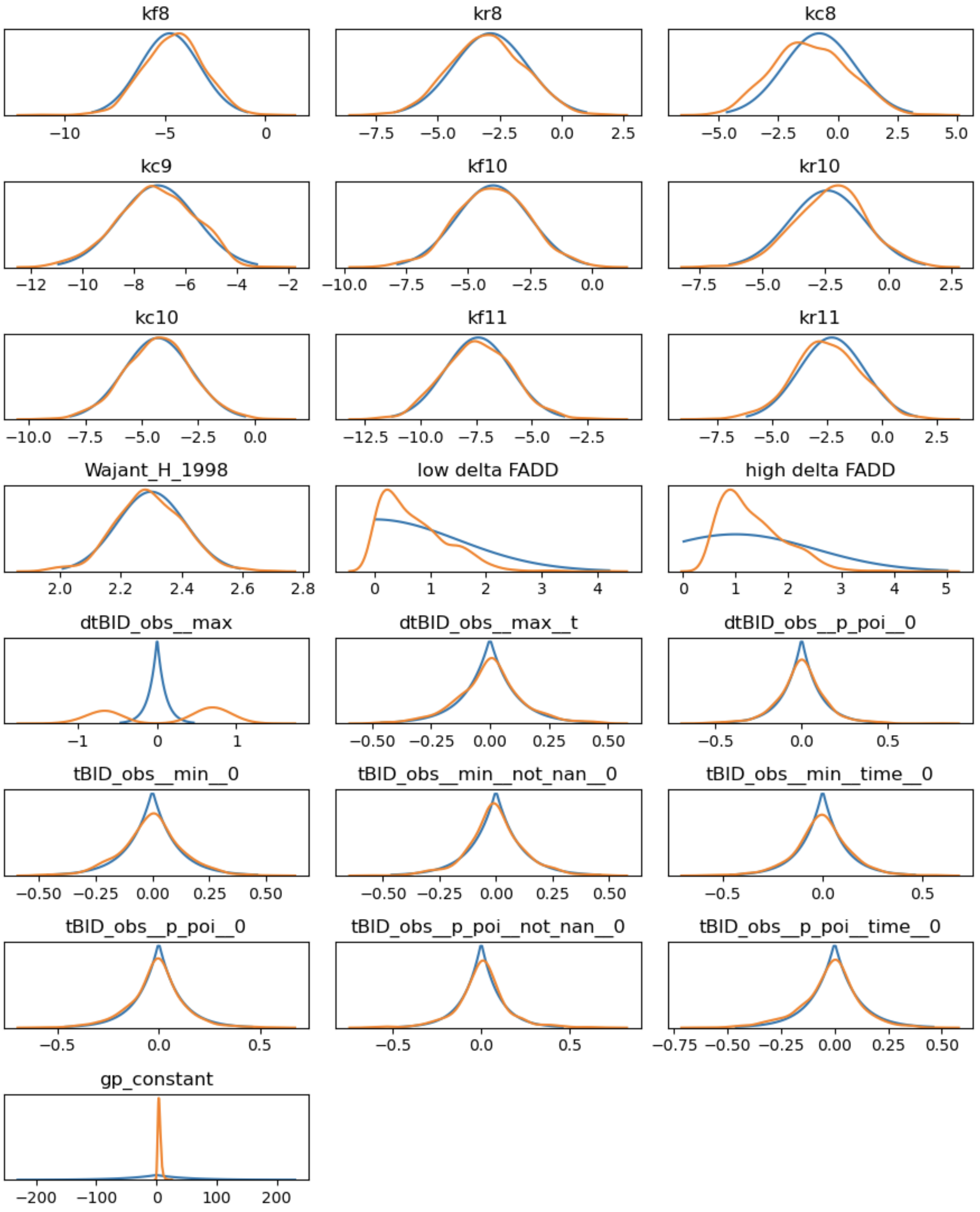

**Supplemental Figure 20B: Model parameters calibrated to published fractional cell death data.** Parameters for aEARM were calibrated to published fractional cell death measurements by Wajant et al. (see *Methods*)[31]. Prior (blue) and posterior (orange) distributions $\log_{10}$ of the value of the aEARM parameter are shown. Prior and posterior distributions of the value of measurement model coefficients are also shown (these are not log-scale).

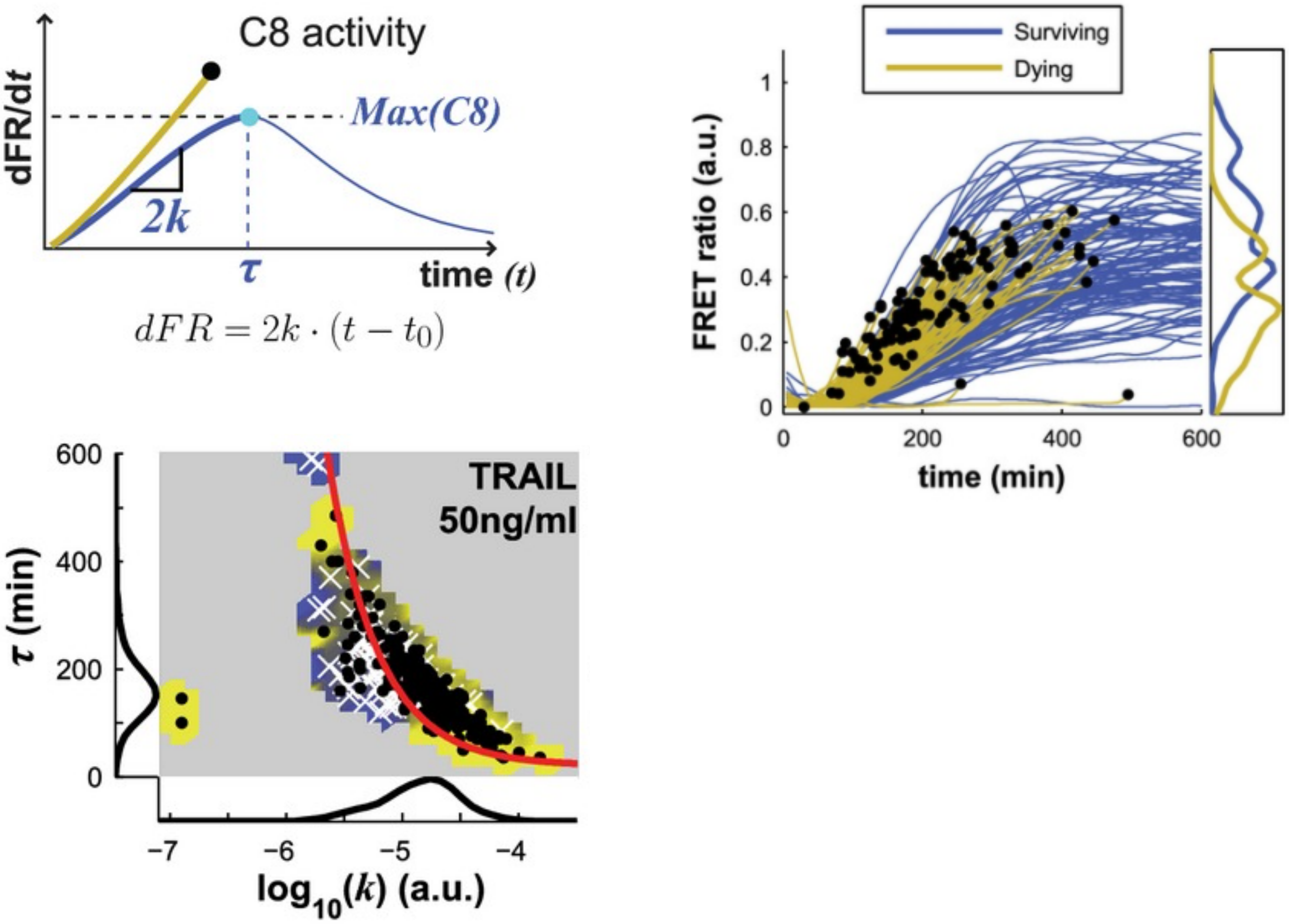

**Supplemental Figure 22: Reprinted from Roux et al.[19] caspase activity as a predictor of apoptosis in HeLa cells.**

A. The dynamics of Caspase activity, and by proxy rate of change in cleaved caspase substrate (e.g., Bid) concentrations, as indicated by a fluorescent indicator, served as features for predicting apoptosis (yellow) and survival (blue). B. The range of FRET ratio trajectories of HeLa cells treated with 25ng/mL TRAIL. C. The features dynamics of Caspase activity: $\log_{10} k$ which is a measure of maximum caspase activity (x-axis) and the time at which caspase activity maximizes $\tau$ (y-axis) served as predictors of apoptosis and survival, as shown in the plot of surviving and apoptotic cells.
